# G-quadruplex structures act as a novel recognition motif for the meiosis-specific histone methyltransferase PRDM9

**DOI:** 10.64898/2026.07.31.742039

**Authors:** Mia K. Hartge, Nuria Pfennig Hernández, Tomáš Pavlík, Nikol Poláková, Esther Schönauer, Julia Huber, Yasmin Striedner, Theresa Mair, Irene Tiemann-Boege, Matthias H. Weissensteiner, Johann Brandstetter, Angela Risch, Monika Cechova, Angelika Lahnsteiner

**Affiliations:** Division of Cancer (Epi-) Genetics, Department of Biosciences and Medical Biology, University of Salzburg, 5020 Salzburg, Austria; Center for Tumor Biology and Immunology (CTBI), University of Salzburg, 5020 Salzburg, Austria; Faculty of Informatics, Masaryk University, 60200 Brno, Czech Republic; Division of Structural Biology, Department of Biosciences and Medical Biology, University of Salzburg, 5020 Salzburg, Austria; Cancer Cluster Salzburg, 5020 Salzburg, Austria; Institute of Biophysics, Johannes Kepler University, 4040 Linz, Austria; Institute of Avian Research, 26386 Wilhelmshaven, Germany

**Keywords:** PRDM9, meiotic recombination, recombination hotspots, G-quadruplex (G4), DNA secondary structure, zinc-finger protein, chromatin accessibility, double-strand breaks (DSBs), promoter regions, hotspot specification

## Abstract

The histone methyltransferase PR domain containing protein 9 (PRDM9) is a key determinant of meiotic recombination in humans. It deposits activating histone marks thereby promoting recruitment of the meiotic recombination machinery. It recognizes DNA through a repetitive zinc-finger array that binds specific sequence motifs whose complementary G-rich strands can form DNA secondary structures, particularly G-quadruplexes (G4s). These may present an additional binding platform for PRDM9 and contribute to the formation of a chromatin environment permissive for meiotic recombination.

We investigated the relationship between PRDM9 binding sites and G4 motifs using computational analyses of predicted and experimentally validated G4s and found that G4 motifs are among the most prevalent features at PRDM9 binding sites, with the strongest enrichment observed for highly stable G4s, largely independent of loop length. Using electrophoretic mobility shift assays, we further examined whether PRDM9 can bind short, single-stranded G4-forming oligonucleotides in addition to its canonical double-stranded DNA targets. PRDM9 directly bound folded G4 structures, and binding increased with G4 stability. This relationship was observed across different G4 motifs and following stabilization of the same G4 by increasing the potassium concentration or adding a G4-stabilizing ligand. PRDM9 also bound an artificial G4-forming sequence absent from the human genome, which was abolished when mutating the G4 motif to avoid structure formation.

Together, these results support a model in which stable G4 structures facilitate PRDM9 recruitment by creating and discrete increased local chromatin accessibility, thereby contributing to the initiation of meiotic recombination.

**Graphical abstract:** 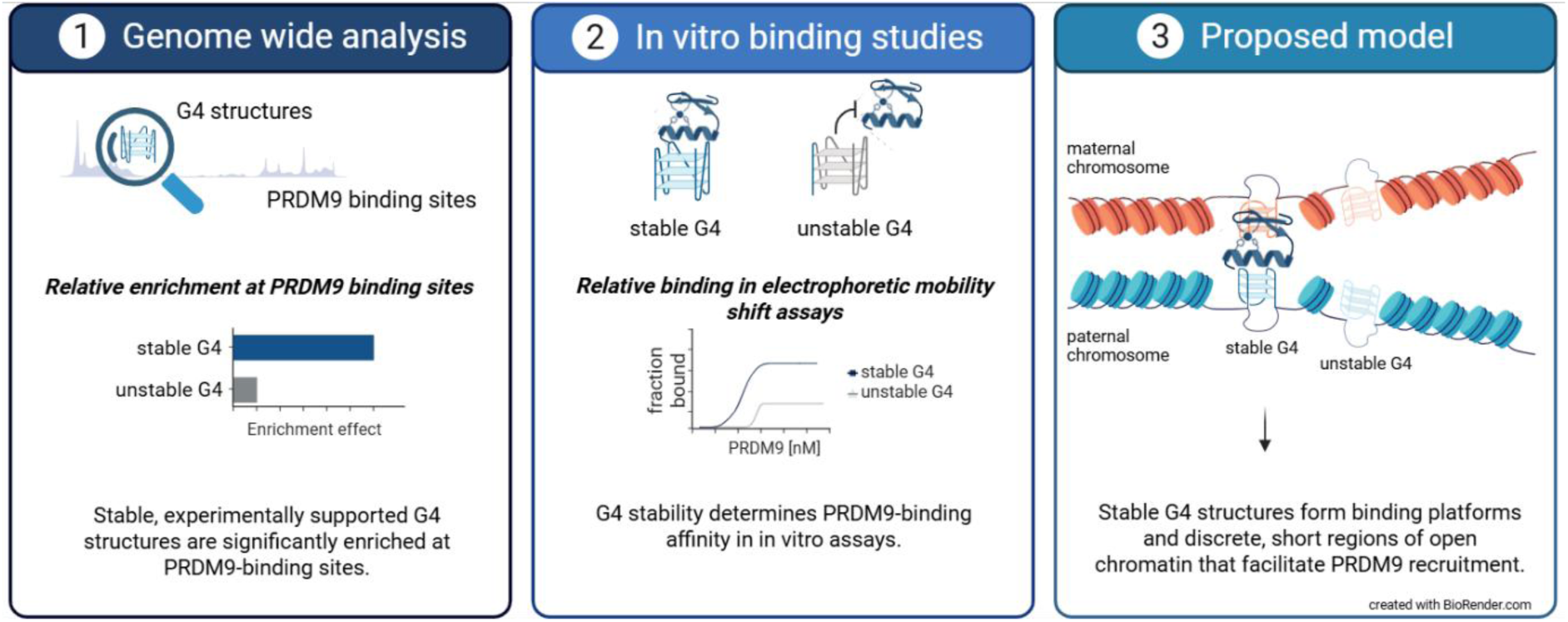

## Introduction

The timely introduction of hundreds of DNA double-strand breaks (DSBs) in narrow 1–2 kb regions called recombination hotspots, followed by their repair through the exchange of maternal and paternal DNA, is a fundamental process during mammalian gametogenesis [1–3]. Meiotic recombination allows the reshuffling of haplotype blocks—genomic regions containing variants that are usually inherited together—to generate new allelic combinations and to ensure the correct segregation of chromosomes during meiosis [3, 4]. In most mammals such as humans, great apes, mice and minke whales, but also in snakes, the zinc-finger (ZnF) protein PR-domain containing protein 9 (PRDM9) is the key determinant of meiotic recombination hotspots by binding to a degenerate sequence motif and placing the activating histone marks H3K4me3 and H3K36me3 to recruit the DSB machinery [5–11]. These histone marks are then recognized by ZCWPW1, which subsequently assists in homology search to allow the successful repair of the introduced DNA breaks [12].

PRDM9 consists of four different domains: a PR/SET domain with its histone methyltransferase activity responsible for placing H3K4me3 and H3K36me3 histone marks; a Kruppel-associated box domain (KRAB) and an SSX repression domain (SSXRD) thought to mediate protein-protein interactions, although their functions are not fully understood; and a C-terminal ZnF array which mediates DNA contact [13–15]. Although the overall architecture of PRDM9 is conserved across vertebrates, it is among the fastest evolving genes in the human genome [16]. This exceptionally rapid evolutionary rate is driven by its variability within the highly repetitive zinc-finger array [17]. Recent long-read sequencing approaches have revealed a far greater allelic complexity than previously thought. Analysis of 720 individuals across seven human populations identified 96 distinct PRDM9 alleles that differ in both the number and arrangement of individual ZnFs [18]. The two most prevalent alleles are PRDM9-A and PRDM9-C, with PRDM9-A predominating in European populations and PRDM9-C in African populations. A third allele, PRDM9-B, differs from PRDM9-A by a single nucleotide substitution within ZnF six and is mostly found in the Finnish population amongst Europeans, but also numerous rarer variants have been identified [18]. By far the greatest diversity has been found in the African population [18, 19], underscoring its rapid and ongoing evolution.

These allelic differences translate into distinct DNA recognition motifs: for example, PRDM9-A binds a 13-bp core motif (CCNCCNTNNCCNC) – also called “Myers” motif – active in at least 40 % of human recombination hotspots, while PRDM9-C recognizes a longer motif (CCNCNNTNNNCNTNNC), reflecting the presence of an additional ZnF [8, 20]. Remarkably, even minor alterations within the PRDM9 zinc-finger array can change its motif specificity, thereby profoundly reshaping the meiotic recombination landscape by directing PRDM9 to different sets of recombination hotspots [18, 20].

The distribution of PRDM9 binding sites is highly dynamic, as the rapid evolution of the PRDM9 ZnF-array continuously generates novel alleles and, consequently, new binding motifs, while previously active sites become progressively less targeted [21, 22]. In addition, the predicted PRDM9 binding motifs alone cannot explain PRDM9’s genome-wide occupancy or the resulting DSB landscape, as only a subset of the predicted motifs is targeted and many PRDM9-bound hotspots lack a recognizable consensus sequence [2, 8, 9, 20]. These observations suggest that additional, sequence-independent mechanisms contribute to the PRDM9-mediated selection of recombination hotspots.

Given that PRDM9 motifs share a high C content and a pronounced G/C-skew, folding of G-quadruplexes (G4s) represent a plausible candidate mechanism. G4s are non-canonical DNA structures with guanine-rich regions satisfying the motif G_≥3_N_x_G_≥3_N_x_G_≥3_N_x_G_≥3_, where N represents any base and x typically denotes a 1-7 bp DNA linker [23, 24]. However, structures with longer loops and more relaxed (non-canonical) patterns are also observed [25]. While stretches of consecutive guanines assemble stacked G-tetrads that form the G4 stem, the intervening nucleotides form connecting loops, giving rise to different topologies, including parallel, antiparallel, and hybrid structures [26]. G4 structures are stabilized by Hoogsteen hydrogen bonds and centrally coordinated monovalent cations, with K⁺ exerting the strongest stabilizing effect [27–29]. G4s are not randomly distributed over the whole genome but enriched in nucleosome-depleted regions and regulatory elements, such as promoters [30–33], enhancers [32, 34], and recombination hotspots [35, 36]. Although G4s were long considered merely as obstacles to polymerases with detrimental effects, it is now well established that they play important roles in transcription regulation [33, 37–41], DNA repair [42] and telomere maintenance [43, 44]. As G4s promote the formation of accessible chromatin [32], assist in recombination [35, 36, 45], are associated with DSBs as well as their repair [46], and recently have already been associated with PRDM9 binding in a recent preprint [47], they may facilitate the recruitment or activity of the meiotic DSB machinery. Hence, we hypothesize that G4 formation contributes to meiotic hotspot targeting by promoting PRDM9 binding and stabilizing nucleosome-free chromatin regions that permit efficient recruitment and activity of the DSB machinery especially at stable G4 structures.

Here, we report that G4 structures are enriched within human PRDM9 binding sites, largely independent of G4 loop length, while non-canonical G4 motifs exhibit substantially weaker enrichment than canonical G4s. We further show that PRDM9 binds both double-stranded DNA (dsDNA) fragments from two recombination hotspots on chromosome 16 and 21, as well as different single-stranded DNA (ssDNA) G4-motifs, with binding affinities strongly influenced by G4 stability. Consistent with these findings, stable G4s effectively compete with dsDNA fragments for PRDM9 binding, whereas less stable G4s display reduced competitive capacity.

Together, these results suggest that G4 structures—particularly highly stable G4s—are present within accessible chromatin prior to PRDM9 binding and may serve as permissive binding platforms that facilitate hotspot recognition during meiotic recombination.

## Methods

### PRDM9 ChIP-seq and BG4 CUT&Tag data analysis

PRDM9 chromatin immunoprecipitation and sequencing (ChIP–seq) data from HEK293 cells were obtained from the NCBI Gene Expression Omnibus (GEO) database with the accession no. GSE99407 [9, 48]. Raw sequencing reads were trimmed using default parameters with TrimGalore and then aligned to the human reference genome (hg38) using BWA-MEM [49, 50]. Peak calling was performed using MACS2 with a false discovery rate (FDR) threshold of q = 1 × 10⁻⁵, applying the broad peak setting on the Galaxy platform [51].

BG4 Cleavage under targets and tagmentation (CUT&Tag) data from HEK293 cells were obtained from [52–55] deposited at the SRA database with entry numbers SRR14300945, SRR14879748, SRR26244907, SRR26244908, SRR26244909, SRR26244910, SRR26244911, SRR26244912, SRR25010684 and SRR25010685 [56–65]. Adapter trimming was carried out using TrimGalore with default parameters [49]. Trimmed reads were then aligned to the human reference genome (hg38) using BWA-MEM [50]. Peak calling was performed using SEACR, with a threshold of 0.005 [66]. Alignments and peak calling steps of PRDM9 and BG4 datasets were performed on the Galaxy platform [51].

To generate a consensus set of G4 peaks, peak files from individual samples were intersected using *bedtools multiinter* in R [67, 68]. Only regions present in at least two samples were retained, while peaks detected in a single sample were excluded. To identify overlaps between PRDM9 binding sites and G4-forming regions, *bedtools intersect* was used, applying a window of ±25 bp around G4 peak regions [67].

Genomic annotation of G4 and PRDM9 peaks was performed using the R package ChIPseeker, which assigns peak regions to genomic features such as transcription start sites (TSS), exons, introns, and intergenic regions [69]. To visualize the density and distribution of peak occurrences around G4 motif centers, signals were stratified according to G4s formed on either the positive or negative strand, and enrichment profiles were generated using the R package ChIPseeker [69].

### Enrichment analysis

To identify genomic features associated with PRDM9 binding in HEK293 cells, we developed a reproducible pipeline characterizing the sequence-, epigenetic-, chromatin-, and recombination context of PRDM9-binding regions. We analyzed three groups of genomic regions: experimentally determined PRDM9-binding sites from ChIP-seq data, used as the foreground dataset (n = 60,580; length range: 195–43,663 bp; mean length: 1,153 bp), and two background (control) datasets—datasetA_matched and datasetB_global—sampled from regions of the hg38 that did not overlap PRDM9-binding sites. Both background datasets matched the foreground length distribution, while datasetA_matched was additionally matched for GC content and sequence entropy (***Supplementary File 1, Methods Section 1.1*** and ***Supplementary File 1, Fig. S1***).

We next used hg38 genomic annotations to derive features for each region in the PRDM9-binding, datasetA_matched, and datasetB_global sets for downstream enrichment analysis:

#### G-quadruplexes

G4 features were derived from genome-wide pqsfinder predictions [70]. Predicted loci were classified by experimental support, sequence architecture, and pqsfinder stability-score quartile. Support was defined by overlap with HEK293 CUT&Tag G4 regions, while sequence classes distinguished canonical short (1-3 nts)-, medium (4-6 nts)-, and long (7-12 nts)-loop motifs from non-canonical predictions. Composite category labels were then used to derive, for each region in the PRDM9-binding, datasetA_matched, and datasetB_global sets, the number of overlapping G4 loci and their coverage fraction. Full classification criteria, thresholds, and preprocessing details are provided in ***Supplementary File 1, Methods Section 1.2***.

#### DNA methylation

DNA methylation features were derived from reduced representation bisulfite sequencing (RRBS) of HEK293 cells from ENCODE experiment ENCSR794HFF [71]. Data from both replicate tracks were combined (3,903,640 calls, mean coverage of 39.9 ×; 27.21 % of sites had coverage <5×, 49.02 % exceeded >20 ×). Calls with coverage below 1 × or methylation values outside the 0–100% range were removed. Duplicate genomic coordinates from two replicate tracks were collapsed using the coverage-weighted mean methylation level.

The resulting dataset had a mean methylation level of 39.51 % and showed a bimodal distribution (***Supplementary File 1, Fig. S2***). Sites were subsequently assigned to four fixed methylation bins: Q1, 0–24 % (55.5 % of sites; mean methylation: 2.2 %); Q2, 25–49 % (3.7 %; mean: 35.0 %); Q3, 50–74 % (5.6 %; mean: 60.0 %); and Q4, 75–100 % (35.3 %; mean: 95.4 %). Q1 therefore represented low methylation and Q4 high methylation. For each region in the PRDM9-binding dataset, datasetA_matched, and datasetB_global, we calculated the fraction of bases covered by methylation calls in each category.

#### PRDM9-binding sequence motifs

Seventeen PRDM9-binding motifs (***Supplementary File 1, Fig. S3***) were obtained in MEME format [9]. Motif probability matrices were regularized using pseudocounts to avoid zero probabilities and converted to log-odds position weight matrices (PWMs) relative to a uniform nucleotide background (derived from 10,000 sites with E-values ≤ 1 × 10⁻⁵).

PRDM9-binding regions, datasetA_matched, and datasetB_global regions were scanned using both the forward and reverse-complement PWMs. Genomic start positions were counted only once, even when both strand-specific matrices produced a positive score. Ambiguous bases were permitted in the input sequences but could not contribute to a valid motif hit.

For each motif and genomic region, two features were calculated: the number of hits, defined as the number of genomic start positions with a positive PWM score, and the coverage fraction (covfrac), defined as the proportion of the region covered by the union of all motif-hit intervals. Overlapping hits were merged before coverage calculation to prevent double counting.

#### Chromatin accessibility

Chromatin accessibility features were derived from an ATAC-seq bigWig signal track for HEK293 cells (ENCODE accession ENCSR205FUM [72]). Each genomic region was divided into non-overlapping 25-bp windows, and the mean, minimum, maximum, standard deviation, 90^th^ percentile, and 95^th^ percentile of the bin-level signal was calculated. A high-accessibility threshold was estimated by randomly sampling 2 million genomic bins and calculating the 80th percentile of bins with positive signal values (of which there was 1.96 million, resulting in threshold equals to 0.253). For each region, we then calculated the fraction of bins exceeding this threshold. We additionally quantified local accessibility contrast as the mean signal within the region minus the mean signal across the adjacent 5-kb upstream and downstream flanks.

#### Recombination hotspots

Features describing meiotic recombination or double-strand-break hotspots were obtained from GEO database under accession no. GSE59836 [2, 73]. Preprocessing was done by deduplicating and merging overlapping intervals (reducing the dataset from 37,481 to 37,458 regions with mean length 1,498 bp). Hotspot centers were defined as interval midpoints. For each region in the PRDM9-binding dataset, datasetA_matched, and datasetB_global, proximity to the nearest hotspot was transformed into a proximity score bounded between 0 and 1, with higher values indicating greater proximity, so that positive enrichment consistently reflected closer association (e.g., proximity of 1 corresponds to overlap, 0.5 to 10 kb, and 0.1 to 90 kb):

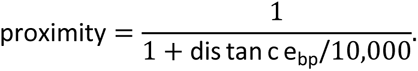

For each region, two features were derived using this proximity score: edge-to-edge proximity (edge of region to edge of nearest recombination hotspot) and center-to-center proximity (center of the region to the center of nearest recombination hotspot).

#### Histone modifications

We quantified histone modification enrichment using seven HEK293 ChIP-seq peak sets: H3K4me3 and H3K36me3 tracks from ENCODE with the accession no. ENCSR000DTU and entries ENCFF498ERO, ENCFF127YXW, ENCFF224LEB within [74], and matched experiments performed with and without PRDM9-B-HA transfection obtained from GEO database under accession no. GSE99407 [9, 48]. For each peak set, we removed exact duplicate intervals. For every PRMD9-binding, datasetA_matched, and datasetB_global region and histone track, we counted the number of overlapping peaks (hits) and computed the fraction of the window covered by their union (covfrac), defined as the total number of base pairs overlapped by at least one peak divided by the region length.

#### Analytical frameworks

Three separate enrichment analyses were calculated. In the global feature analysis (Figure 2A; Supplementary File 1, Fig. S4), each genomic feature was compared between PRDM9-binding regions (n = 60,580) and the datasetA_matched and datasetB_global backgrounds (each n = 181,740). Enrichment was evaluated using standardized mean differences, Welch’s t-tests, and a block-wise Monte Carlo procedure, with significance defined conservatively from both analytical and empirical false discovery rates. Full details are provided in ***Supplementary File 1, Methods Section 1.3***.

**Figure 1.**
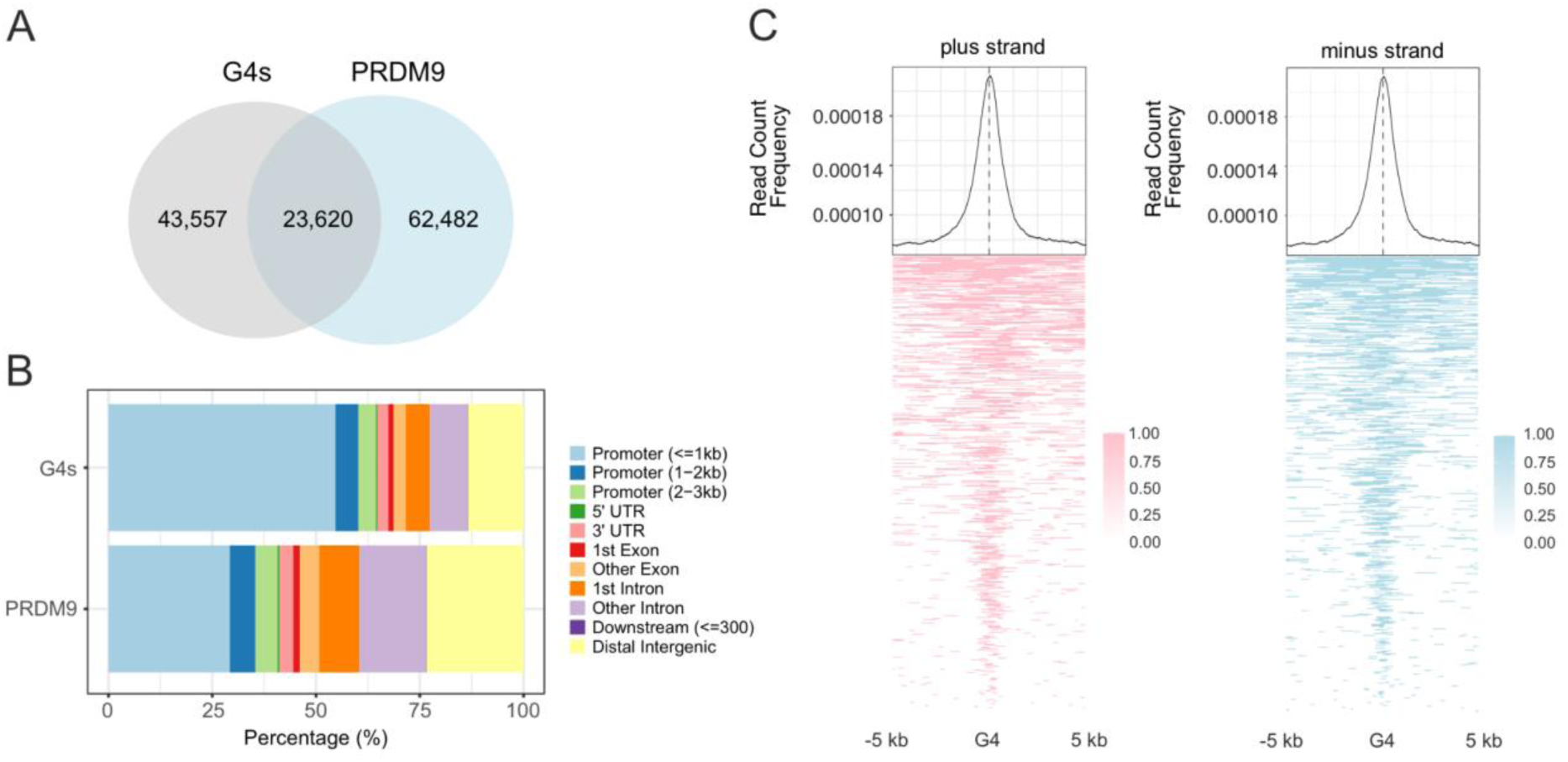
Overlap of G4 structures with PRDM9 binding sites. **A.** Venn diagram showing the number of unique and shared peaks identified from BG4 CUT&Tag and PRDM9 ChIP-seq data from HEK293 cells. **B.** illustrates the overlap of G4 forming sites and PRDM9 binding in different genomic features. **C.** Read count frequency plots and heatmaps at ± 5 kb of the G4 motif center showed clustering of PRDM9 binding at the center of the G4 motif without a difference in the strand orientation.

**Figure 2.**
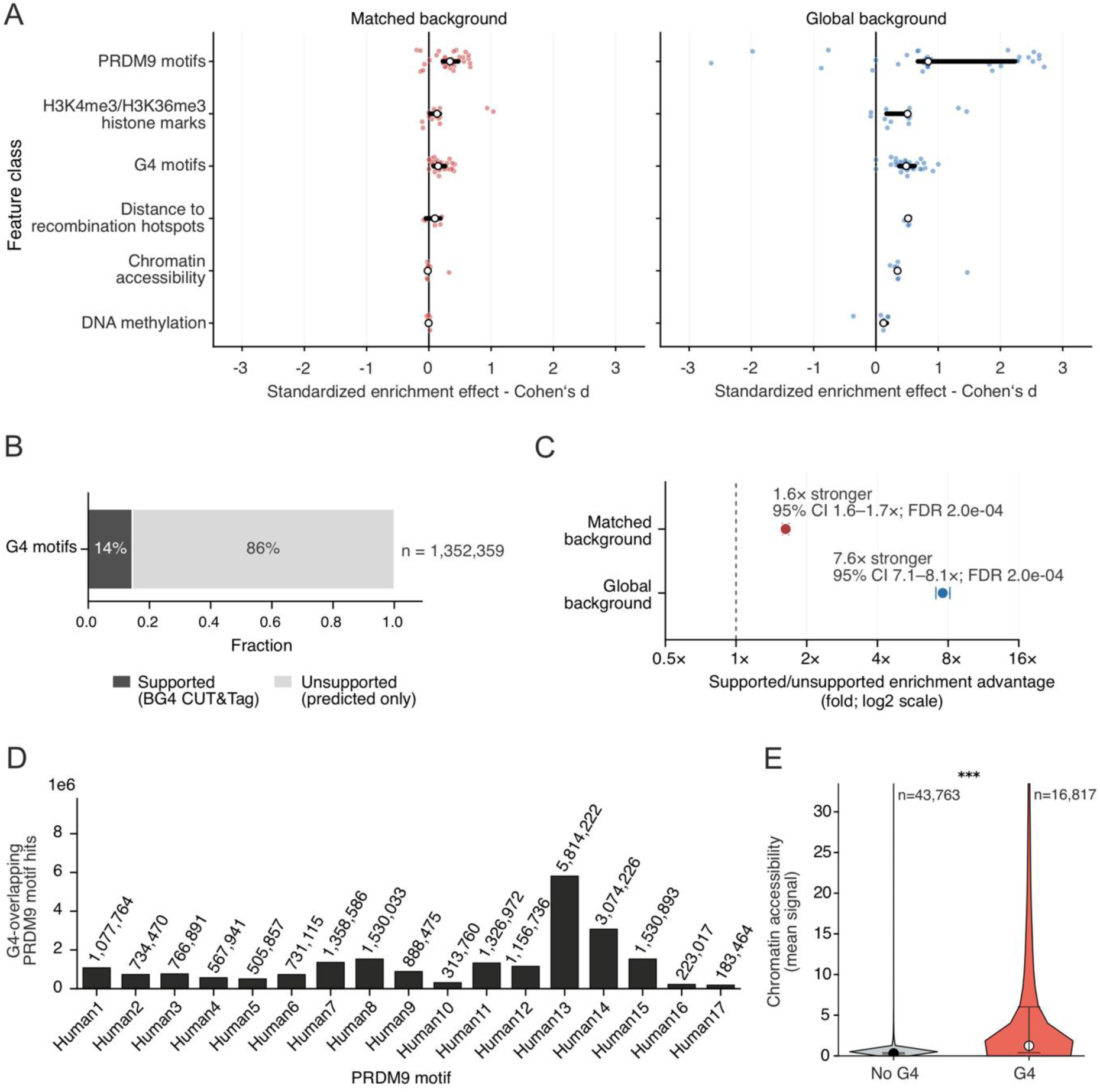
Enrichment analysis confirms significant overlap of PRDM9 binding sites with G4 structures. **A.** displays the enrichment of different features at PRDM9 binding windows using the matched background set (left panel) or the global dataset (right panel) as comparison. **B.** shows that only 14 % of predicted G4 motifs used in the analysis were validated by CUT&Tag data. **C.** Experimentally supported G4s were more strongly associated with PRDM9-binding regions and showed greater predictive value than unsupported (computationally predicted) G4 motifs. **D.** shows the overlap of G4 motifs with different classes of PRDM9 motifs. Position weight matrices for motif classes can be found in Supplementary File 1, Fig. S3. **E.** Median chromatin accessibility was 4.14-fold higher at PRDM9-binding sites overlapping experimentally supported G4s than at sites without G4s. Statistical significance was assessed using a one-tailed Mann–Whitney U test; *** p<0.001

Grouped G4 enrichment was used to identify the G4 subtypes most strongly associated with PRDM9 binding (Figure 3). Sequence predictions were stratified by sequence class, experimental support, and score quartile, and category-specific coverage fractions were compared between PRDM9-binding regions and both background datasets using the statistical framework defined in ***Supplementary File 1, Methods Section 1.4***.

**Figure 3.**
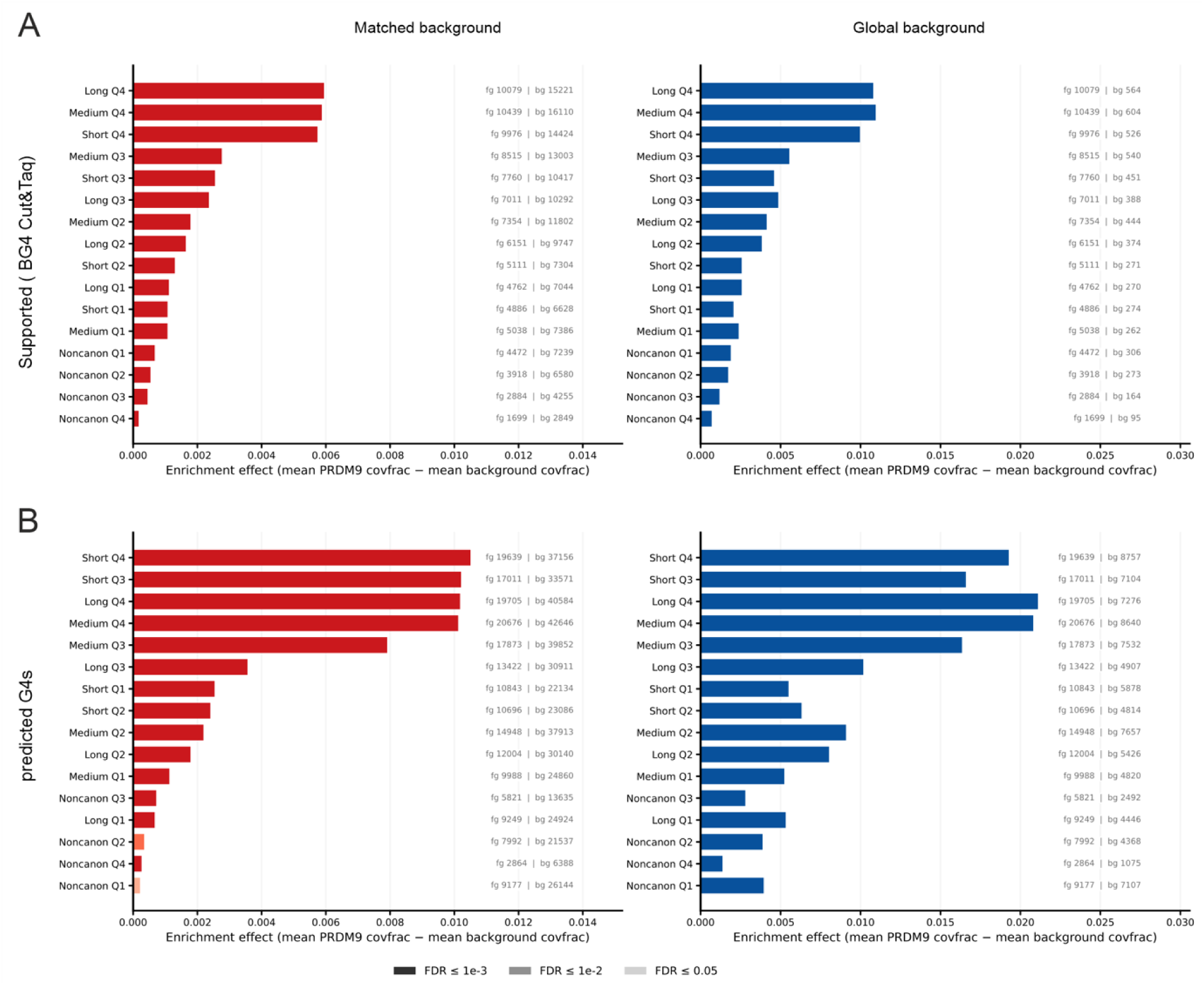
Analysis of G4 motif enrichment in PRDM9 binding sites across different motif classes. **A.** Enrichment of supported (experimentally validated) G4s stratified by pqsfinder scores (Q4 representing the most stable and Q1 the least stable structures) and by loop lengths. Enrichment was assessed relative to PRDM9-unbound genomic windows using either a matched background (left), controlled for length, GC content, and sequence entropy, or a global background (right), matched only for length distribution. **B.** Corresponding enrichment analysis for all in silico–predicted G4s, compared against the same matched (left) and global (right) background datasets.

Support-normalized G4 enrichment was used to test whether experimentally supported G4s were more strongly associated with PRDM9-binding regions than unsupported sequence predictions (Figure 2C–D). Predictions were classified by experimental support, sequence class, pqsfinder-score quartile, and length quartile. PRDM9-binding regions were compared separately with datasetA_matched and datasetB_global using support-specific rate ratios and a stratified odds-ratio. The framework is described in ***Supplementary File 1, Methods Section 1.5***.

Further details for regarding the analysis of chromatin accessibility and motif enrichment at G4 regions can be found in ***Supplementary File 1, Methods Section 1.6*** and ***1.7*.**

### Preparation of biotinylated DNA fragments for EMSAs

#### PCR fragments

Biotinylated PCR fragments were generated of regions covering the centers of recombination hotspot I (HSI) located in intron 2 of the *PCP4* gene (chr21: 39,905,075–39,908,853; hg38) [75–77], and recombination hotspot II (HSII) located in intron 2 of the *RBFOX1* gene (chr16: 6,309,072–6,312,440; hg38) [75, 78]. These regions were amplified using 1× GC buffer, 200 µM dNTPs, and 500 nM of each forward and reverse primer (with the forward primer carrying a 5′ biotin label), together with 0.1 U Phusion HSII polymerase (Thermo Scientific, Austria) and 10 ng genomic DNA isolated from HEK293 cells. PCR amplicons were purified using the ReliaPrep DNA clean-up and concentration kit (Promega, Austria) and validated on 1.5 % agarose gels.

#### Biotinylated oligonucleotides

Short single-stranded DNA oligonucleotides were ordered with a 5′ biotin label (Sigma-Aldrich, Austria) and reconstituted in 10 mM Tris-buffer pH 7.5 including 0 or 150 mM KCl. The unspecific dsDNA fragment was prepared by mixing complementary oligonucleotides, heating the mixture to 95 °C for 5 min, and then allowing it to cool slowly to room temperature.

All PCR primer and oligonucleotide sequences are listed in the ***Supplementary File 1, Methods Section 1.8*.**

### PRDM9 cloning, expression and extraction

The coding sequence of exons 1-9 of human PRDM9-A was amplified from cDNA generated from a testicular-sperm-extraction (TESE) and of exon 10 (encoding the zinc finger of the human PRDM9-A allele) was amplified from gDNA extracted from blood. The two fragments were fused via PCR and cloned into the empty pOPIN-M vector backbone containing an eYFP reporter generating the final expression construct. Further details can be found in ***Supplementary File 1, Section 1.9*.** The vector map is included as ***Supplementary File 2***.

To overexpress PRDM9-eYFP in HEK293 cells, 0.5 × 10⁶ cells were seeded in 6-well plates in RPMI + 10 % fetal bovine serum without antibiotics 24 h prior to transfection. A total of 1 µg of plasmid DNA was transfected using the Mirus TransIT-LT1 (Mirus Bio, USA) transfection reagent according to the manufacturer’s guidelines, followed by 24 h incubation at 37 °C and 5 % CO_2_.

Cells were harvested by scraping them from the surface, transferred to a 2 ml tube, followed by centrifugation for 5 min at 300 × g. The cells were then washed twice with ice-cold 1 × PBS and centrifuged for 5 min at 300 × g. The pellet was resuspended in 150 µl of EMSA extraction buffer (50 mM Tris-HCl, pH 7.5; 1 % Triton X-100; 1 mM EDTA; 140 mM NaCl; 500 µM ZnCl₂; 0.1 % sodium deoxycholate; and 1 × protease inhibitor cocktail). After incubation for 20 min on ice, the samples were centrifuged at 16,000 × g for 5 min at 4 °C. The supernatant was transferred to a new tube, and the total protein concentration was determined using the Bradford assay (Bio-Rad, Austria). 40 µg of whole cell protein were loaded onto an 8 % SDS-PAGE without heating to 95°C to preserve the eYFP folding, followed by fluorescence detection of eYFP on an Odyssey Fc imaging system (Li-Cor Biosciences, Germany) to assess expression efficiency.

PRDM9 concentration was determined by measuring absorbance at 514 nm using a Tecan Spark plate reader (Tecan, Austria) in a 384-well format. The extinction coefficient of eYFP was taken as 79,000 M⁻¹ cm⁻¹ according to the FPbase database [79, 80]. Concentrations were calculated using the Lambert–Beer equation:

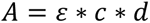

where *A* is the measured absorbance, *ε* the molar extinction coefficient in M⁻¹ cm⁻¹, *c* the protein concentration in mol/L, and *d* the path length in cm.

### RT-PCR to detect PRDM9 expression

Total RNA isolation was carried out using the innuPREP RNA Mini Kit 2.0 and the innuPREP DNase I Digest Kit (IST Innuscreen, Germany) according to the manufacturer’s instructions. cDNA was prepared using the iScript™ cDNA Synthesis Kit (Bio-Rad, Austria) with 500 ng input RNA and RT-PCR was carried out using the CFX Opus 384 Real-Time PCR system (Bio-Rad, Austria). In brief, primers for PRDM9 were designed to maximize the specificity for the gene in comparison to its homolog PRDM7. The forward primer (PRDM9-F-GTTTGA<u>A</u>AGAATTGTCA<u>A</u>GAAC<u>AG</u>; base differences to PRDM7 underlined) includes four PRDM9 specific positions and the reverse primer (PRDM9-R-CTGACACATCTCACAATAGAGG) was designed spanning exon 6 and 7. As reference gene GAPDH was used (GAPDH-F-GGGTGTGAACCATGAGAAGTAT and GAPDH-R-AGTAGAGGCAGGGATGATGT). The reaction mix was assembled in a final volume of 10 µl using the Blue S’Green qPCR Mix (Biozym, Germany) in a final 1 × concentration, 5 ng cDNA and the forward and reverse primers each to a final concentration of 0.25 µM. The following cycling conditions were used: 2 min of initial heating at 95 °C followed by 45 cycles of amplification with 5 sec at 95 °C, 10 sec at 66 °C, 10 sec at 65 °C including a SYBR Green plate read. Melting curve analysis was performed ranging from 65-95 °C with a temperature increase of 0.5 °C per 5 sec and a plate read after each increment. Gene expression changes were determined using the ΔΔCq method. All reactions were performed in at least triplicates.

### Electrophoretic Mobility Shift Assay

#### Binding reaction

EMSA binding reactions were adapted from previously reported protocols [81, 82]. 20 nM biotinylated dsDNA or ssDNA fragments were incubated with increasing amounts of whole cell extract reflecting 0-82 nM PRDM9 for dsDNA EMSAs and 0-620 nM PRDM9 for ssDNA G4 motifs in a total volume of 23 µl containing 1 × TKZN buffer (10 mM Tris–HCl, pH 7.5, 50 mM KCl, 0.05 % NP-40, and 50 µM ZnCl₂) and 0.3 % Sarcosyl. To enhance G4 stability, the final KCl concentration was adjusted to 150 mM while maintaining constant concentrations of all other buffer components. For competition assays, a constant protein concentration of 82 nM and 20 nM biotinylated (“hot”) DNA were incubated with increasing concentrations up to a 100-fold excess (2,000 nM) of unbiotinylated (“cold”) competitor DNA in a total reaction volume of 23 µl. Reaction mixtures were incubated at room temperature for 30 min and subsequently mixed with 5 µl of 5 × EMSA loading dye (1 × TKZN buffer, 15 % glycerol, and Orange G) prior to loading onto the polyacrylamide (PAA) gel.

Poly(dI-dC) was not included in the reactions because we found that it produced highly smeared EMSA signals likely due to nonspecific interactions with G4 motifs. Instead, we included several negative controls, including EMSAs performed with whole cell extracts lacking PRDM9 expression and EMSAs using a short unspecific dsDNA fragment.

#### Electrophoresis

A 6 % PAA gel was pre-electrophoresed in 0.5 × TBE pH 8 buffer at 100 V for 30 min to remove any unpolymerized acrylamide and to equilibrate the gel in the running buffer. The samples were then loaded, and electrophoresis was performed at 100 V for 90 min.

#### Blotting onto a nylon membrane

Blotting was performed using an Amersham Hybond-N+ nylon membrane (VWR, Austria) in a tank blotting system (Bio-Rad, Austria). Transfer was carried out in 0.5× TBE buffer (pH 8.0) under cooling on ice at a constant current of 320 mA for 1 h.

#### Blocking, incubation and signal detection

The membrane was UV-crosslinked with the DNA-facing side exposed to a 245 nm UV source using a gel documentation/cutting system for 12 min. To block nonspecific binding sites, the membrane was incubated in 1 × blocking buffer (5 % BSA in 1 × TBS) for 15 min at room temperature, followed by three washes with 1 × TBS for 5 min each.

Membranes were incubated with a 1:15,000 dilution of streptavidin–horseradish peroxidase (HRP) conjugate (stock concentration: 1.25 mg/ml; Thermo Fisher Scientific, Austria) in blocking buffer (1 × TBS containing 5 % BSA) for 15 min at room temperature under gentle agitation. Unbound streptavidin–HRP was removed by four washes with 1 × TBS-T (1 × TBS and 0.1 % Tween-20), each performed for 5 min under gentle agitation.

For chemiluminescence detection, membranes were incubated with ECL Select substrate solution (Bio-Rad, Austria) mixed in a 1:1 ratio for 2 min in the dark. Signal detection was performed using a ChemiDoc™ MP imaging system (Bio-Rad, Austria) with auto optimal exposure times.

The fraction of DNA bound by protein was calculated by quantifying band intensities of free DNA and the shifted protein–DNA complex using ImageLab software (Bio-Rad, Austria). The fraction bound was determined using the following equation:

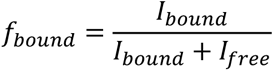

where *I_bound_*is the intensity of the shifted band and *I_free_*is the intensity of the unbound DNA band and plotted against the PRDM9 concentration determined via eYFP fluorescence measurements as described. In case that EMSA reactions resulted in several free DNA bands due to a heterogeneous mixture of B-like DNA and G4 structures, the intensities of the free DNA bands were calculated as the sum of all free DNA bands in one lane. Dissociation constants (K_d_) were obtained by fitting the resulting curves using a Hill slope with default parameters in GraphPad Prism (version 8; San Diego, CA, USA). Results from competition assays were fitted using a one-way decay curve with default parameters in GraphPad Prism (version 8; San Diego, CA, USA). EMSAs were performed at least in triplicates for each condition.

### Circular Dichroism Spectroscopy

Circular dichroism (CD) spectra were obtained from ssDNA oligonucleotides reconstituted in 50 mM Tris-HCl (pH 7.5) and either in 0 mM or 150 mM KCl in a final volume of 200 µL with an oligo concentration of 5 µM. CD spectra were recorded using the Chirascan™ Plus CD Spectrophotometer instrument (Applied Photophysics, Leatherhead, United Kingdom) equipped with a Peltier temperature-controlled cuvette holder and in-cuvette temperature sensor in a 0.5-mm path length quartz cuvette. Spectra were scanned from 340 to 220 nm with a step size of 0.2 nm and a bandwidth of 1 nm at 25°C. Four spectra per sample were averaged. After blank correction, the CD ellipticity θ in millidegrees was normalized using the equation

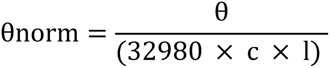

where c is the DNA concentration in mol/L and l is the path length in cm.

CD spectra were smoothed using locally weighted scatterplot smoothing (LOWESS) in GraphPad Prism (version 8; San Diego, CA, USA). Ellipticity values (θnorm, mdeg) were plotted against wavelength (nm), and a non-parametric LOWESS fit was applied to minimize spectral noise and highlight local trends without distorting the overall shape of the spectrum. The default smoothing factor was used. The resulting smoothed curves were used for visualization.

Melting curve analyses were performed at 260 nm from 24-92 °C in four replicates (temperature setting time: 200 s, step: 1 °C, scan time per point: 0.5 s). Data points were blank corrected, normalized to 24 °C, visualized and fitted with a sigmoidal curve using the default parameters in GraphPad Prism (version 8; San Diego, CA, USA).

### Statistical analysis

#### PRDM9 expression

Significant differences between untreated HEK293 cells and PRDM9-overexpression in qPCR experiments expressed as mean ± SD were assessed by using a student’s t-test.

#### EMSA

The fraction bound values were expressed as mean ± SD and significance was tested using a student’s t-test.

Statistical analysis for the bioinformatics analysis (enrichment analysis) was listed each in individual section above.

### Code availability

Custom scripts are available under https://github.com/angelikaLahn/PRDM9_G4_overlap and https://github.com/tommsko/PRDM9-enrichment development.

## Results

### G4 motifs are among the top enriched features at PRDM9 binding sites

Given PRDM9s preference for C-rich motifs able to form G4 structures on the complementary strand, and the evidence from recent studies that ZnF proteins such as MAZ, SP1 or the meiotic YY1 bind G4 structures [37, 38, 83–86], we aimed to understand whether PRDM9 can also bind to G4s. To do so, we analyzed ChIP-seq data from HEK293 cells overexpressing PRDM9-B [9] to assess PRDM9 binding sites and CUT&Tag datasets using the BG4 antibody specific for G4 structures to show G4 formation in HEK293 cells [52–55]. Quality controlled raw sequencing reads were aligned to the human reference genome assembly hg38, followed by peak calling as described in the methods section. In total, we identified 62,482 PRDM9 binding sites and 43,557 G4 peaks in HEK293 cells, of which 23,620 (37.8 %) regions were shared (***Figure 1A***). 54.7 % of all G4 peaks were located within 0-1 kb distance of the transcription start site (TSS) and a further 9.8 % within 1-3 kb of the TSS (***Figure 1B***). The largest fraction of PRDM9 binding sites accounting for 29.2 % of all peaks was identified within 0-1 kb of the TSS, followed by ∼25 % located in distal intergenic regions, indicating that both features are highly associated with promoter regions.

Although PRDM9 directs the recombination machinery away from promoters and promoter-like regulatory regions such as CpG islands and hence these regions are typically depleted of recombination events [9, 87, 88], the presence of PRDM9 binding sites within these regions suggests an intrinsic propensity of PRDM9 to recognize promoter-associated sequences or secondary structures such as G4s. When assessing the overlap between PRDM9 binding and G4s using the center of the G4 motif as the reference point, there was no difference between the strand orientation of G4 motifs detectable. (***Figure 1C***).

To analyze if G4s are enriched at PRDM9 binding sites in comparison to other genome-wide features such as PRDM9 binding motifs (***Supplementary File 1, Fig. S3***), H3K4me3 and H3K36me3 histone marks, DNA methylation, nucleosome accessibility and distance to recombination hotspots, we quantified their overlap with PRDM9 binding sites using both matched and global background models. Background datasets were derived from genomic windows without detected PRDM9 binding from ChIP-seq experiments and were either matched by length, GC content, and sequence entropy (matched background), or only by length distribution (global background) to the PRDM9-bound windows.

The sequence motif was the strongest enriched feature at PRDM9 binding windows, followed by histone marks and G4 motifs for both background sets (***Figure 2A***). The median absolute enrichment effect expressed as absolute Cohen’s d was slightly higher for G4s than for histone marks H3K4me3 and H3K36me3 for the matched background set, but not for the global background (***Supplementary File 1, Fig. S4***). Although the magnitude of enrichment varied depending on the background model used, the association between PRDM9 binding sites and G4 structures remained robust after controlling for genomic context. Recombination hotspot proximity also displayed consistent enrichment, whereas DNA methylation and chromatin accessibility showed comparatively less enrichment and least for comparisons with the matched background.

As G4 formation is strongly cell-type-dependent [32], we assessed whether G4 enrichment at PRDM9 binding sites was strongest for experimentally supported G4s or was also observed across all predicted motifs. Enrichment of all predicted motifs would suggest that the association is driven primarily by sequence composition rather than by a structure-dependent mechanism. Of the 1,352,359 predicted G4 motifs, approximately 14% overlapped with experimentally supported G4 sites in HEK293 cells that were already present prior to PRDM9 binding (***Figure 2B***). Among unsupported G4s, modest enrichment was observed at PRDM9 binding windows, with a 1.5-fold enrichment relative to the global background and a 9.4-fold enrichment relative to the sequence-matched background. In contrast, supported G4s showed substantially stronger enrichment, with 2.4-fold enrichment over the global background and 71-fold enrichment over the sequence-matched background (***Supplementary File 1, Fig. S5***). Thus, PRDM9 binding is associated predominantly with G4 structures that are experimentally detected in the genome, rather than with G-rich sequence motifs that merely have the potential to form G4s (***Figure 2C***). Among the 17 investigated PRDM9-binding motifs (***Supplementary File 1, Fig. S3***), motifs 13, 14, and 15 showed the greatest overlap, followed by motifs 7 and 8 (***Figure 2D***).

PRDM9, together with HELLS, have been identified as a pioneer factors capable of establishing accessible chromatin at recombination hotspots [89]. The relatively weak overall enrichment of pre-existing chromatin accessibility observed in our analysis is therefore consistent with accessibility being generated upon PRDM9–HELLS recruitment. However, G4 structures have also been associated with accessible chromatin [32]. We therefore examined whether the modest overall ATAC-seq signal identified in figure 2A concealed differences in specific features of the accessibility profile. Relative to the global genomic background, PRDM9-binding regions showed increased accessibility across all evaluated features, with the largest effect observed for the fraction of highly accessible bins (***Supplementary File 1, Fig. S6***). When compared with a sequence-matched background, however, most signal-intensity measures and local accessibility contrast showed little or no positive association. In contrast, the fraction of 25-bp bins exceeding the high-accessibility threshold remained elevated. Comparison of G4 occurrence and chromatin accessibility signals at genomic regions later bound by PRDM9 revealed that G4-containing regions were associated with significantly greater pre-existing chromatin accessibility. Because these data were obtained from HEK293 cells not transfected with exogenous PRDM9, this accessibility was already present independently of PRDM9 binding (***Figure 2F***). These findings suggest that PRDM9-binding regions contain a greater proportion of discrete, highly accessible subregions rather than showing uniformly increased accessibility across the entire binding region.

### PRDM9 binding sites are enriched with stable and canonical G4 motifs

Given that our results demonstrated the enrichment of G4 motifs within PRDM9 binding windows, particularly among experimentally validated G4s, and considering that G4 stability has been recognized as an important determinant of promoter function [33, 41], we sought to determine whether PRDM9 binding is similarly influenced by the stability of G4 structures. Therefore, we used G4 motifs predicted by the pqsfinder tool [70] with a minimum stability score of 47 and further classified them into canonical motifs satisfying the sequence motif G2-4N1-12G2-4N1-12G2-4N1-12G2-4 (with N being any base), as well as non-canonical sequence motifs with a more relaxed pattern. Canonical G4 motifs were further subclassified based on their loop length with 1-3 nts loops as “short”, 4-6 nts “medium” and 7-12 bp as “long”. Pqsfinder stability scores were divided into four quartiles (Q1–Q4), with Q4 representing the most stable and Q1 the least stable G4 motifs, corresponding to the lowest stability scores. In addition, BG4 CUT&Tag datasets were used to distinguish G4 motifs that were experimentally validated in HEK293 cells (supported G4s) from those identified solely through in silico prediction (unsupported G4s). To evaluate statistical significance and determine enrichment above background levels, both matched and global background datasets were included for comparison.

PRDM9 binding sites showed significant enrichment for highly stable (Q4) and experimentally validated G4 motifs relative to both the matched and global background sets (***Figure 3A***). Among canonical G4s, motifs with long and medium loop lengths exhibited the strongest enrichment, followed by motifs with short loops. In contrast, non-canonical G4s displayed even lower enrichment at PRDM9 binding sites than the least stable canonical G4 motifs (Q1).

When all in silico predicted G4 motifs were considered, a similar trend was observed, with more stable G4 motifs showing greater enrichment within PRDM9 binding regions (***Figure 3B***). However, G4 motifs with short loop lengths exhibited relatively stronger enrichment compared to the matched background set. As observed previously, greater variability was evident when the global background set was used for comparison. Consistent with the results obtained for experimentally supported G4s, non-canonical G4 motifs showed lower enrichment at PRDM9 binding sites than canonical G4 motifs when all computationally predicted G4s were analyzed.

In summary, stable G4 motifs, regardless of loop length, were significantly enriched at PRDM9 binding sites across both computational predictions and BG4 CUT&Tag–validated datasets. Notably, canonical G4 structures displayed a stronger overlap with PRDM9 binding sites compared to non-canonical G4s with more complex motifs including mismatches or bulges in the G4-stems.

### Electrophoretic mobility shift assays confirm binding of PRDM9 to single-stranded G4 motifs

To verify whether the observed genome-wide enrichment of G4 motifs at PRDM9 binding sites also results in a binding of PRDM9 to the secondary structure, or if this overlap was only sequence driven, we performed electrophoretic mobility shift assays (EMSAs). We overexpressed the most common European PRDM9-A full-length variant including the KRAB, SSRXD and PR/SET domain as well as the complete ZnF array fused to eYFP in HEK293 cells cloned into a pOPIN-M expression vector (***Figure 4A***). Following transient transfection and overnight expression, whole cell protein extracts were prepared in PRDM9 lysis buffer, as purification of PRDM9 remains challenging due to its repetitive structures, which promotes protein aggregation during extraction. We first confirmed successful PRDM9–eYFP expression at the transcript level by qPCR (***Figure 4B***). Transfected HEK293 cells showed a 5,745 ± 392-fold increase in PRDM9 transcript levels relative to untransfected cells. We then verified expression of the PRDM9–eYFP fusion protein by SDS–PAGE analysis. We included protein extracts from untransfected HEK293 cells and from cells transfected with an eYFP-only plasmid as controls. We detected successful expression of PRDM9–eYFP fusion protein and the eYFP-only protein in the corresponding transfected cells, whereas untransfected cells showed no signal in the Western blot.

**Figure 4.**
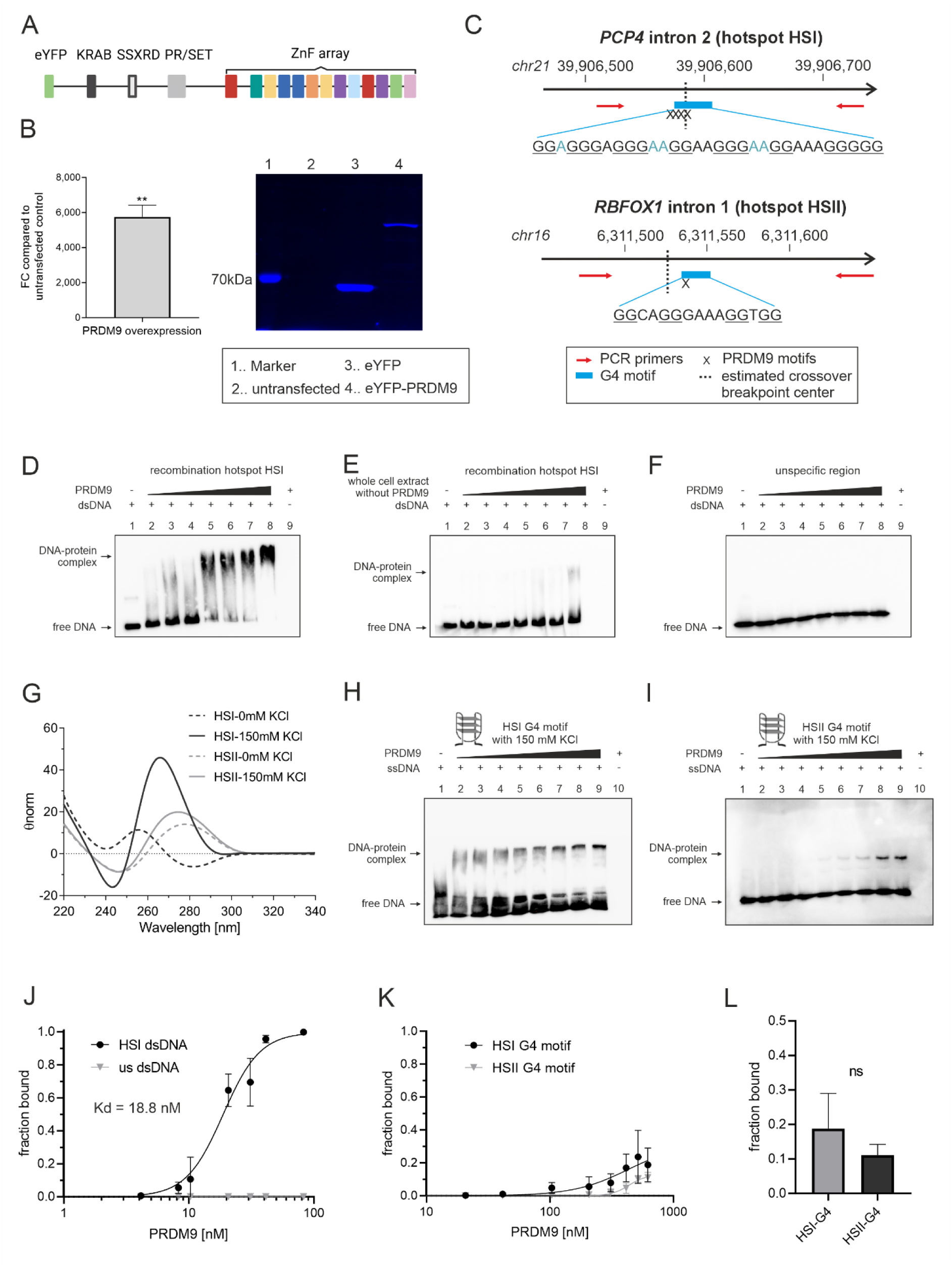
PRDM9 selectively binds recombination hotspots regions and single-stranded G4 structures. **A.** For EMSA experiments, the most common European PRDM9-A full-length variant fused to an eYFP reporter was transiently expressed in HEK293 cells. **B.** PRDM9 overexpression was verified using qPCR and SDS-Page followed by fluorescence detection using the eYFP tag. Lane 1: marker band, 2: untransfected HEK293 cells, 3: eYFP transfected HEK293 cells, 4: PRDM9-eYFP transfected HEK293 cells. **C.** shows the genomic region of recombination hotspot I (HSI) located in *PCP4* intron 2 and HSII located in *RBFOX1* intron 1. Both hotspot centers included a G4 motif with the HSI G4 motif being more stable (pqsfinder score 89) compared to the HSII G4 motif (pqsfinder score 29). Biotinylated PCR primers (red arrows) target 224 bp (HSI) and 178 bp (HSII) regions covering the G4 motif (blue rectangle) as well as the PRDM9 motifs (X) at the estimated crossover breakpoint center from sperm typing experiments. **D.-E.** EMSA experiments using the biotinylated PCR product from HSI with PRDM9 overexpressed whole cell extract (D) or the whole cell extract without PRDM9 (E) reveal a mobility shift indicative of specific PRDM9 binding to the target DNA sequence. **F.** No binding was detected when using a 48-bp double-stranded unspecific DNA fragment. **G.** CD spectra confirmed K^+^-dependent parallel G4 formation indicated by a negative peak at 245 nm and a positive peak at 260 nm of the HSI G4 motif (black solid line). The less stable HSII sequence showed a broad positive peak at approximately 270–280 nm, consistent with a heterogeneous mixture of B-DNA-like and G4-conformations. **H.-I.** EMSA analysis indicates specific interaction of PRDM9 with recombination hotspot HSI (H) containing a central single-stranded G4 motif overlapping with several PRDM9 motifs and recombination hotspot HSII (I) containing a G4 motif overlapping with a single Myers motif. **J.-K.** Binding curves for the double-stranded hotspot DNA fragment (J) and the single-stranded G4-forming oligonucleotides (K) were generated by plotting the fraction of bound DNA against the PRDM9 concentration and fitting the data using a Hill equation. An apparent dissociation constant (Kd) was determined only for the HSI dsDNA fragment, as the HSI-G4 and HSII-G4 ssDNA oligonucleotides did not reach a plateau under the used EMSA conditions. **L.** Fraction of HSI-G4 and HSII-G4 ssDNA bound at 620 nM PRDM9, the highest concentration tested in the EMSA experiments. Bars represent mean ± SD. Statistical comparison using an unpaired t-test showed no significant difference. ** *p*<0.01

Next, we aimed to determine whether PRDM9 binds to predicted PRDM9 motifs found at recombination hotspots. We selected two well-characterized recombination hotspots that were investigated using sperm-typing experiments and that harbor PRDM9-binding sites overlapping with G4 motifs near the double-strand break (DSB) center and the estimated crossover breakpoints. Recombination hotspot I (HSI) is located in intron 2 of the *PCP4* gene (chr21:39,905,075–39,908,853; hg38; ***Figure 4C* *top***) [75–77], and recombination hotspot II (HSII) is located in intron 2 of the *RBFOX1* gene (chr16:6,309,072–6,312,440; hg38; ***Figure 4C* *bottom***) [75, 78]. Target regions covering the hotspot centers including the PRDM9 and G4 motifs were PCR amplified using biotinylated primers, yielding fragments of 224 bp for HSI and 178 bp for HSII. EMSA reactions were prepared in EMSA binding buffer with constant DNA concentrations, while protein concentrations were incrementally increased across reactions. Each EMSA included DNA-only (lane 1) and protein-only (lane 9 and 10) controls to assess nonspecific interactions. Protein-DNA complexes were detected using a streptavidin–horseradish peroxidase conjugate which targets the biotinylated DNA fragments in a chemiluminescent reaction.

We detected strong PRDM9 binding to the biotinylated dsDNA fragment spanning the center of recombination hotspot HSI containing four PRDM9 motifs and a G4 motif with increasing protein concentrations resulting in higher signal intensities. Both control lanes (lane 1, DNA only; lane 9, protein only) did not show any unspecific signals (***Figure 4D***). Similar to HSI, also for HSII strong binding of PRDM9 to dsDNA was obtained, although we just used the highest protein concentration to show binding (***Supplementary File 1, Fig. S7***). To exclude the possibility that the observed binding originated from background in the whole cell extract, we repeated the EMSA using biotinylated HSI dsDNA incubated with extract from HEK293 cells lacking PRDM9 transfection. Only weak complex formation was observed under these conditions **(*Figure 4E*)**, indicating that the observed protein–DNA complex was predominantly formed by PRDM9. Binding specificity was further assessed by performing EMSAs with a nonspecific biotinylated dsDNA fragment, which did not yield detectable complex formation with PRDM9 **(*Figure 4F*)**.

Since our EMSAs showed selective binding of PRDM9 to dsDNA including the PRDM9 binding motifs originating from recombination hotspots, we aimed to understand if PRDM9 is able to recognize the single-stranded G4 motifs found in these two hotspots. Recombination hotspot HSI contains a G4 motif (chr21: 39,906,576–39,906,606; hg38) with a predicted pqsfinder stability score of 89 which overlaps with several PRDM9 motifs in the hotspot center, while the G4 motif in HSII is less stable with a score of only 29 (chr16: 6,311,536-6,311,558; hg38). These two motifs were then slightly extended by the flanking sequence in the hotspots to obtain a 50-bp single-stranded biotinylated fragment used for EMSA experiments. First, G4 formation of hotspot-derived G4 motifs was assessed via circular dichroism (CD) spectroscopy containing either 0 mM or 150 mM KCl in 50mM Tris-buffer pH 7.5. While the five-tetrad HSI G4 motif showed clear parallel G4 formation with a negative peak at 245 nm and a positive peak at 260nm in a K^+^-dependent manner **(*Figure 4G*)**. In contrast, the two-tetrad HSII G4 motif displayed a broad positive maximum at approximately 275–280 nm and a negative minimum near 245 nm. This suggests that the less stable HSII-G4 predominantly adopts a mixture of duplex- or hairpin-like conformation with the presence of a G4 structure.

At the initial KCl concentration of 50 mM in the EMSA binding buffer, the HSI G4 ssDNA produced a broad smear in the DNA-only lane (lane 1), suggesting the presence of a heterogeneous mixture of folded G4 structures and unfolded ssDNA ***(Supplementary File 1, Fig. S8***). Therefore, also the PRDM9-ssDNA complex formation showed additional bands near the slot, probably due to intermolecular G4 formation and aggregation of DNA-protein complexes. To overcome this, G4 stabilization was enhanced by increasing the KCl concentration to 150 mM in the EMSA binding buffer. Under these conditions, the DNA smear became less prominent, and a well-defined DNA–protein complex appeared with increasing PRDM9 concentrations, demonstrating that PRDM9 binds the single-stranded HSI G4-forming sequence (***Figure 4H***). We performed the same experiment with the less stable HSII sequence and observed substantially weaker PRDM9 binding (***Figure 4I***). Together, the stronger binding to HSI than to HSII supports the conclusion that the G4 stability influences its recognition by the PRDM9 zinc-finger domain.

Although EMSA experiments represent a simplified in vitro system that does not account for chromatin context, including histones or additional interacting factors like proteins stabilizing the complex or important during recombination initiation, they nonetheless enable comparison of the relative binding affinities of PRDM9 to different DNA fragments. This was achieved by estimating dissociation constants Kd from signal intensities, expressed as the fraction bound, calculated as the intensity of the shifted band divided by the sum of the intensities of the free and shifted DNA bands. The calculated Kd for PRDM9 to its specific binding region in HSI was calculated as 18.8 nM (95% CI: 15.9–22.4 nM, ***Figure 4J***), which was within the range of other reported values for PRDM9 [81, 82]. Because PRDM9 binding to the HSI-G4 and HSII-G4 ssDNA oligonucleotides did not reach a plateau within the tested concentration range, reliable Kd values could not be determined (***Figure 4K***). Therefore, we compared the fraction bound at the highest tested PRDM9 concentration (620 nM) and observed an approximately twofold higher fraction bound for HSI-G4 than for HSII-G4, although this difference did not reach statistical significance (***Figure 4L***).

### PRDM9 preferentially binds stable G4 structures

To determine more in detail whether the stability of a G4 structure influences PRDM9 binding, we selected two similar G4-forming sequences from opposite ends pqsfinder stability distribution—defined as quartile (Q) 1 and 4—with high and low pqsfinder-inferred stability, respectively. The Q4 motif originated from the most stable quartile and had a predicted stability score of 73, whereas the Q1 motif originated from the least stable quartile and had a score of 53 **(*Figure 5A*)**. Importantly, the motifs had comparable overall lengths—15 nts for Q4 and 16 nts for Q1—and both contained short loops, thereby limiting differences in general motif architecture. CD spectroscopy confirmed that both sequences formed K^+^-dependent G4 structures. We next compared their thermal stabilities by monitoring the CD signal at 260 nm during heating from 24 to 94 °C. The Q1-G4 displayed a melting transition with a melting temperature of 78 °C. By contrast, the Q4-G4 did not completely unfold within the measured temperature range, preventing reliable determination of its melting temperature and indicating substantially greater thermal stability **(*Figure 5B*)**. Thus, the experimentally observed difference in thermal stability was consistent with the pqsfinder predictions.

**Figure 5.**
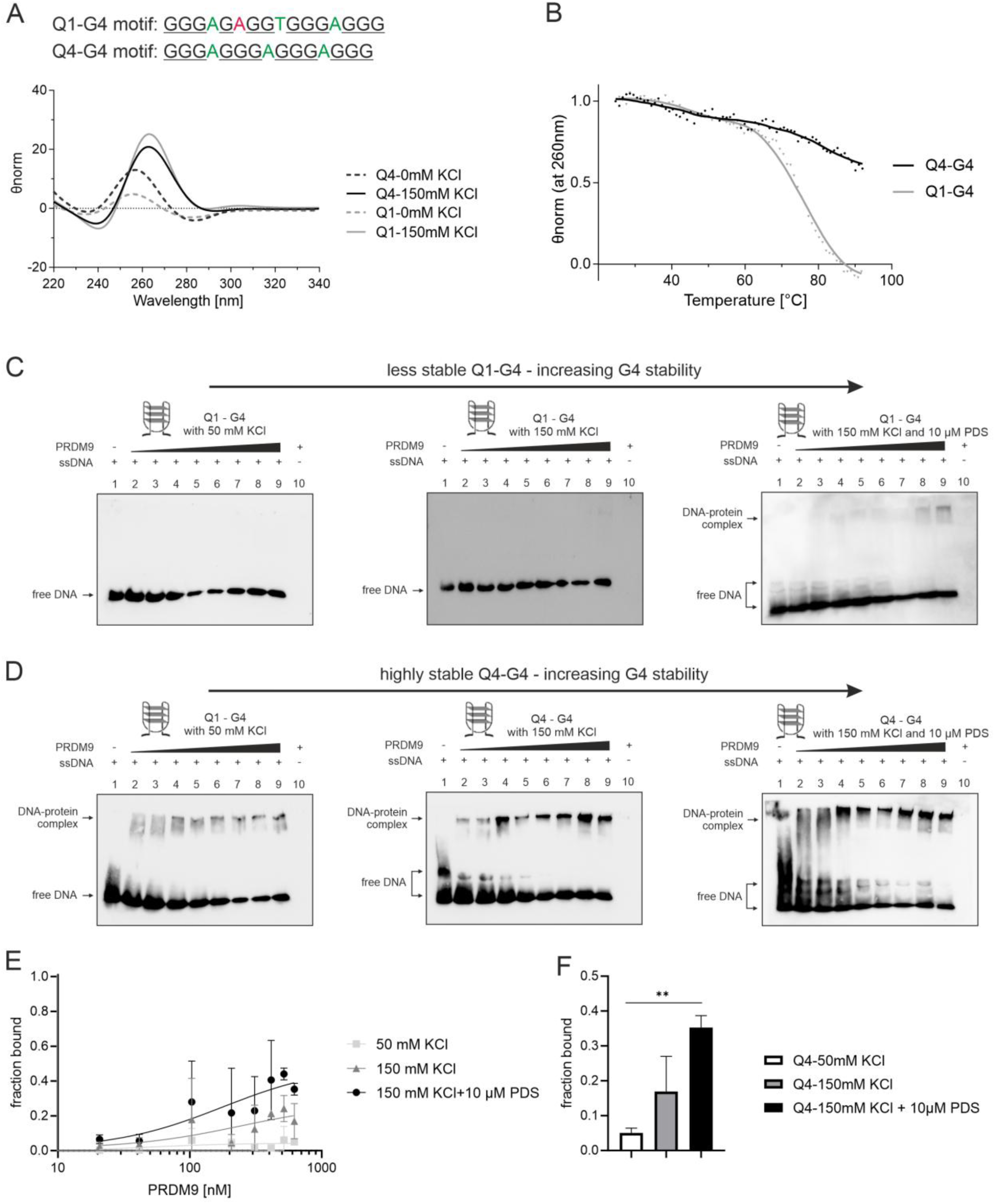
PRDM9 prefers stable G4 structures. **A.** depicts Q4 and Q1 G4 motifs used in EMSA experiments. Bases in green reflect G4-loops and in red a bulge in a G4 stem. CD spectra confirmed K^+^-dependent parallel G4 formation for both motifs. **B.** Melting curve analysis confirmed the predicted differences in G4 stability, with Q4 exhibiting a highly stable G4 structure. **C.-D.** EMSA experiments with PRDM9 using either the single-stranded Q1 (least stable, pqsfinder stability score= 53, C) or Q4 (most stable, pqsfinder stability score= 73, D) G4 motif for binding studies in EMSA binding buffer (left EMSA), EMSA buffer with 150 mM KCl (middle) or EMSA buffer with 150 mM KCl and 10 µM pyridostatin (PDS, right). PRDM9 binding was detected for the Q4 G4 motif under all three conditions, whereas the Q1 G4 motif showed only weak binding upon stabilization with PDS. **E.** Binding curves were obtained by estimating the fraction bound from EMSA blots using a Hill fit. **F.** Fraction of Q4-G4 ssDNA bound at the highest PRDM9 concentration tested (620 nM) under the indicated EMSA conditions. G4 stabilization with 150 mM KCl and 10 µM PDS significantly increased PRDM9 binding compared with 50 mM KCl. Bars represent the mean ± SD. Statistical significance was assessed using an unpaired t-test; **p<0.01.

We then tested whether this difference in G4 stability was associated with changes in PRDM9 binding affinities. We performed EMSAs under three conditions designed to progressively enhance G4 stability: 50 mM KCl, 150 mM KCl, and 150 mM KCl with 10 µM of the G4 stabilizing agent pyridostatin (PDS). PRDM9 binding to the Q1-G4 was undetectable at both 50 and 150 mM KCl, and only a small bound fraction was observed following PDS treatment (***Figure 5C***). In contrast, PRDM9 bound the more stable Q4-G4, and the extent of binding increased under conditions that promoted G4 stabilization (***Figure 5D***). Because PRDM9 binding to the Q4-G4 ssDNA oligonucleotide did not reach a clear plateau under any of the tested conditions, reliable Kd values could not be determined from the binding curves (***Figure 5E***). Instead, we compared the fraction of bound DNA at the highest PRDM9 concentration tested, 620 nM. The fraction bound was lowest in EMSA buffer containing 50 mM KCl, whereas stabilization of the G4 structure with 150 mM KCl and 10 µM PDS significantly increased the fraction of Q4-G4 ssDNA bound by PRDM9 (***Figure 5F***).

Together, these results establish an association between G4 stability and PRDM9 binding: the more stable Q4-G4 was preferentially bound by PRDM9, and further stabilization of this structure enhanced binding. These findings support a model in which the structural stability of a G4 motif contributes to its recognition by PRDM9.

### PRDM9 recognizes artificial G4 structures formed by a sequence not found in the human genome

To test whether PRDM9 binding depends on the general structural features and stability of a G4 motif rather than on its specific nucleotide context, we designed an artificial G4 motif with short loop lengths and a total length of 27 nts that cannot be found in the human genome (pqsfinder stability score = 64) and does not harbor a predicted PRDM9 binding motif. In addition, we mutated the G4 motif by exchanging four Gs from the stems against Ts (pqsfinder stability score= 0), to avoid G4 formation (***Figure 6A***).

**Figure 6.**
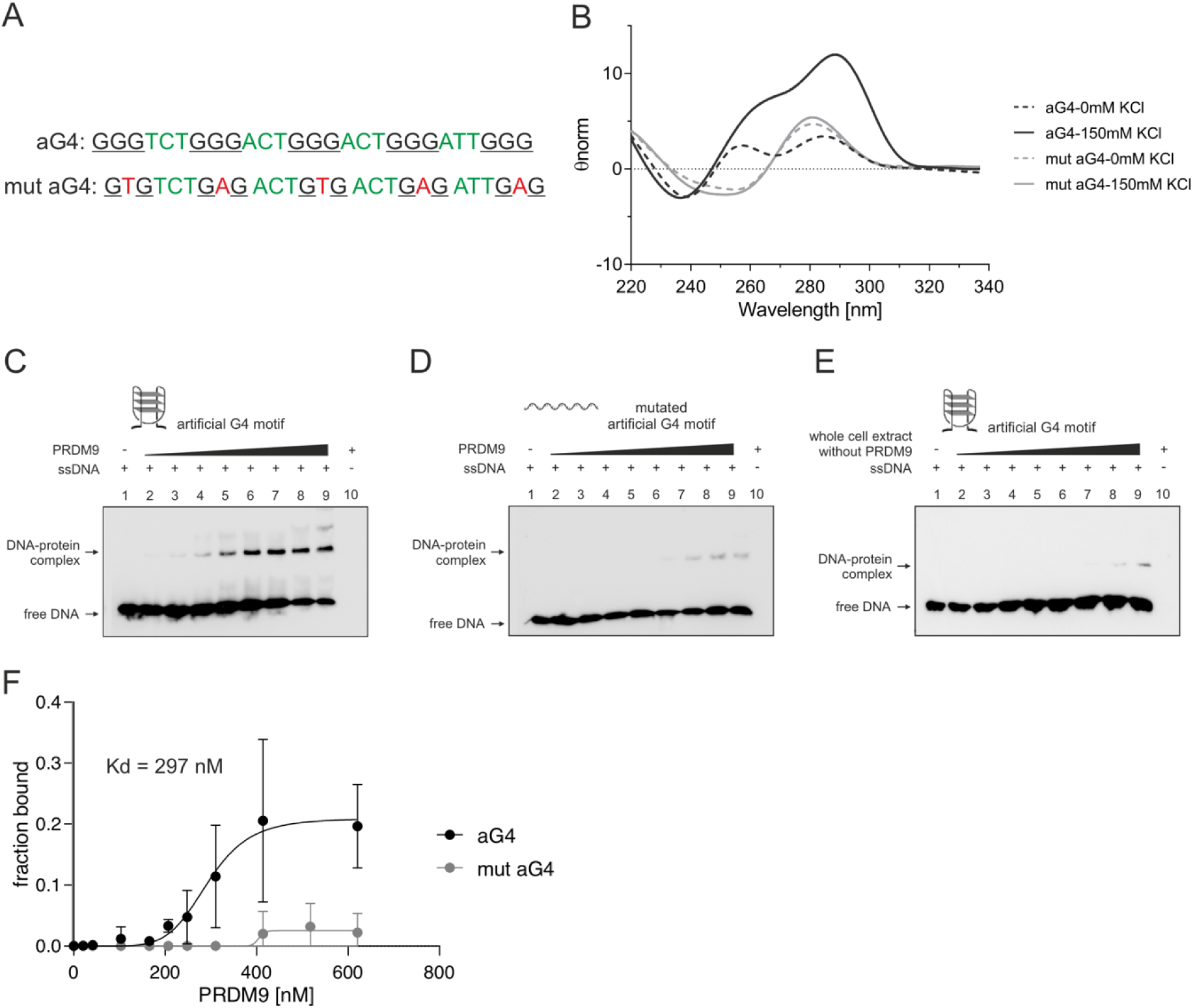
PRDM9 recognizes an artificial G4 structure. **A.** indicates the sequences used for EMSA experiments. Black underlined bases are the G4 stems, green represent loop bases and red bases indicate the introduced mutated bases to interrupt G4 stems. B. CD spectra confirmed K+-dependent formation of a hybrid G4 structure by the artificial G4 motif, whereas mutations introduced into the G4 stems abolished G4 formation and showed B-like structures without K+ dependence. **C.-D.** human PRDM9-A was able to recognize an artificial G4 structure not found in the human genome (A), while the mutated G4 motif is less bound (B). **E.** When using the whole cell extract without overexpressed PRDM9 only a slight background signal is visible. **F.** Binding curves fitted by a Hill slope resulted in a Kd value of 297 nM for the artificial G4, while no Kd value was obtained for the mutated structure.

CD spectroscopy confirmed that the artificial G4 sequence (aG4) adopts a K⁺-dependent hybrid G-quadruplex topology, whereas the mutated sequence (mut-aG4) showed no evidence of G4 formation (***Figure 6B***). In subsequent EMSAs, PRDM9 bound the folded aG4 oligonucleotide, whereas targeted disruption of G4 formation abolished detectable PRDM9 binding (***Figure 6C and D***). Whole cell extracts lacking PRDM9 expression showed only weak complex formation with aG4, supporting the specificity of the observed interaction (***Figure 6E***). The apparent Kd for aG4 was 297 nM, whereas no Kd could be determined for mut-aG4 because binding was not detectable (***Figure 6F***). Thus, the loss of G4 folding upon mutation was accompanied by the loss of PRDM9 binding. These results demonstrate that PRDM9 can recognize a G4 DNA secondary structure independently of its canonical sequence motif, even when the G4 is formed by an artificial sequence not found in the human genome.

### Folded G4 structures compete for PRDM9 binding with dsDNA from recombination hotspots

To assess the relative competition effect of G4s with different stabilities with its canonical dsDNA target, we performed EMSA competition assays using constant biotinylated HSI dsDNA (20 nM) and a constant concentration of PRDM9 (82 nM) in each reaction, while adding increasing amounts of an unbiotinylated (“cold”) HSI-derived single-stranded G4-forming oligonucleotide as competitor. Competitor concentrations ranged up to a 25-fold molar excess relative to the labeled probe **(*Figure 7A*)**. Although PRDM9 binding to the dsDNA target remained predominant, a reduction in complex formation was already observed at a 5-fold excess of the cold G4 competitor, indicating that the single-stranded G4 motif can effectively compete with the canonical dsDNA substrate for PRDM9 binding. In contrast, a less stable G4 competitor produced a slower decline in PRDM9 binding to the HSI dsDNA probe, consistent with its reduced competitive capacity **(*Figure 7B*)**. This difference was also reflected in the corresponding binding curves (***Figure 7C***).

**Figure 7.**
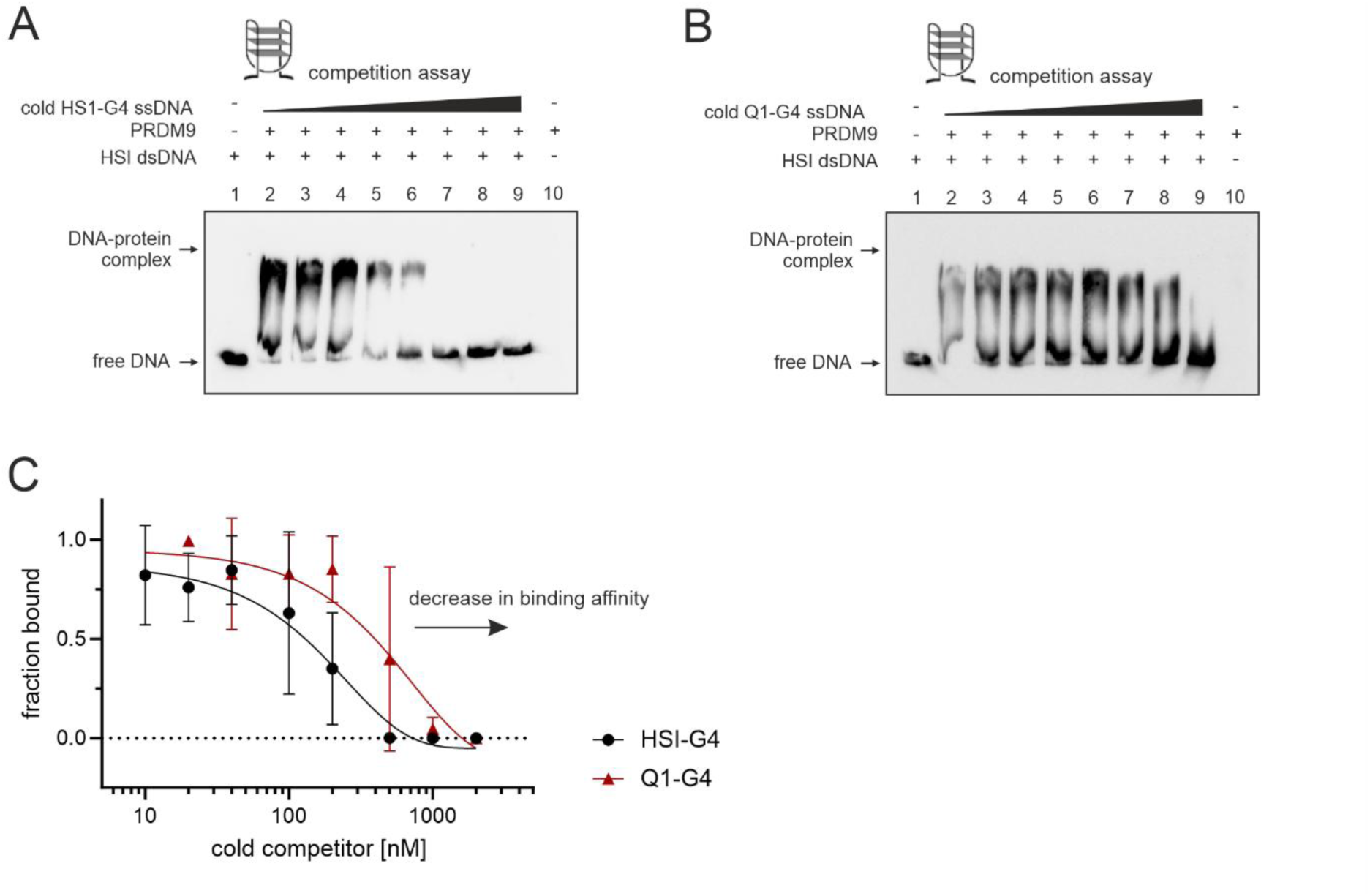
G4s can compete for binding with PRDM9 compared to dsDNA. **A.-B.** shows EMSA competition blots using 20 nM biotinylated HSI – dsDNA with increasing concentrations of HSI-G4 ssDNA (A, pqsfinder stability score= 89) and Q1-G4 (B, pqsfinder stability score= 53) showing that the more stable G4 is competing with binding compared to the less stable. **C.** Binding curves fitted using a non-linear decay curve indicate increased competition of the stable compared to the less stable G4 structure with the dsDNA fragment of recombination hotspot HSI.

## Discussion

PRDM9 is a major determinant of meiotic recombination hotspot location in many vertebrate species [5–9] and it is among the fastest evolving genes in humans [16]. Numerous PRDM9 alleles exist, differing in ZnF composition, DNA-binding specificity, and hotspot usage patterns [17, 18]. Despite its well-established sequence-specific DNA-binding activity, accumulating evidence suggests that the primary DNA sequence alone cannot fully explain PRDM9 occupancy across the genome. A substantial fraction of PRDM9-bound hotspots lacks the consensus motifs, and PRDM9 has also been reported to bind promoter-associated regions and CpG islands, albeit these sites are generally depleted of recombination activity [9, 87, 88]. In mice, a subset of PRDM9-binding sites which are not targeted for recombination are characterized by high CpG density and strong overlap with CpG islands. Many of these sites also lack the canonical PRDM9-binding motif [90, 91]. In line with that, in species lacking functional PRDM9 (e.g., dogs, plants, birds, and yeast), as well as in PRDM9 knockout mice, recombination preferentially localizes to promoter-associated features, particularly hypomethylated CpG islands [88, 92–94]. Together, these observations suggest that additional genomic determinants beyond the PRDM9 binding motif contribute to hotspot recognition and selection.

Although PRDM9 displayed generally lower binding affinities for G4 substrates than for canonical double-stranded DNA, potentially owing to differences in oligonucleotide length and flanking sequence context, our findings indicate that G4 structures may contribute to PRDM9-mediated hotspot targeting in addition to the canonical DNA-binding motif. This interpretation is further supported by our enrichment analyses, which showed that highly stable and experimentally validated G4s were preferentially enriched at PRDM9-binding sites, whereas less stable and non-canonical G4 motifs exhibited substantially weaker enrichment. This pattern supports the recognition of a specific structural feature rather than G-rich primary sequence alone, as a purely sequence-driven interaction would be expected to produce a similarly strong enrichment of predicted G4 motifs lacking experimental evidence of G4 formation. Moreover, because G4 structures are already present in untransfected HEK293 cells and are associated with discrete regions of open chromatin, PRDM9 may preferentially bind to pre-existing G4-associated accessible regions rather than actively inducing G4 formation. Although PRDM9, together with HELLS, acts as a pioneer factor that promotes chromatin opening, stable G4 structures may provide pre-existing binding platforms that facilitate the initial recruitment of PRDM9 and support subsequent chromatin remodeling. This model is further supported by our EMSA experiments, in which PRDM9 preferentially bound more stable G4 structures, and experimentally increasing G4 stability enhanced PRDM9 binding.

Beyond direct protein recruitment, G4 structures could also contribute to homology search during meiotic recombination. Given their documented roles in mediating long-range chromosomal interactions [95, 96], G4s may participate in stabilizing interactions between homologous chromosomes during meiosis as proposed in a recent preprint [97]. These observations may reflect two distinct scenarios: either G4 structures facilitate synaptonemal complex formation before PRDM9 binding, thereby enhancing PRDM9 recruitment to stable-anchored sites bringing homologous regions into close proximity, or PRDM9 initially binds to stable G4 structures, which subsequently promote homology search.

An important question arising from our findings concerns allele-specific interactions between different PRDM9 alleles and G4 structures. PRDM9 alleles differ in their DNA-binding specificity and hotspot usage across human populations [9, 16, 18]. Consequently, variation in hotspot activity may not be determined solely by differences in motif recognition, but also in their affinity for different G4 motifs, their stabilities or topologies, thereby influencing hotspot selection through both sequence-dependent and structure-dependent mechanisms. Such interactions could contribute to population-specific recombination landscapes and may help explain variation in hotspot usage that cannot be attributed to sequence motifs alone. In addition, recombination hotspots are dynamic genomic features that can erode or shift into cold spots over evolutionary time, for example through mutations at PRDM9 binding sites [98]. Spoken more generally, meiotic recombination is mutagenic [75, 99–101]. One mechanism that may contribute to hotspot turnover, in addition to changes in PRDM9 motifs, is the accumulation of sequence variants that alter G4-forming potential and stability. This could in turn reduce PRDM9 binding affinity and decrease hotspot activity. A compelling example is provided by the rs74149540 C/G single nucleotide polymorphism (SNP) at the MS32/D1S8 recombination [102], located immediately adjacent to a PRDM9-A-like motif: the G allele supports formation of a substantially more stable predicted G4 structure than the C allele (pqsfinder scores 50 versus 26) and is associated with high hotspot activity, whereas the destabilizing C allele acts as a cold or suppressor allele, strongly supported by our finding that PRDM9 preferentially recognizes more stable G4 structures (***Figure 8***). Although this correlation remains to be demonstrated genome-wide, the principle that G4 stability can modulate regulatory activity—as for example on transcription—is already well-established [41]

**Figure 8.**
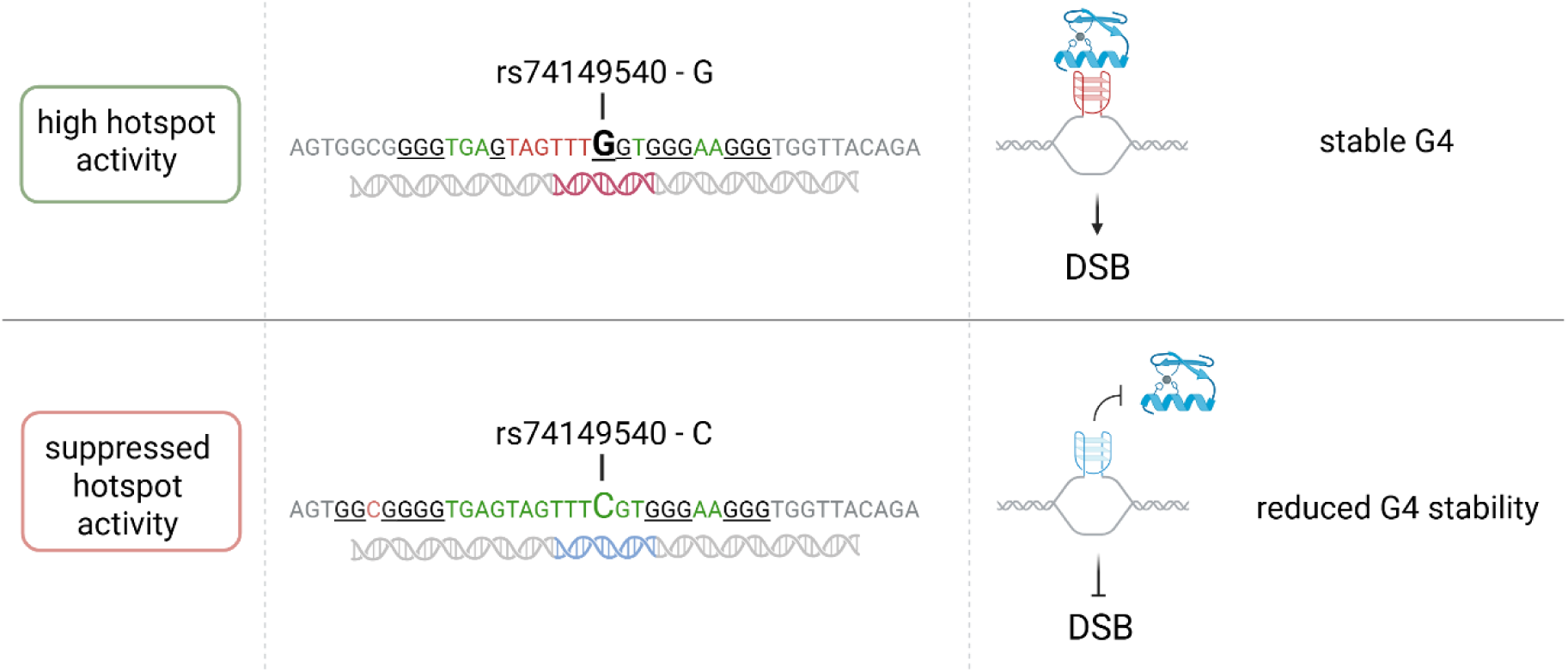
Proposed model for allele-specific effects on PRDM9 binding mediated by G4 stability at the MS32/D1S8 recombination hotspot. The rs74149540 C/G SNP is located adjacent to a PRDM9-A-like binding motif, with the C allele acting as a recombination-suppressing allele. The G allele is predicted to support formation of a stable G4 structure with a pqsfinder score of 50, whereas substitution with the C allele reduces the predicted G4 stability to a score of 26 and may consequently weaken PRDM9 binding. Underlined bases represent the G4 stems, green bases indicate nucleotides within the loops, red bases indicate bulges within the G4 stems, and grey bases represent the flanking sequences. Figure created with BioRender.com

Another interesting observation is that the ZnF protein YY1 suppresses G4-mediated homologous recombination and may utilize G4 structures to bind and protect G4-rich genes from recombination events [86]. Consistent with this model, G4s associated with YY1 exhibited lower recombination rates than G4s lacking YY1 occupancy [86]. Our findings suggest that PRDM9 preferentially associates with stable G4 structures linked to recombination hotspot activity. Given that YY1 and PRDM9 binding sites show little overlap [86], these observations indicate that G4 structures can serve as recognition elements for distinct ZnF proteins with opposing effects on recombination. One possibility is that YY1 occupancy protects specific G4-containing genomic regions from recombination, whereas PRDM9 preferentially targets a separate subset of G4-containing loci for hotspot activation, or that prior to PRDM9 binding, YY1 shields G4s in regions which should not undergo recombination events.

Together, these observations suggest that recombination hotspot activity is governed by a dynamic interplay among DNA sequence, G4 stability, chromatin accessibility, and epigenetic state. Future studies examining these factors across developmental stages, cell types, and evolutionary timescales will be important for understanding how recombination hotspots are established, maintained, and ultimately lost. More broadly, our findings support the emerging view that proteins involved in meiotic recombination like YY1 or PRDM9 can recognize and utilize DNA secondary structures in addition to primary nucleotide sequences. Rather than competing with canonical PRDM9 binding motifs, stable G4 structures may facilitate PRDM9 recruitment by promoting discrete accessible chromatin environments and providing structural binding platforms. This model offers a potential explanation for PRDM9 occupancy at sites lacking consensus motifs and provides a framework for reconciling PRDM9-dependent hotspot specification with the enrichment of promoter-associated and CpG island-associated genomic features. Ultimately, these results suggest that DNA secondary structure constitutes an additional layer of information contributing to the regulation and evolution of meiotic recombination landscapes.

Several limitations of this study should be acknowledged. Only a limited number of hotspot-associated G4 motifs and a single artificial G4 substrate were experimentally examined. Furthermore, the present study focused primarily on in vitro binding assays and computational analyses and therefore does not directly establish a causal role for G4 recognition in meiotic double-strand break formation or recombination activity in vivo. Future genome-wide, structural, and functional studies will be required to determine how broadly G4-mediated PRDM9 recruitment contributes to hotspot specification and whether different PRDM9 alleles exhibit distinct preferences for specific G4 architectures. In addition, approaches such as surface plasmon resonance (SPR) could provide quantitative insights into the kinetics and thermodynamics of PRDM9–G4 interactions. Nevertheless, the consistent relationship observed between G4 stability and PRDM9 binding, together with the enrichment of stable G4s at PRDM9 binding sites, provides strong evidence that DNA secondary structure contributes to PRDM9 target recognition and may represent an important component of recombination hotspot biology.

## Abbreviations

DSB: double-strand break
EMSA: electrophoretic mobility shift assay
G4: G-quadruplex
PDS: pyridostatin
PRDM9: PR domain containing protein 9
TSS: transcription start site,
ZnF: zinc finger

## Declarations

### Data availability

PRDM9, H3K4me3 and H3K36me3 ChIP-seq data from HEK293 cells were obtained from GEO database under the accession number GSE99407 [9, 48]. BG4 CUT&Tag data from HEK293 cells were obtained from the SRA database with entry numbers SRR14300945, SRR14879748, SRR26244907, SRR26244908, SRR26244909, SRR26244910, SRR26244911, SRR26244912, SRR25010684 and SRR25010685 [52–65]. DNA methylation of HEK293 cells was obtained from ENCODE experiment ENCSR794HFF [71]. Chromatin accessibility was assessed from ATAC-seq experiments from HEK293 cells deposited in the ENCODE database with the accession no. ENCSR205FUM [72]. Recombination hotspots maps were inferred from GEO database with the entry no. GSE59836 [2, 73].

### Supplementary Data Statement

Authors must add the following statement if their manuscript includes supplementary material for publication: ‘Supplementary Data are available at *NAR* Online’.

Supplementary File 1: consists of supplementary methods and figures

Supplementary File 2: contains the exact PRDM9-eYFP-pOPIN-M vector map

## Acknowledgments

We thank Nicolas Altemose for important input on their PRDM9 datasets. The authors acknowledge the computational resources and services provided by Salzburg Collaborative Computing (SCC), funded by the Federal Ministry of Education, Science and Research (BMBWF) and the State of Salzburg. Computational resources were in part provided by the e-INFRA CZ project (ID:90254), supported by the Ministry of Education, Youth and Sports of the Czech Republic. Computational resources were in part provided by the ELIXIR-CZ project (ID:90255), part of the international ELIXIR infrastructure.

## Authorś contribution

AL analyzed BG4 CUT&Tag and PRDM9 ChIP-seq data; TP, NP and MC performed enrichment analysis; TM, YS and ITB performed cloning of the PRDM9 construct; AL, MKH, NPH, YS and JH performed PRDM9 overexpression and extractions, EMSA and Western Blot experiments; ES performed CD spectra; A.L. conceptualized the study and wrote the manuscript, with input from YS, AR, JB and MHW; all authors read, corrected and approved the final manuscript.

## Consent for publication

Not applicable

