## Supplement_file_1 for "G-quadruplex structures act as a novel recognition motif for the meiosis-specific histone methyltransferase PRDM9"

### G-quadruplex structures act as a novel recognition motif for the meiosis-specific histone methyltransferase PRDM9

Mia K. Hartge[1,2+], Nuria Pfennig Hernandez[1,2+], Tomáš Pavlík[3], Nikol Poláková[3], Esther Schönauer[2,4], Julia Huber[1,2,5], Yasmin Striedner[6], Theresa Mair[6], Irene Tiemann-Boege[6], Matthias H. Weissensteiner[7], Johann Brandstetter[2,4], Angela Risch[1,2,5], Monika Cechova[3], Angelika Lahnsteiner[1,2,5*]

[1]Division of Cancer (Epi-) Genetics, Department of Biosciences and Medical Biology, University of Salzburg, 5020 Salzburg, Austria
[2]Center for Tumor Biology and Immunology (CTBI), University of Salzburg, 5020 Salzburg, Austria
[3]Faculty of Informatics, Masaryk University, 60200 Brno, Czech Republic
[4]Division of Structural Biology, Department of Biosciences and Medical Biology, University of Salzburg, 5020 Salzburg, Austria
[5]Cancer Cluster Salzburg, 5020 Salzburg, Austria
[6]Institute of Biophysics, Johannes Kepler University, 4040 Linz, Austria
[7]Institute of Avian Research, 26386 Wilhelmshaven, Germany

[+] contributed equally
* corresponding author: Angelika Lahnsteiner, Department of Biosciences and Medical Biology, University of Salzburg, 5020 Salzburg, Austria; Tel.: +43 662 8044 7223,

**Table of Content**

1. Supplementary Methods ... 2
1.1. Enrichment analysis: sampling of datasetA_matched and datasetB_global ... 2
1.2. Enrichment analysis: G-quadruplexes ... 2
1.3. Enrichment analysis: Global feature enrichment ... 3
1.4. Enrichment analysis: Grouped G4 enrichment ... 3
1.5. Enrichment analysis: Support-normalized G4 enrichment ... 4
1.6. Chromatin accessibility at G-quadruplex sites ... 4
1.7. Overlap between PRDM9-binding motifs and G-quadruplexes ... 5
1.8. PCR primer and ssDNA oligos for EMSA experiments: ... 6
1.9. Cloning of human PRDM9-A variant ... 7
2. Supplementary Figures ... 9
3. References ... 15

### 1. Supplementary Methods

#### 1.1. Enrichment analysis: sampling of datasetA_matched and datasetB_global

For the enrichment analysis, PRDM9-binding regions were used as the foreground dataset (n = 60,580; length range: 195–43,663 bp; mean: 1,153 bp) and compared with background regions sampled from the hg38 complement of PRDM9-binding regions. Two background datasets were generated, each containing three times as many regions as the foreground (n = 181,740): **datasetA_matched** and **datasetB_global**.

Both datasets were sampled to exactly reproduce the empirical length distribution of the foreground. This was achieved by randomly sampling background regions of predefined lengths until the target number of regions was obtained for every observed region length. **datasetA_matched** was additionally matched for GC content and sequence entropy using binning strategy: foreground regions were assigned to GC-content bins of 1 percentage point and entropy bins of 0.05 bits, and the corresponding target counts were scaled threefold. During sampling, candidate background regions were accepted only if capacity remained in both the relevant GC-content and entropy bins; otherwise, they were rejected. Sampling continued until all target counts were reached. The restrictive background datasetA_matched was introduced because PRDM9-binding regions have distinctive sequence properties, particularly high GC content, allowing PRDM9-binding associations to be distinguished more precisely from effects driven by sequence composition.

Length, GC-content, and entropy distributions are shown in Supplementary Fig. S1.

#### 1.2. Enrichment analysis: G-quadruplexes

G-quadruplex (G4) annotations were obtained from genome-wide predictions generated using pqsfinder (v. 2.28.0) [1]. The predicted G4 loci (n = 1,352,359; length range: 14–50 bp; mean length: 31.3 bp) were classified according to experimental support and sequence architecture.
Experimental support was determined by any overlap with G4 regions identified using CUT&Tag (n = 43,557; length range: 50–30,700 bp; mean length: 1,698.34 bp), where predicted loci overlapping at least one experimentally identified region were classified as supported (14.44% of sequence-predicted G4 loci), whereas predictions without such overlap were classified as unsupported.

Sequence architecture was classified as canonical or noncanonical. Canonical G4s (representing approximately 74% of pqsfinder predictions) conformed to the motif G(1–3)N(1–12)G(1–3)N(1–12)G(1–3)N(1–12)G(1–3). They were further categorized according to the maximum loop length as short (1–3 nt; 45.5% of canonical G4s), medium (4–6 nt; 51.2%), or long (7–12 nt; 39.1%). Noncanonical G4s comprised pqsfinder-predicted loci that did not satisfy these constraints (more complex motifs with mismatches or bulges in the G-stems).

Predictions were also divided into quartiles according to their pqsfinder stability scores (ranged from 52 to 395: Q1, 52–61; Q2, 62–68; Q3, 69–80; and Q4, 81–395). Q1 contained the lowest-scoring predictions and Q4 the highest-scoring predictions. Each locus was subsequently assigned a composite annotation combining its sequence class, score quartile, and support status, such as long–Q4–supported or noncanonical–Q2–unsupported.
PRDM9-binding regions, datasetA_matched and datasetB_global, were annotated using these composite categories. For each region and category, we calculated the number of intersecting G4 loci and the coverage fraction (covfrac), defined as the proportion of the region's base pairs covered by the union of G4 intervals belonging to that category. Overlapping intervals within the same category were merged before coverage calculation, ensuring that coverage represented the unique genomic

span rather than duplicated overlap. For analyses requiring a single loop-length category, including Figure 3, overlapping classifications were resolved using the hierarchy short < medium < long. Thus, a locus assigned to both medium and long categories was represented as long.

##### 1.3. Enrichment analysis: Global feature enrichment

For the global feature enrichment analysis (Figure 2A; Supplementary Figure S1), PRDM9-binding regions (n = 60,580) were compared separately with datasetA_matched and datasetB_global (each n = 181,740). All region-level features were combined into a single feature matrix. G4 features in this analysis comprised sequence predictions stratified by sequence class and score quartile, without stratification by experimental support.

For each feature, enrichment was summarized using the standardized mean difference:

$$d = \frac{\overline{x}_{PRDM9} - \overline{x}_{bg}}{\sqrt{\left(s_{PRDM9}^{2} + s_{bg}^{2}\right)/2}}$$

where $\overline{x}$ and $s$ denote the group mean and standard deviation. Analytical significance was assessed using a two-sided Welch's two-sample t-test, and p-values were adjusted using the Benjamini-Hochberg procedure to obtain $FDR_{exact}$.

Empirical significance was assessed using a block-wise Monte Carlo procedure. Background regions were ordered by genomic position and divided into consecutive superblocks of 130 regions (to preserve local genomic structure). In each of 200,000 iterations, superblocks were sampled with replacement to generate two synthetic groups of 60,580 regions each. For every feature, the absolute difference between the synthetic group means was compared with the observed absolute difference $|\Delta_{obs}|$. The empirical p-value was calculated using a pseudocount:

$$p_{(MC)} = \frac{1 + \left|\left\{i\colon \left|\Delta_{null,i}\right| \geq |\Delta_{obs}|\right\}\right|}{1 + N_{iter}}$$

Empirical p-values were adjusted independently using the Benjamini-Hochberg procedure to obtain$FDR_{MC}$. Final significance was defined conservatively as $\max(FDR_{exact}, FDR_{mc})$.

Features with FDRfinal $< 0.05$ were considered significant. Significant features were classified as enriched when $\overline{x}_{PRDM9} \geq \overline{x}_{bg}$ and depleted otherwise. This statistical framework was also used for the grouped G4 enrichment analysis described below.

##### 1.4. Enrichment analysis: Grouped G4 enrichment

To identify the G4 subtypes driving PRDM9 association, pqsfinder predictions were classified by sequence class, experimental support, and score quartile (described in Supplementary Methods 1.4). This produced 16 categories per support state, corresponding to 32 group-level features. For each category and genomic region, coverage fraction was calculated as the proportion of the region covered by the union of all G4 loci in that category (sequence-class annotations were non-exclusive; therefore, loci matching multiple classes contributed to each corresponding class-specific feature, as each category was analyzed independently.). PRDM9-binding regions were compared separately with datasetA_matched and datasetB_global using the Welch's t-test, block-wise Monte Carlo procedure, multiple-testing correction, and conservative final FDR criterion defined for the global feature analysis described in Supplementary Methods, Section 1.5. Features with $FDR_{final} < 0.05$ were considered significant.

##### 1.5. Enrichment analysis: Support-normalized G4 enrichment

To determine whether experimentally supported G4s were preferentially associated with PRDM9-binding regions beyond the enrichment expected from computational predictions alone, we performed a support-normalized analysis (Figure 2C–D). Sequence predictions were assigned sequence class, experimental support, and score quartile (described in Supplementary Methods 1.4).

Separately, an exact-sequence catalogue was generated by grouping predictions with identical sequences. Sequences for which ≥75% of loci were supported were classified as supported, those with ≤25% supported loci as unsupported, and the remainder as mixed. For the locus-level enrichment analysis, supported and unsupported G4 hits were counted separately within each PRDM9-binding and background region using two complementary approaches. First, support-specific hit rates were calculated using a pseudocount:

$$rate = \frac{hits_{s,W} + 0.5}{n_W}$$

where s ∈ {Supported, Unsupported} and W ∈ {PRDM9, background}, and $n_W$ is the number of regions in set W. Enrichment was expressed as the $\log_2$ rate ratio:

$$RR_s = \log_2\left(\frac{rate_{s,PRDM9}}{rate_{s,background}}\right)$$

The support-normalized contrast was defined as:

$$\Delta_{sup-unsup} = \log_2(RR_{sup}) - \log_2(RR_{unsup})$$

Positive values indicate stronger enrichment of supported than unsupported predictions. Two-sided p-values were calculated using a Poisson approximation. Confidence intervals were estimated from 5,000 bootstrap replicates in which complete regions were resampled with replacement, preserving the within-region dependence between supported and unsupported counts. P-values were adjusted using the Benjamini-Hochberg procedure.

Second, we estimated the conditional odds that a G4 overlapping a PRDM9-binding region was experimentally supported. Strata were jointly defined by G4 class, score quartile, length quartile, and background-prevalence quartile. Within each stratum, a $2 \times 2$ table compared PRDM9 versus background regions and supported versus unsupported G4 hits. After applying a 0.5 Haldane correction to all cells, the stratum-specific log odds ratio was calculated as:

$$\log\left(\frac{(a + 0.5)(d + 0.5)}{(b + 0.5)(c + 0.5)}\right)$$

where a, b, c, and d denote supported PRDM9, unsupported PRDM9, supported background, and unsupported background hits, respectively. Strata containing fewer than 20 total hits were excluded, and the remaining log odds ratios were pooled by inverse-variance weighting. The resulting adjusted odds ratio quantified whether G4s observed in PRDM9-binding regions had greater conditional odds of experimental support than G4s observed in background regions.

##### 1.6. Chromatin accessibility at G-quadruplex sites

To test whether G4-containing regions were associated with increased chromatin accessibility, we used the previously derived mean ATAC-seq signal (described in Methods, Enrichment analysis,

Chromatin accessibility) as the region-level accessibility measure. Within the PRDM9-binding, datasetA_matched, and datasetB_global sets, regions were classified separately for each G4 category according to whether they overlapped at least one experimental G4 interval or a sequence-predicted short, medium, long, or noncanonical G4 locus by at least 1 bp. These classifications were independent and non-mutually exclusive.

For each dataset and G4 category, the distributions of region-level mean accessibility were compared between G4-overlapping and non-overlapping regions using a one-sided Mann–Whitney U test, testing whether accessibility tended to be higher in G4-overlapping regions. Effect size was quantified using the rank-biserial correlation, and differences were additionally summarized as the ratio of median accessibility between overlapping and non-overlapping regions. In total, 15 comparisons were performed (five G4 categories and three region sets), with significance evaluated after Bonferroni correction.

##### 1.7. Overlap between PRDM9-binding motifs and G-quadruplexes

To examine the spatial association between PRDM9-binding motifs and G4s, we combined the motif hits (described in Methods, Enrichment analysis: PRDM9-binding sequence motifs) with the G4 annotations (described in Supplementary Methods, Section 1.4). Experimental G4 regions and sequence-predicted G4 loci were analyzed separately; sequence predictions were not stratified by experimental support. Within each region of the PRDM9-binding, datasetA_matched, and datasetB_global sets, all 17 PRDM9 motifs were scanned on both strands using the previously described PWM procedure. Hits detected at the same genomic position on both strands were counted once, retaining the higher-scoring hit.

For each motif, we counted hits overlapping by at least 1 bp with either experimental G4 regions or sequence-predicted short, medium, long, and noncanonical G4 loci. A motif hit was counted once per G4 category, regardless of how many intervals within that category it overlapped. Counts were aggregated across regions within each dataset to compare motif–G4 overlap patterns between PRDM9-binding regions and the two background datasets.

##### 1.8. PCR primer and ssDNA oligos for EMSA experiments

| Primer-ID | Sequence 5´-3´ | Note |
|---|---|---|
| F-HS2cen2-Btn | GAATCCGCTCCTGAAGTCAAA | Biotinylated forward primer |
| R-HS2cen2 | GAAGATCTCTGCACCCTGAAC | Reverse primer |
| G4-HS2cen2-FL-Btn | GAAATGCCTACGCCTGATGGCAGGGAAAGGTGGTGCTTTAACCCATGACT | Oligo for CD spectra and EMSA oligo |
| F-HS1cen-Btn | CTCTTCCTCACCCTTTCTCTTTC | Biotinylated forward primer |
| R-HS1cen | CACCAAGGTGTATAAGCTTTCTCT | Reverse primer |
| G4-HS1cen-FL-Btn | AGAAAAGACGGAGGGAGGGAAGGAAGGGAAGGAAAGGGGGAAGGAAGGGG | Oligo for CD spectra and EMSA oligo |
| F-usDNA | CTGCCTAAAGGTCAGAATCCACCATAGTGAGAGATAGCAAGTGCTGCT | Upper strand |
| R-usDNA-Btn | AGCAGCACTTGCTATCTCTCACTATGGTGGATTCTGACCTTTAGGCAG | Biotinylated lower strand |
| aG4 | GGGTCTGGGACTGGGACTGGGATTGGG | Biotinylated single stranded artificial G4 motif |
| mut-aG4 | GTGTCTGAGACTGTGACTGAGATTGAG | Biotinylated single stranded mutated artificial G4 motif |
| G4-Q4-short-Btn | GGGAGGGAGGGAGGG | Biotinylated single stranded G4 motif |
| G4-Q1-short-Btn | GGGAGAGGTGGGAGGG | Biotinylated single stranded G4 motif |
| hP9_5'UTR_newP_F | GGGCCCTTCTCACACTCAGAatt | PRDM9 cloning primer |
| hP9_Zn-fingerP_R | GTGTGTGGTGACCACATTTGTCttt | PRDM9 cloning primer |
| Not-TEV-hP9-FL_F | aaggaaaaaaGCGGCCGCaGAAAACTTGTATTTCCAGGGCgagcagaagctgatctcagaggaag acctgATGAGCCCTGAAAAGTCCCAAGAG | PRDM9 cloning primer |
| hP9_ZnF-Hind_R | CGTCGTAAGCTTGTGTGTGGTGACCACATTTGTcttt | PRDM9 cloning primer |
| eYFP_pos542_fwd | ACCACTACCAGCAGAACACC | PRDM9 sequencing primer |
| cMyc_sequ_rvs | GGTCTTCCTCTGAGATCAGCT | PRDM9 sequencing primer |

#### 1.9. Cloning of human PRDM9-A variant

##### 1.9.1. RNA isolation from human testicular tissue and cDNA synthesis

Total RNA was isolated from cryopreserved human testicular sperm extraction (TESE) samples using the ZR RNA MiniPrep™ Kit (Zymo Research, Austria) according to the manufacturer's instructions with minor modifications. Briefly, frozen TESE aliquots were lysed in RNA lysis buffer supplemented with dithiothreitol (DTT) and Proteinase K to facilitate tissue digestion. Following clarification of the lysate by centrifugation, RNA was purified on silica spin columns using ethanol-assisted binding, washed extensively, and eluted in RNase-free water. RNA concentration and purity were determined spectrophotometrically using a NanoDrop instrument (Thermo Fisher Scientific, Austria). RNA samples with A260/A280 ratios of approximately 1.8-2.0 were used for downstream applications.

First-strand cDNA was synthesized from up to 1 µg of total RNA using ProtoScript II Reverse Transcriptase (New England Biolabs, Austria) with a mixture of oligo(dT)15 and oligo(dT)18 primers according to the manufacturer's recommendations. Following reverse transcription, the enzyme was heat-inactivated, and cDNA was stored at -80°C until use.

##### 1.9.2. Amplification of human PRDM9 cDNA

Full-length human *PRDM9* was amplified from TESE-derived cDNA using Phusion High-Fidelity DNA Polymerase (Thermo Scientific, Austria). Primer pairs were designed to amplify the complete coding sequence (approximately 2.8 kb). PCR amplifications were performed using high-fidelity reaction buffer under optimized annealing conditions with controlled temperature ramping to improve amplification of GC-rich regions and repetitive zinc-finger sequences. PCR products were analysed by agarose gel electrophoresis, purified using silica-based PCR purification columns, and used for subsequent cloning. Cloning primers are listed in Supplementary Methods, Section 1.8.

##### 1.9.3. Cloning of PRDM9 PCR products into pJET1.2

Purified blunt-ended PCR products were cloned into the pJET1.2/blunt cloning vector (Thermo Scientific, Austria) according to the manufacturer's protocol. Ligation reactions were assembled using an approximately 1:3 molar ratio of vector to insert and incubated with T4 DNA ligase.

Recombinant plasmids were introduced into chemically competent *Escherichia coli* XL1-Blue cells (Agilent Technologies, Austria) by heat-shock transformation. Transformed bacteria were plated on LB agar supplemented with ampicillin and incubated overnight at 37°C. The pJET1.2 positive-selection system was used to enrich for recombinant plasmids.

##### 1.9.4. Colony screening and clone verification

Individual colonies were screened by colony PCR using vector-specific pJET1.2 forward and reverse sequencing primers supplied with the cloning kit. PCR products were analysed by agarose gel electrophoresis, and clones exhibiting inserts of the expected size were selected for subsequent plasmid preparation and sequence verification.

##### 1.9.5. Cloning of human PRDM9 into pOPIN-M

To generate N-terminal fluorescent fusion constructs, the full-length *PRDM9* coding sequence was re-amplified from sequence-verified pJET1.2 clones using Phusion High-Fidelity DNA Polymerase. Forward primers incorporated a NotI restriction site together with the pOPIN-compatible TEV protease recognition sequence, whereas reverse primers introduced a HindIII restriction site downstream of the coding sequence. PCR conditions were optimized using different annealing temperatures and amplification strategies to maximize recovery of the repetitive zinc-finger region.

PCR products were analysed by agarose gel electrophoresis, purified using silica membrane columns, and subjected to restriction digestion with NotI and HindIII. The pOPIN-M expression vector containing an N-terminal enhanced yellow fluorescent protein (eYFP) fusion cassette was digested with the same enzymes and purified prior to ligation. The pOPIN-M-eYFP fusion plasmid preparation was described in [2].

Insert and vector DNA were ligated using T4 DNA ligase at an approximately 1:3 molar vector-to-insert ratio. Recombinant plasmids were transformed into chemically competent *E. coli* XL1-Blue cells by heat shock and plated on LB agar containing ampicillin.

##### 1.9.6. Screening of pOPIN-M recombinant clones

Transformants were screened by colony PCR using vector-specific primers flanking the multiple cloning site (eYFP_pos542 forward and cMyc reverse primers). Colonies yielding PCR products of the expected size were selected for plasmid isolation and subsequent sequence verification to confirm correct insertion and reading-frame integrity of the *PRDM9* coding sequence with the N-terminal eYFP fusion. Sequencing primers are listed in Supplementary Methods, Section 1.8.

A synonymous A-to-G substitution was identified at nucleotide position 623, resulting in a codon change from GCA to GCG without altering the encoded amino acid (Ala175). Although both codons encode alanine, GCA is the more frequently used synonymous codon (23%) in homo sapiens according to [3], whereas GCG is used less frequently (11%), indicating a shift toward a less preferred codon. This synonymous variant may therefore influence codon usage bias without affecting the primary amino acid sequence and in principle, affect only translation efficiency without altering the primary amino acid sequence. However, robust PRDM9 expression was obtained in transfected HEK293 cells, indicating that the substitution did not substantially impair protein production under our experimental conditions. Because the plasmid was used solely to produce PRDM9 for subsequent in vitro binding experiments, all experiments were therefore performed using this construct.

#### 2. Supplementary Figures

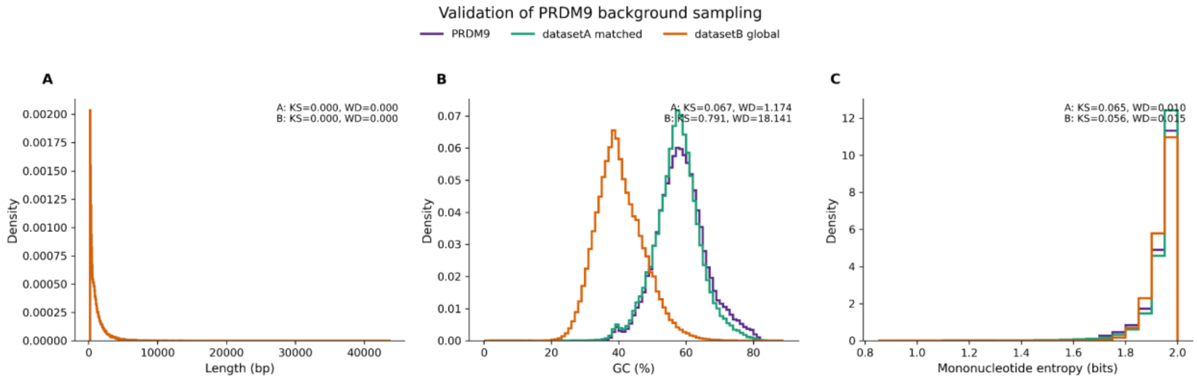


**Figure S1. Distribution of the lengths, GC content, and sequence entropy of the analyzed regions.** For the enrichment analysis, PRDM9-binding regions were used as the foreground dataset and compared with randomly sampled non-PRDM9-binding regions from the hg38 genome. Two background datasets were generated. datasetA_matched contained 181,740 non-PRDM9-binding regions sampled to match the length, GC-content, and entropy distributions of the PRDM9-binding regions. datasetB_global contained 181,740 non-PRDM9-binding regions sampled to match only the length distribution of the PRDM9-binding regions. The figure compares the length, GC-content, and entropy distributions across the PRDM9-binding regions, datasetA_matched, and datasetB_global regions. The sampling methodology is described in Supplementary Materials and Methods, Section 1.3.

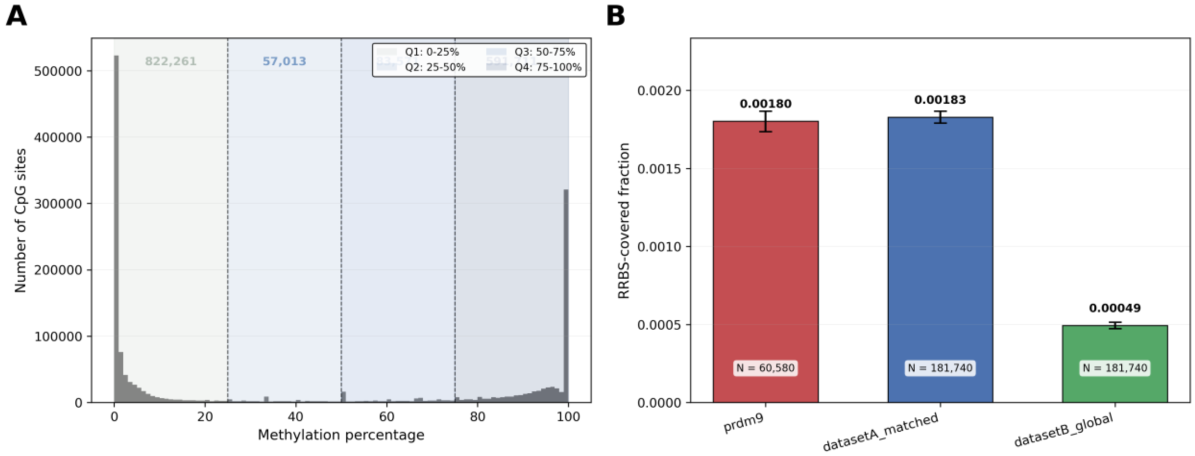


**Figure S2. Methylation overview. A.** Genome-wide distribution of methylation levels across 1,554,556 RRBS-covered CpG sites in HEK293 cells, showing characteristic bimodal pattern where 51.33% of sites are unmethylated or weakly methylated (<10%) and 35.25% are hypermethylated (>75%). Quartile boundaries (Q1–Q4) indicated by shaded regions. **B.** RRBS coverage fraction across PRDM9-binding windows (n=60,580), matched background (datasetA_matched, n=181,740), and global background (datasetB_global, n=181,740). Bars show mean ± 95% bootstrap CI. PRDM9-binding regions and matched background show comparable coverage (~0.0018), while global background shows 3.6× lower coverage (0.0005), reflecting GC/sequence composition matching in datasetA.

#### PRDM9 Binding Motifs - Sequence Logos

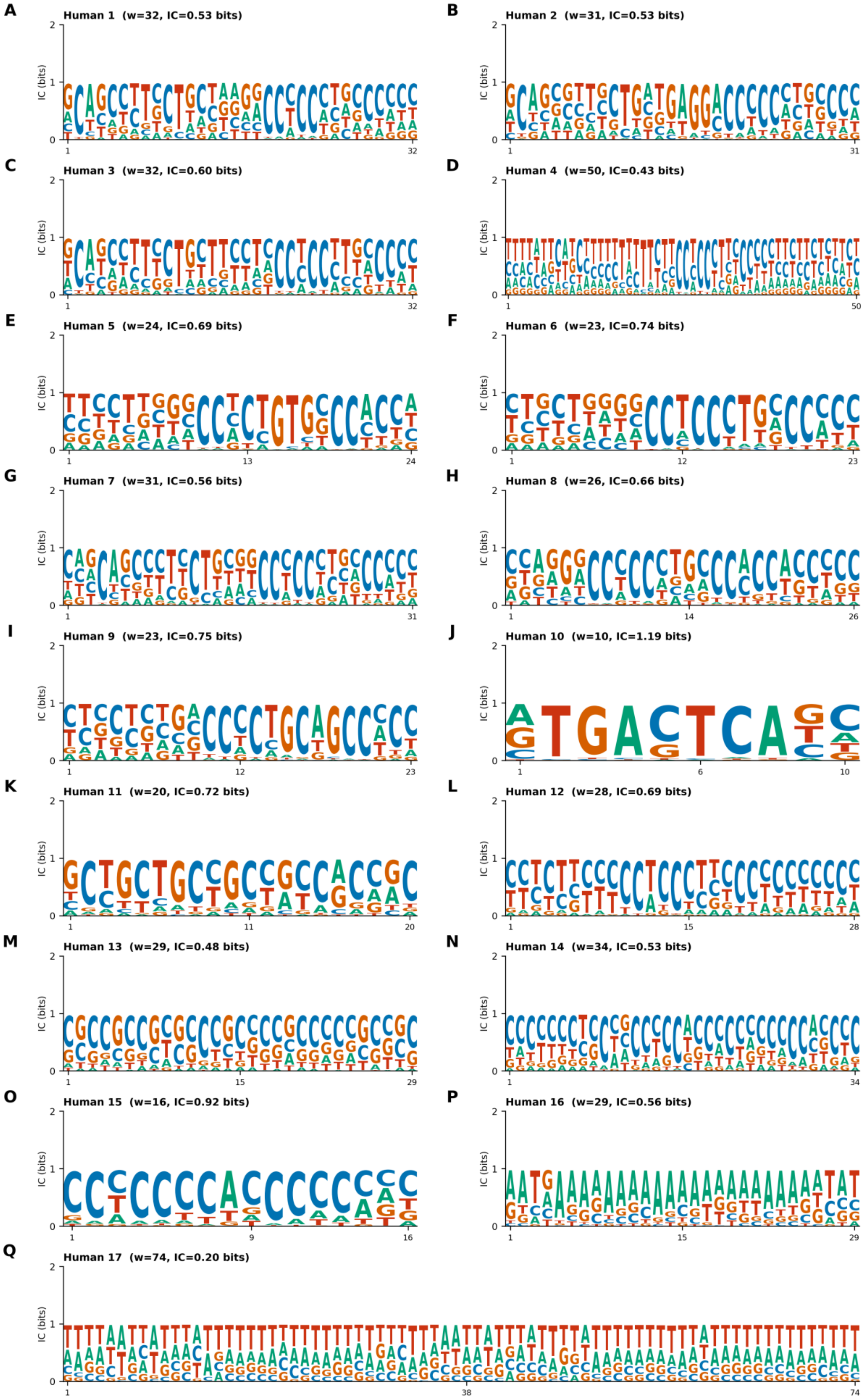

**Figure S3. PRDM9 Binding Motifs – Sequence Logos.** Sequence logos for all 17 human PRDM9 binding motifs given as position-weight matrices (PWMs) in MEME format. **A–Q**. Individual motif logos showing PWMs. Standard DNA coloring: A=green, C=blue, G=orange, T=red. All motifs: nsites=10,000, E-value=1e-5, scanned on both strands. Mean IC per motif ranges from 0.20 bits (Human 17) to 1.19 bits (Human 10, highest specificity). Consensus sequences extracted as maximum-probability base per position.

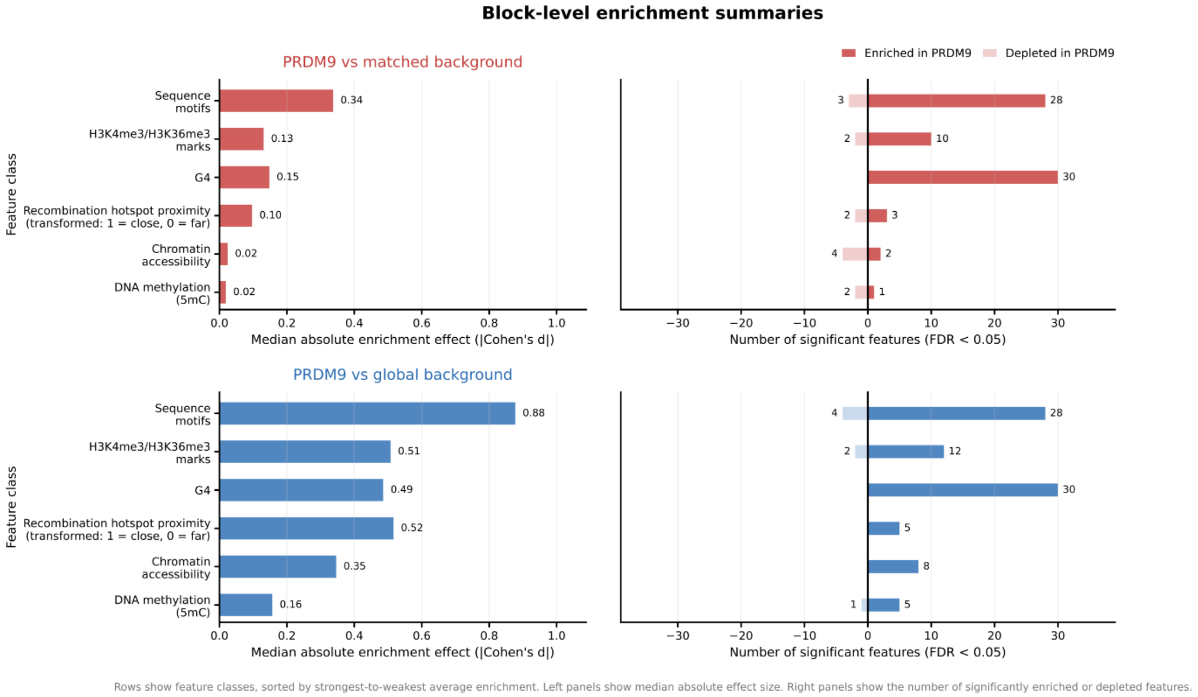


**Figure S4. Enrichment analysis confirms significant overlap of PRDM9 binding sites and G4 motifs.** Block-level enrichment analysis comparing genomic features enriched in PRDM9-associated regions relative to matched background regions. Left panels show median absolute enrichment effect sizes, expressed as Cohen's d, for blocks enriched above the background. Right panels show the number of significant features enriched or depleted in PRDM9-associated regions at FDR < 0.05. The upper panels summarize features enriched in PRDM9 compared with the matched background, whereas the lower panels summarize features enriched in the matched background compared with PRDM9. Features include sequence context, chromatin-associated marks, G4 formation, recombination hotspot proximity, chromatin accessibility, and DNA methylation.

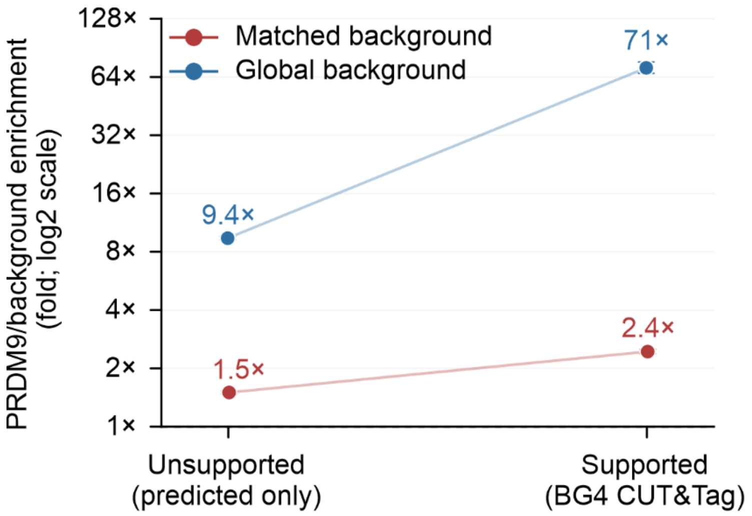


**Figure S5. Enrichment of experimentally supported and unsupported predicted G4 motifs at PRDM9-binding windows.** G4 enrichment was markedly stronger for experimentally supported G4s than for computationally predicted motifs lacking experimental support in HEK293 cells.

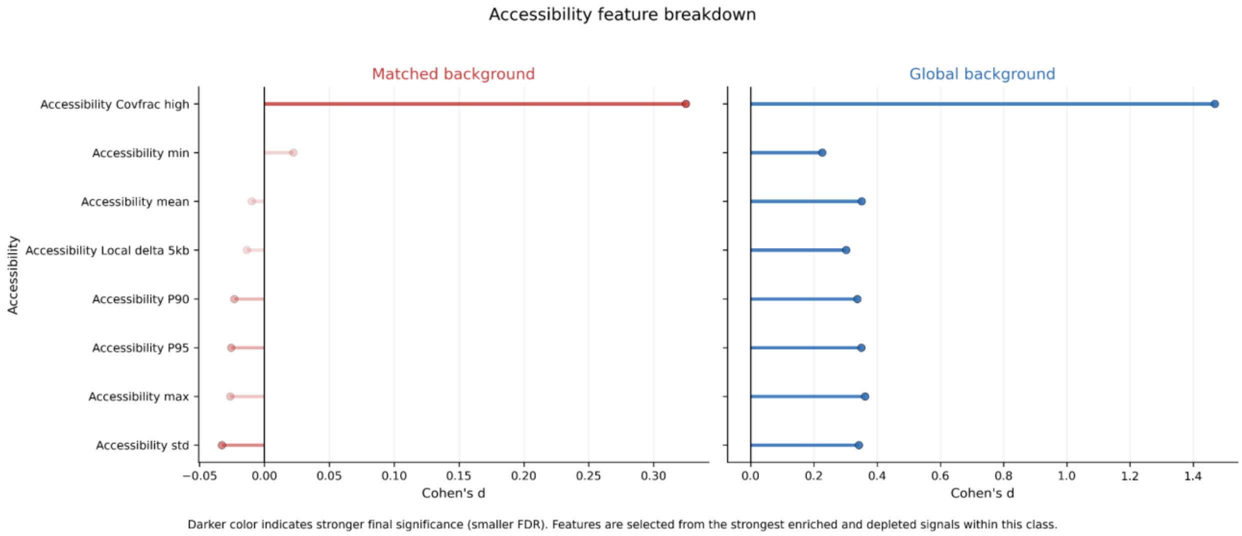


**Figure S6. Chromatin accessibility features at PRDM9-binding regions.** Standardized effect sizes are shown as Cohen's *d* relative to a sequence-matched background and a global genomic background. Accessibility was quantified from the HEK293 ATAC-seq signal using 25-bp bins. "CovFrac high" denotes the fraction of bins within each region with an ATAC-seq signal above the high-accessibility threshold of 0.253. Local contrast represents the difference between the mean signal within the region and the mean signal across the adjacent 5-kb flanks. Positive values indicate higher accessibility at PRDM9-binding regions.

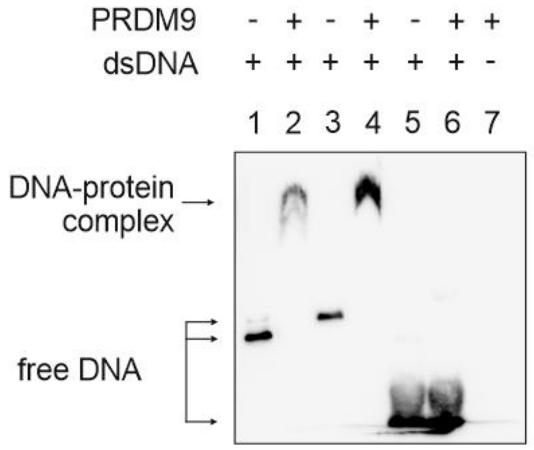


**Figure S7. PRDM9 selectively binds to recombination hotspots.** The EMSA shows dsDNA of HSI in lanes 1 and 2, dsDNA of HSII in lanes 3 and 4, and unspecific control DNA in lanes 5 and 6. Lanes 1, 3, and 5 contain DNA only, while lanes 2, 4, and 6 contain DNA + PRDM9. Lane 7 shows the protein-only control.

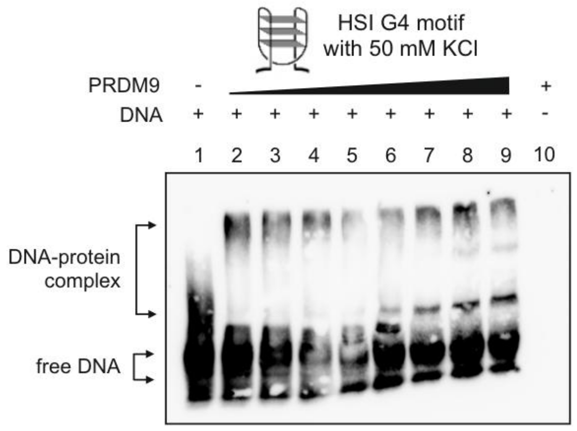


**Figure S8. EMSA of the HSI G4 motif in the presence of 50 mM KCl.** EMSA experiments were performed using the biotinylated G4 motif from the HSI center together with PRDM9 in EMSA binding buffer containing 50 mM KCl. The broad smear of the DNA-only band in lane 1 suggests that several DNA structures coexist and are likely not fully stable under these conditions. This may explain the presence of two shifted bands with different migration heights for the DNA–protein complex.
