## Supplement_file_2 for "G-quadruplex structures act as a novel recognition motif for the meiosis-specific histone methyltransferase PRDM9"

Wed Jul 29, 2026 20:44 CEST
pOPIN-M containing eYFP-hPrdm9A-FL.dna

```
              10        20        30        40        50        60        70        80        90       100
          *    *    *    *    *    *    *    *    *    *    *    *    *    *    *    *    *    *    *    *
                                                    42 AciI
                                                 39 NlaIII
                                                38 SphI
                                                38 NspI     50 MluCI
       1 HaeIII                                 38 Cac8I   49 ApoI                                   92 HpyCH4V
       |                                        ||  |      ||                                        |
     1 ggcctctaaacgggtcttgaggggtttttttgctgaaagcatgcggaggaaattctccttgaagtttccctggtgttcaaagtaaaggagtttgcaccaga 100
       G  L  *  T  G  L  E  G  F  F  A  E  S  M  R  R  K  F  S  L  K  F  P  W  C  S  K  *  R  S  L  H  Q  T

              110       120       130       140       150       160       170       180       190       200
          *    *    *    *    *    *    *    *    *    *    *    *    *    *    *    *    *    *    *    *
                                                                                 168 BfaI
                                                                              165 HhaI
                                                                             164 HaeII
                                          128 MseI               152 RsaI    164 AfeI                          198 MluCI
                110 Hpy8I 120 HpaII                              151 TatI  161 MseI                           197 MfeI
                |         |       |                              ||        |  ||  |                           ||
   101 cgcacctctgttcactggtccggcgtattaaaacacgatacattgttattagtacatttattaagcgctagattctgtgcgttgttgatttacagacaat 200
         H  L  C  S  L  V  R  R  I  K  T  R  Y  I  V  I  S  T  F  I  K  R  *  I  L  C  V  V  D  L  Q  T  I

              210       220       230       240       250       260       270       280       290       300
          *    *    *    *    *    *    *    *    *    *    *    *    *    *    *    *    *    *    *    *
                                                                                                          300 SspI
                                                                                                       297 MseI
             207 HpyCH4IV         228 MluCI                                                           296 DraI
            206 SnaBI           227 ApoI                                                              295 SwaI
            206 BsaAI    219 MluCI   231 PsiI                                                        294 MluCI
           205 RsaI 214 MseI    225 MseI                                                            293 ApoI
           |||      |    |      | ||  |                                                             |||||  |
   201 tgttgtacgtattttaataattcattaaatttataatctttagggtggtatgttagagcgaaaatcaaatgattttcagcgtctttatatctgaatttaa 300
        V  V  R  I  L  I  I  H  *  I  Y  N  L  *  G  G  M  L  E  R  K  S  N  D  F  Q  R  L  Y  I  *  I  *

              310       320       330       340       350       360       370       380       390       400
          *    *    *    *    *    *    *    *    *    *    *    *    *    *    *    *    *    *    *    *
          304 MseI                       335 TaqI
          |                              |
   301 atattaaatcctcaatagatttgtaaaataggtttcgattagtttcaaacaagggttgtttttccgaaccgatggctggactatctaatggattttcgct 400
       I  L  N  P  Q  *  I  C  K  I  G  F  D  *  F  Q  T  R  V  V  F  P  N  R  W  L  D  Y  L  M  D  F  R  S

              410       420       430       440       450       460       470       480       490       500
          *    *    *    *    *    *    *    *    *    *    *    *    *    *    *    *    *    *    *    *
                                                                                                  486 HpyCH4IV
                                                                                                 485 ZraI
                                                  440 AluI                                       485 AatII
                                                438 BfaI 447 TaqI                              483 TaqI      497 SspI
                                                | |      |                                     | ||          |
   401 caacgccacaaaacttgccaaatcttgtagcagcaatctagctttgtcgatattcgtttgtgttttgttttgtaataaaggttcgacgtcgttcaaaata 500
         T  P  Q  N  L  P  N  L  V  A  A  I  *  L  C  R  Y  S  F  V  F  C  F  V  I  K  V  R  R  R  S  K  Y

              510       520       530       540       550       560       570       580       590       600
          *    *    *    *    *    *    *    *    *    *    *    *    *    *    *    *    *    *    *    *
                                                       543 MluCI
                                                     542 MfeI
                                                  539 RsaI                564 MseI
                                                538 TatI         555 Hpy8I
                                                538 BsrGI      553 HpyCH4IV       572 AluI
           505 HhaI                            537 Hpy8I    550 TaqI   561 HpyCH4IV            585 MseI
           |                                   |||  ||      |  | |     |  |       |            |
   501 ttatgcgcttttgtatttctttcatcactgtcgttagtgtacaattgactcgacgtaaacacgttaaatagagcttggacatatttaacatcgggcgtgt 600
        Y  A  L  L  Y  F  F  H  H  C  R  *  C  T  I  D  S  T  *  T  R  *  I  E  L  G  H  I  *  H  R  A  C

              610       620       630       640       650       660       670       680       690       700
          *    *    *    *    *    *    *    *    *    *    *    *    *    *    *    *    *    *    *    *
                                                                                                      695 MluCI
                                                                                                    693 MseI
        602 AluI  612 HaeIII                                       660 BbsI                          692 AseI
        |         |                                                |                                ||  |
   601 tagctttattaggccgattatcgtcgtcgtcccaaccctcgtcgttagaagttgcttccgaagacgattttgccatagccacacgacgcctattaattgt 700
       *  L  Y  *  A  D  Y  R  R  R  P  N  P  R  R  *  K  L  L  P  K  T  I  L  P  *  P  H  D  A  Y  *  L  C

              710       720       730       740       750       760       770       780       790       800
          *    *    *    *    *    *    *    *    *    *    *    *    *    *    *    *    *    *    *    *
                          719 DpnI
                       716 BstUI                           752 DpnI
                      715 AciI                            751 PvuI
                  711 HpyCH4IV 724 MluCI                  750 SgfI                                  793 AluI
                 710 BmgBI    723 ApoI     736 AluI  746 MluCI                                 788 PsiI
                 ||   ||  |   ||           |         |   |||                                   |    |
   701 gtcggctaacacgtccgcgatcaaatttgtagttgagctttttggaattgcgatcgcataacttcgtatagcatacattatacgaagttataagctcgga 800
```

```
       R  L  T  R  P  R  S  N  L  *  L  S  F  L  E  L  R  S  H  N  F  V  *  H  T  L  Y  E  V  I  S  S  E


             810       820       830       840       850       860       870       880       890       900
         *    *    *    *    *    *    *    *    *    *    *    *    *    *    *    *    *    *    *    *
                                   831 AciI
                                 829 BsrBI                                                             899 NlaIII
                               827 Cac8I                                                              898 PciI
          806 HhaI          824 AciI       839 AluI     852 AciI                                      898 NspI
          |                 |  | | |       |            |                                             ||
 801 acgctgcgctcggtcgttcggctgcggcgagcggtatcagctcactcaaaggcggtaatacggttatccacagaatcaggggataacgcaggaaagaaca 900
      R  C  A  R  S  F  G  C  G  E  R  Y  Q  L  T  Q  R  R  *  Y  G  Y  P  Q  N  Q  G  I  T  Q  E  R  T


             910       920       930       940       950       960       970       980       990      1000
         *    *    *    *    *    *    *    *    *    *    *    *    *    *    *    *    *    *    *    *
                                                      950 Cac8I
                                                950 BstUI
                 913 Cac8I                     943 AciI
                912 HaeIII 923 HaeIII        941 HaeIII                    971 AciI                  997 TaqI
                ||         |                 | ||     |                    |                         |
 901 tgtgagcaaaaggccagcaaaaggccaggaaccgtaaaaaggccgcgttgctggcgtttttccataggctccgcccccctgacgagcatcacaaaaatcga 1000
     C  E  Q  K  A  S  K  R  P  G  T  V  K  R  P  R  C  W  R  F  S  I  G  S  A  P  *  R  A  S  Q  K  S  T


            1010      1020      1030      1040      1050      1060      1070      1080      1090      1100
         *    *    *    *    *    *    *    *    *    *    *    *    *    *    *    *    *    *    *    *
                                                                          1070 BssSI
                                                                    1064 AluI  1075 HhaI             1097 AciI
                                                                    |     |    |                     |
1001 cgctcaagtcagaggtggcgaaacccgacaggactataaagataccaggcgtttccccctggaagctccctcgtgcgctctcctgttccgaccctgccgc 1100
       L  K  S  E  V  A  K  P  D  R  T  I  K  I  P  G  V  S  P  W  K  L  P  R  A  L  S  C  S  D  P  A  A


            1110      1120      1130      1140      1150      1160      1170      1180      1190      1200
         *    *    *    *    *    *    *    *    *    *    *    *    *    *    *    *    *    *    *    *
        1104 HpaII                            1142 HhaI
       1103 BsaWI   1116 AciI                1141 HaeII                                                 1200 AluI
       ||           |                        ||                                                         |
1101 ttaccggatacctgtccgcctttctcccttcgggaagcgtggcgctttctcaatgctcacgctgtaggtatctcagttcggtgtaggtcgttcgctccaa 1200
      Y  R  I  P  V  R  L  S  P  F  G  K  R  G  A  F  S  M  L  T  L  *  V  S  Q  F  G  V  G  R  S  L  Q


            1210      1220      1230      1240      1250      1260      1270      1280      1290      1300
         *    *    *    *    *    *    *    *    *    *    *    *    *    *    *    *    *    *    *    *
                1212 HpyCH4V                           1251 HpaII
               1211 Hpy8I                     1242 HhaI
               1211 ApaLI                1237 AciI    1250 BsaWI                 1277 HpaII
               ||                        |    |       ||                        |
1201 gctgggctgtgtgcacgaaccccccgttcagcccgaccgctgcgccttatccggtaactatcgtcttgagtccaacccggtaagacacgacttatcgcca 1300
     A  G  L  C  A  R  T  P  R  S  A  R  P  L  R  L  I  R  *  L  S  S  *  V  Q  P  G  K  T  R  L  I  A  T


            1310      1320      1330      1340      1350      1360      1370      1380      1390      1400
         *    *    *    *    *    *    *    *    *    *    *    *    *    *    *    *    *    *    *    *
                                                   1347 AciI                  1374 HaeIII       1392 BfaI
                                                   |                          |                 |
1301 ctggcagcagccactggtaacaggattagcagagcgaggtatgtaggcggtgctacagagttcttgaagtggtggcctaactacggctacactagaagaa 1400
       G  S  S  H  W  *  Q  D  *  Q  S  E  V  C  R  R  C  Y  R  V  L  E  V  V  A  *  L  R  L  H  *  K  N


            1410      1420      1430      1440      1450      1460      1470      1480      1490      1500
         *    *    *    *    *    *    *    *    *    *    *    *    *    *    *    *    *    *    *    *
                                                                    1464 DpnI                  1491 AciI
                    1416 HhaI                                1457 AluI 1467 HpaII     1482 AciI
                    |                                        |      |  |              |        |
1401 cagtatttggtatctgcgctctgctgaagccagttaccttcggaaaaagagttggtagctcttgatccggcaaacaaaccaccgctggtagcggtggttt 1500
      S  I  W  Y  L  R  S  A  E  A  S  Y  L  R  K  K  S  W  *  L  L  I  R  Q  T  N  H  R  W  *  R  W  F


            1510      1520      1530      1540      1550      1560      1570      1580      1590      1600
         *    *    *    *    *    *    *    *    *    *    *    *    *    *    *    *    *    *    *    *
             1509 Cac8I      1525 HhaI     1539 DpnI  1550 DpnI
            1508 HpyCH4V    1524 BstUI    1538 BstYI 1549 BstYI          1569 MseI                 1595 DpnI
            ||              ||            ||         ||                  |                         |
1501 ttttgtttgcaagcagcagattacgcgcagaaaaaaaggatctcaagaagatcctttgTTACCAATGCTTAATCAGTGAGGCACCTATCTCAGCGATCTG 1600
     F  C  L  Q  A  A  D  Y  A  Q  K  K  R  I  S  R  R  S  F  V  T  N  A  *  S  V  R  H  L  S  Q  R  S  V


            1610      1620      1630      1640      1650      1660      1670      1680      1690      1700
         *    *    *    *    *    *    *    *    *    *    *    *    *    *    *    *    *    *    *    *
                                                                                                    1696 BstUI
                                                                                         1685 HpyCH4V  1699 BsaI
                                                                              1674 HaeIII          1695 AciI
                                                                              |          |         ||  |
1601 TCTATTTCGTTCATCCATAGTTGCCTGACTCCCCGTCGTGTAGATAACTACGATACGGGAGGGCTTACCATCTGGCCCCAGTGCTGCAATGATACCGCGA 1700
       Y  F  V  H  P  *  L  P  D  S  P  S  C  R  *  L  R  Y  G  R  A  Y  H  L  A  P  V  L  Q  *  Y  R  E
```

```
            1710      1720      1730      1740      1750      1760      1770      1780      1790      1800
         *    *    *    *    *    *    *    *    *    *    *    *    *    *    *    *    *    *    *    *
                                                   1747 HpaII   1760 HhaI                 1786 AciI
                 1713 HpaII                   1742 Cac8I  1754 HaeIII          1775 HpyCH4V
                 |                            |    |      |     |              |          |
1701 GACCCACGCTCACCGGCTCCAGATTTATCAGCAATAAACCAGCCAGCCGGAAGGGCCGAGCGCAGAAGTGGTCCTGCAACTTTATCCGCCTCCATCCAGT 1800
      T  H  A  H  R  L  Q  I  Y  Q  Q  *  T  S  Q  P  E  G  P  S  A  E  V  V  L  Q  L  Y  P  P  P  S  S

            1810      1820      1830      1840      1850      1860      1870      1880      1890      1900
         *    *    *    *    *    *    *    *    *    *    *    *    *    *    *    *    *    *    *    *
                                                              1858 HpyCH4IV
          1806 MluCI      1822 BfaI                          1857 AclI
        1804 MseI       1820 AluI                       1852 FspI
       1803 AseI  1814 HpaII                    1843 MseI 1853 HhaI
       || |       |     | |                     |        ||   ||
1801 CTATTAATTGTTGCCGGGAAGCTAGAGTAAGTAGTTCGCCAGTTAATAGTTTGCGCAACGTTGTTGCCATTGCTACAGGCATCGTGGTGTCACGCTCGTC 1900
     L  L  I  V  A  G  K  L  E  *  V  V  R  Q  L  I  V  C  A  T  L  L  P  L  L  Q  A  S  W  C  H  A  R  R

            1910      1920      1930      1940      1950      1960      1970      1980      1990      2000
         *    *    *    *    *    *    *    *    *    *    *    *    *    *    *    *    *    *    *    *
                            1924 HpaII                           1961 NlaIII
                           1923 BsaWI                     1954 DpnI              1977 AciI              2000 DpnI
                        1920 AluI      1936 DpnI      1951 NlaIII      1968 HpyCH4V   1983 AluI      1999 PvuI
                        |  ||           |              |  |      |      |        |     |               ||
1901 GTTTGGTATGGCTTCATTCAGCTCCGGTTCCCAACGATCAAGGCGAGTTACATGATCCCCCATGTTGTGCAAAAAAGCGGTTAGCTCCTTCGGTCCTCCG 2000
       L  V  W  L  H  S  A  P  V  P  N  D  Q  G  E  L  H  D  P  P  C  C  A  K  K  R  L  A  P  S  V  L  R

            2010      2020      2030      2040      2050      2060      2070      2080      2090      2100
         *    *    *    *    *    *    *    *    *    *    *    *    *    *    *    *    *    *    *    *
                           2023 AciI                            2061 MluCI
                         2021 HaeIII      2039 NlaIII     2056 HpyCH4V      2075 NlaIII
                         | |               |                |    |             |
2001 ATCGTTGTCAGAAGTAAGTTGGCCGCAGTGTTATCACTCATGGTTATGGCAGCACTGCATAATTCTCTTACTGTCATGCCATCCGTAAGATGCTTTTCTG 2100
      S  L  S  E  V  S  W  P  Q  C  Y  H  S  W  L  W  Q  H  C  I  I  L  L  L  S  C  H  P  *  D  A  F  L

            2110      2120      2130      2140      2150      2160      2170      2180      2190      2200
         *    *    *    *    *    *    *    *    *    *    *    *    *    *    *    *    *    *    *    *
               2111 RsaI                                                                     2190 HhaI
              2110 TatI                                                                     2189 BstUI
              2110 ScaI                          2144 AciI             2166 HpaII           2188 AciI
              ||                                |                     |                     |||
2101 TGACTGGTGAGTACTCAACCAAGTCATTCTGAGAATAGTGTATGCGGCGACCGAGTTGCTCTTGCCCGGCGTCAATACGGGATAATACCGCGCCACATAG 2200
     *  L  V  S  T  Q  P  S  H  S  E  N  S  V  C  G  D  R  V  A  L  A  R  R  Q  Y  G  I  I  P  R  H  I  A

            2210      2220      2230      2240      2250      2260      2270      2280      2290      2300
         *    *    *    *    *    *    *    *    *    *    *    *    *    *    *    *    *    *    *    *
                                                                                                        2300 HpyCH4V
                                   2231 HpyCH4IV                     2265 AciI         2283 TaqI       2299 Hpy8I
            2208 MseI             2230 AclI                   2258 DpnI        2275 DpnI               2299 ApaLI
           2207 DraI           2227 XmnI                     2257 BstYI       2274 BstYI           2296 BssSI
           ||                  |  ||                         ||      |        ||       |            |  ||
2201 CAGAACTTTAAAAGTGCTCATCATtggaaaacgttcttcggggcgaaaactctcaaggatcttaccgctgttgagatccagttcgatgtaacccactcgt 2300
       E  L  *  K  C  S  S  L  E  N  V  L  R  G  E  N  S  Q  G  S  Y  R  C  *  D  P  V  R  C  N  P  L  V

            2310      2320      2330      2340      2350      2360      2370      2380      2390      2400
         *    *    *    *    *    *    *    *    *    *    *    *    *    *    *    *    *    *    *    *
               2311 DpnI                                                      2374 AciI
               |                                                              |
2301 gcacccaactgatcttcagcatcttttactttcaccagcgtttctgggtgagcaaaaacaggaaggcaaaatgccgcaaaaaagggaataagggcgacac 2400
      H  P  T  D  L  Q  H  L  L  L  S  P  A  F  L  G  E  Q  K  Q  E  G  K  M  P  Q  K  R  E  *  G  R  H

            2410      2420      2430      2440      2450      2460      2470      2480      2490      2500
         *    *    *    *    *    *    *    *    *    *    *    *    *    *    *    *    *    *    *    *
                                                                                2476 BstUI
                                                                               2475 HhaI
                                                                              2474 BstUI
                                                                             2473 AciI           2493 MseI
                                      2434 SspI                         2468 NlaIII      2485 HpyCH4V
                                      |                                 |    ||||        |       |
2401 ggaaatgttgaatactcatactcttcctttttcaatattattgaagcatttatcagggttattgtctcatgtccgcgcgtttcctgcatcttttaatcaa 2500
     G  N  V  E  Y  S  Y  S  S  F  F  N  I  I  E  A  F  I  R  V  I  V  S  C  P  R  V  S  C  I  F  *  S  N

            2510      2520      2530      2540      2550      2560      2570      2580      2590      2600
         *    *    *    *    *    *    *    *    *    *    *    *    *    *    *    *    *    *    *    *
                                  2530 TatI
                                  2530 BsrGI
                            2524 HpaII
                         2521 HhaI                                       2569 HpyCH4IV
                        2520 BstUI 2531 RsaI                        2564 HpyCH4V
```

```
                        ||  |     ||                                |    |
2501 atcccaagatgtgtataaacgcgccggtatgtacaggaagaggtttatactaaactgttacattgcaaacgtggtttcgtgtgccaagtgtgaaaaccga 2600
       P  K  M  C  I  N  A  P  V  C  T  G  R  G  L  Y  *  T  V  T  L  Q  T  W  F  R  V  P  S  V  K  T  D


            2610      2620      2630      2640      2650      2660      2670      2680      2690      2700
         *    *    *    *    *    *    *    *    *    *    *    *    *    *    *    *    *    *    *    *
                                                                          2670 MseI
                                                                  2662 HpyCH4V
                                                                 2661 NsiI
        2604 MseI                                               2660 NlaIII
        |                                                       |||       |
2601 tgtttaatcaaggctctgacgcatttctacaaccacgactccaagtgtgtgggtgaagtcatgcatcttttaatcaaatcccaagatgtgtataaaccac 2700
      V  *  S  R  L  *  R  I  S  T  T  T  T  P  S  V  W  V  K  S  C  I  F  *  S  N  P  K  M  C  I  N  H


            2710      2720      2730      2740      2750      2760      2770      2780      2790      2800
         *    *    *    *    *    *    *    *    *    *    *    *    *    *    *    *    *    *    *    *
                                   2731 AluI
                             2725 TaqI
                            2724 SalI
                            2724 Hpy8I                    2754 HpyCH4V                             2799 PsiI
                            2724 HincII          2745 Cac8I     2760 BsaI         2778 MluCI         2797 MluCI
                            ||     |             |        |     |                 |                  | |
2701 caaactgccaaaaaatgaaaactgtcgacaagctctgtccgtttgctggcaactgcaagggtctcaatcctatttgtaattattgaataataaaacaatt 2800
     Q  T  A  K  K  *  K  L  S  T  S  S  V  R  L  L  A  T  A  R  V  S  I  L  F  V  I  I  E  *  *  N  N  Y


            2810      2820      2830      2840      2850      2860      2870      2880      2890      2900
         *    *    *    *    *    *    *    *    *    *    *    *    *    *    *    *    *    *    *    *
                                                                                     2881 BstUI
                                                                                    2880 MluI
               2811 MluCI
              2810 ApoI     2824 MseI                                       2872 MluCI      2888 PsiI
              ||            |                                               |       ||      |
2801 ataaatgtcaaatttgtttttattaacgatacaaaccaaacgcaacaagaacatttgtagtattatctataattgaaaacgcgtagttataatcgctga 2900
       K  C  Q  I  C  F  L  L  T  I  Q  T  K  R  N  K  N  I  C  S  I  I  Y  N  *  K  R  V  V  I  I  A  E


            2910      2920      2930      2940      2950      2960      2970      2980      2990      3000
         *    *    *    *    *    *    *    *    *    *    *    *    *    *    *    *    *    *    *    *
             2909 MseI
            2908 DraI                     2938 MluCI
        2904 SspI                       2936 MseI        2953 MluCI         2972 BfaI     2986 BbsI
        |   ||                          | |              |                  |             |
2901 ggtaatatttaaaatcattttcaaatgattcacagttaatttgcgacaatataatttatttcacataaactagacgccttgtcgtcttcttcttcgta 3000
      V  I  F  K  I  I  F  K  *  F  T  V  N  L  R  Q  Y  N  F  I  F  T  *  T  R  R  L  V  V  F  F  F  V


            3010      3020      3030      3040      3050      3060      3070      3080      3090      3100
         *    *    *    *    *    *    *    *    *    *    *    *    *    *    *    *    *    *    *    *
                                       3035 MseI                               3083 MluCI
                                     3033 MluCI                               3082 ApoI
                                     | |                                      ||
3001 ttccttctcttttcatttttctcttcataaaaattaacatagttattatcgtatccatatatgtatctatcgtatagagtaaatttttttgttgtcataa 3100
     F  L  L  F  F  I  F  L  F  I  K  I  N  I  V  I  I  V  S  I  Y  V  S  I  V  *  S  K  F  F  V  V  I  N


            3110      3120      3130      3140      3150      3160      3170      3180      3190      3200
         *    *    *    *    *    *    *    *    *    *    *    *    *    *    *    *    *    *    *    *
                                           3139 HhaI
                                          3138 FspI
                                      3134 AciI
                    3116 MseI      3131 RsaI                3156 MluCI
                    |              |  |   ||                |
3101 atatatatgtctttttaatggggtgtatagtaccgctgcgcatagtttttctgtaatttacaacagtgctattttctggtagttcttcggagtgtgttg 3200
       I  Y  V  F  F  N  G  V  Y  S  T  A  A  H  S  F  S  V  I  Y  N  S  A  I  F  W  *  F  F  G  V  C  C


            3210      3220      3230      3240      3250      3260      3270      3280      3290      3300
         *    *    *    *    *    *    *    *    *    *    *    *    *    *    *    *    *    *    *    *
                                                                           3271 RsaI
                3212 ApoI                                          3263 HpaII
              3210 MseI                  3237 DpnI    3250 RsaI   3262 NgoMIV
         3205 MluCI               3230 MluCI         3249 TatI    3262 NaeI              3285 BfaI
       3203 MseI 3213 MluCI      3229 ApoI           3249 BsrGI   3262 Cac8I     3277 AluI      3292 MluCI
       | |    | ||               ||      |           ||           ||       |     |       |      |
3201 ctttaattattaaatttatataatcaatgaatttgggatcgtcggttttgtacaatatgttgccggcatagtacgcagcttcttctagttcaattacacc 3300
      F  N  Y  *  I  Y  I  I  N  E  F  G  I  V  G  F  V  Q  Y  V  A  G  I  V  R  S  F  F  *  F  N  Y  T


            3310      3320      3330      3340      3350      3360      3370      3380      3390      3400
         *    *    *    *    *    *    *    *    *    *    *    *    *    *    *    *    *    *    *    *
                        3320 MseI
                   3315 HpaII
                  3314 BsaWI                   3343 RsaI 3353 MseI    3366 Hpy8I                   3395 Cac8I
                  ||    |                      |         |            |                            |
3301 atttttagcagcaccggattaacataactttccaaaatgttgtacgaaccgttaaacaaaaacagttcacctcccttttctatactattgtctgcgagc 3400
     I  F  *  Q  H  R  I  N  I  T  F  Q  N  V  V  R  T  V  K  Q  K  Q  F  T  S  L  F  Y  T  I  V  C  E  Q
```

```
          3410      3420      3430      3440      3450      3460      3470      3480      3490      3500
       *    *    *    *    *    *    *    *    *    *    *    *    *    *    *    *    *    *    *    *
                                                                                                  3497 HaeIII
                                                                                                 3496 EagI
                                                                                                3495 HpaII
                                                                                               3494 NgoMIV
                                                                                               3494 NaeI
                                                                                               3494 Cac8I
                                                                                              3493 HaeIII
            3413 MseI              3436 BsmBI                                                 3493 FseI
             |                      |                                                         |||||
3401 agttgtttgttgttaaaaataacagccattgtaatgagacgcacaaactaatatcacaaactggaaatgtctatcaatatatagttgctgatggccggcc 3500
       L  F  V  V  K  N  N  S  H  C  N  E  T  H  K  L  I  S  Q  T  G  N  V  Y  Q  Y  I  V  A  D  G  R  P


          3510      3520      3530      3540      3550      3560      3570      3580      3590      3600
       *    *    *    *    *    *    *    *    *    *    *    *    *    *    *    *    *    *    *    *
                                                                             3567 MseI
                                                                            3566 AseI
        3506 BsmBI                                                    3560 BfaI           3579 MluCI
         |                                                             |     ||            |
3501 gtaatgagacgcacaaactaatatcacaaactggaaatgtctatcaatatatagttgctctagttattaatagtaatcaattacggggtcattagttcat 3600
      *  *  D  A  Q  T  N  I  T  N  W  K  C  L  S  I  Y  S  C  S  S  Y  *  *  *  S  I  T  G  S  L  V  H


          3610      3620      3630      3640      3650      3660      3670      3680      3690      3700
       *    *    *    *    *    *    *    *    *    *    *    *    *    *    *    *    *    *    *    *
                                                    3646 AciI                                3681 ZraI
                  3619 BstUI                      3644 Cac8I                                 3681 AatII   3694 HpyCH4IV
                 3618 AciI                       3643 HaeIII    3658 AciI     3672 AciI 3682 HpyCH4IV
                 ||                              || |           |             |        ||            |
3601 agcccatatatggagttccgcgttacataacttacggtaaatggcccgcctggctgaccgcccaacgacccccgcccattgacgtcaataatgacgtatg 3700
     S  P  Y  M  E  F  R  V  T  *  L  T  V  N  G  P  P  G  *  P  P  N  D  P  R  P  L  T  S  I  M  T  Y  V


          3710      3720      3730      3740      3750      3760      3770      3780      3790      3800
       *    *    *    *    *    *    *    *    *    *    *    *    *    *    *    *    *    *    *    *
                                   3735 HpyCH4IV
                                  3734 ZraI                                  3778 RsaI
                                  3734 AatII              3759 Hpy8I        3777 TatI       3793 NdeI
                                  ||                       |                 ||              |
3701 ttcccatagtaacgccaatagggactttccattgacgtcaatgggtggagtatttacggtaaactgcccacttggcagtacatcaagtgtatcatatgcc 3800
       P  I  V  T  P  I  G  T  F  H  *  R  Q  W  V  E  Y  L  R  *  T  A  H  L  A  V  H  Q  V  Y  H  M  P


          3810      3820      3830      3840      3850      3860      3870      3880      3890      3900
       *    *    *    *    *    *    *    *    *    *    *    *    *    *    *    *    *    *    *    *
                                                                                                   3899 HpyCH4IV
                                                                                                  3898 SnaBI
                                                                                                  3898 BsaAI
                 3818 HpyCH4IV        3839 AciI            3861 NlaIII
                3817 ZraI           3837 Cac8I           3858 RsaI                          3891 RsaI
       3803 RsaI    3817 AatII         3836 HaeIII          3857 TatI                         3890 TatI
        |           ||                 || |                 ||  |                             ||      ||
3801 aagtacgccccctattgacgtcaatgacggtaaatggcccgcctggcattatgcccagtacatgaccttatgggactttcctacttggcagtacatctac 3900
      S  T  P  P  I  D  V  N  D  G  K  W  P  A  W  H  Y  A  Q  Y  M  T  L  W  D  F  P  T  W  Q  Y  I  Y


          3910      3920      3930      3940      3950      3960      3970      3980      3990      4000
       *    *    *    *    *    *    *    *    *    *    *    *    *    *    *    *    *    *    *    *
                             3930 TaqI
                        3925 NlaIII
                      3923 HpyCH4V
                     3922 NsiI
                    3921 NlaIII           3944 HpyCH4IV                              3988 MluCI
                    ||| |    |            |                                          |
3901 gtattagtcatcgctattaccatgcatggtcgaggtgagccccacgttctgcttcactctccccatctcccccccctccccacccccaattttgtattta 4000
     V  L  V  I  A  I  T  M  H  G  R  G  E  P  H  V  L  L  H  S  P  H  L  P  P  L  P  T  P  N  F  V  F  I


          4010      4020      4030      4040      4050      4060      4070      4080      4090      4100
       *    *    *    *    *    *    *    *    *    *    *    *    *    *    *    *    *    *    *    *
                                                               4055 HhaI
                                                              4054 BstUI
                                                             4053 HhaI
                                                             4053 Cac8I
                                                             4053 BssHII
                                                            4052 BstUI
                                                           4051 HhaI
            4011 MluCI                                     4051 Cac8I 4062 AciI 4072 AciI       4089 AciI
          4009 MseI   4021 HpyCH4V 4034 AciI               4051 BssHII     4067 AciI       4084 AciI      4099 AciI
           | |         |            |                      |||||      |    |    |          |    |         |
4001 tttatttttaattattttgtgcagcgatgggggcgggggggggggggggcgcgcgccaggcggggcggggcggggcgaggggcggggcggggcgaggc 4100
       Y  F  L  I  I  L  C  S  D  G  G  G  G  G  G  G  A  R  A  R  R  G  G  A  G  R  G  A  G  R  G  E  A


          4110      4120      4130      4140      4150      4160      4170      4180      4190      4200
       *    *    *    *    *    *    *    *    *    *    *    *    *    *    *    *    *    *    *    *
                                4133 HhaI                                                         4198 AciI
```

```
                          4132 BstUI                                                    4197 BstUI
                         4131 HhaI                                                     4196 HhaI
                         4131 Cac8I                            4171 AciI              4195 BstUI
                         4131 BssHII                        4168 AciI                4194 HhaI
           4112 AciI       4128 AciI                     4165 AciI  4176 HaeIII      4194 Cac8I
        4109 AciI        4126 BsrBI                   4162 AciI   4174 AciI          4194 BssHII
        |  |             | |  |||                        |  |  |  |  | |             |||||
4101 ggagaggtgcggcggcagccaatcagagcggcgcgctccgaaagtttccttttatggcgaggcggcggcggcggcggccctataaaaagcgaagcgcgcg 4200
      E  R  C  G  G  S  Q  S  E  R  R  A  P  K  V  S  F  Y  G  E  A  A  A  A  A  A  L  *  K  A  K  R  A

          4210      4220      4230      4240      4250      4260      4270      4280      4290      4300
        *    *    *    *    *    *    *    *    *    *    *    *    *    *    *    *    *    *    *    *
                  4219 HhaI                              4265 AciI
                 4218 BstUI                 4248 AciI   4263 Cac8I
    4205 AciI   4217 HhaI                 4245 AciI    4258 HhaI
 4201 Cac8I     4217 Cac8I            4240 BsrBI      4257 BstUI   4270 HpaII     4285 BstUI
 4201 AciI      4217 BssHII           4240 AciI  4251 AciI 4261 AciI              4284 AciI
 |   |          |||                   |     |  |  |     ||  | | |    |             ||
4201 gcgggcgggagtcgctgcgcgctgccttcgccccgtgccccgctccgccgccgcctcgcgccgcccgccccggctctgactgaccgcgttactcccacag 4300
     A  G  G  S  R  C  A  L  P  S  P  R  A  P  L  R  R  R  L  A  P  P  A  P  A  L  T  D  R  V  T  P  T  G

          4310      4320      4330      4340      4350      4360      4370      4380      4390      4400
        *    *    *    *    *    *    *    *    *    *    *    *    *    *    *    *    *    *    *    *
        4309 AciI                     4342 HhaI
    4305 Cac8I                       4341 HaeII
    4305 AciI                        4341 AfeI
  4303 BsrBI   4316 HaeIII         4337 MluCI    4351 MseI
   | |   |      |                  |   ||        |
4301 gtgagcgggcgggacggcccttctcctccgggctgtaattagcgcttggtttaatgacggcttgtttcttttctgtggctgcgtgaaagccttgaggggc 4400
       E  R  A  G  R  P  F  S  F  G  L  *  L  A  L  G  L  M  T  A  C  F  F  S  V  A  A  *  K  P  *  G  A

          4410      4420      4430      4440      4450      4460      4470      4480      4490      4500
        *    *    *    *    *    *    *    *    *    *    *    *    *    *    *    *    *    *    *    *
                                            4445 AciI
        4409 HaeIII        4428 AciI       4444 BstUI
       4408 ApaI         4426 BsrBI       4443 SacII
  4402 HpaII       4419 AciI    4432 AvaI  4443 AciI                             4482 AciI
  |      ||        |       | |    |        |||                                    |
4401 tccgggagggccctttgtgcggggggagcggctcggggctgtccgcggggggacggctgccttcgggggggacggggcagggcggggttcggcttctggc 4500
      P  G  G  P  F  V  R  G  E  R  L  G  A  V  R  G  G  T  A  A  F  G  G  D  G  A  G  R  G  S  A  S  G

          4510      4520      4530      4540      4550      4560      4570      4580      4590      4600
        *    *    *    *    *    *    *    *    *    *    *    *    *    *    *    *    *    *    *    *
               4516 BfaI
              4515 XbaI
         4510 AciI                   4539 NlaIII
      4507 HpaII             4533 NlaIII                 4561 AluI   4573 HpyCH4IV
       |  |    ||            |     |                      |           |
4501 gtgtgaccggcggctctagagcctctgctaaccatgttcatgccttcttctttttcctacagctcctgggcaacgtgctggttattgtgctgtctcatca 4600
     V  *  P  A  A  L  E  P  L  L  T  M  F  M  P  S  S  F  S  Y  S  S  W  A  T  C  W  L  L  C  C  L  I  I

          4610      4620      4630      4640      4650      4660      4670      4680      4690      4700
        *    *    *    *    *    *    *    *    *    *    *    *    *    *    *    *    *    *    *    *
                             4630 MseI                    4660 AciI
               4617 DpnI   4629 AseI                    4658 BsrBI      4675 HpaII 4686 HpaII 4697 MluCI
          4612 MluCI      4628 MluCI              4652 MluCI      4669 MluCI  4681 MseI     4695 MseI
          |    |          |||                     |     | |       |     |     |    |        | |
4601 ttttggcaaagaattggatcggaccgaaatTAATACGACTCACTATAGGGGAATTGTGAGCGGATAACAATTCCCcggagttaatccgggacctttaatt 4700
       L  A  K  N  W  I  G  P  K  L  I  R  L  T  I  G  E  L  *  A  D  N  N  S  P  E  L  I  R  D  L  *  F
                              >>>>>>>>>>>>>>>>>>>>
                              T7 promoter sequencing primer

          4710      4720      4730      4740      4750      4760      4770      4780      4790      4800
        *    *    *    *    *    *    *    *    *    *    *    *    *    *    *    *    *    *    *    *
                                                            4761 MseI
                                                          4759 MluCI
                                                        4757 PacI
                                4733 MluCI              4757 MseI
                      4724 MseI                        4756 AseI             4779 MluCI
                       |        |                      || | |                 |
4701 caacccaacacaatatattatagttaaataagaattattatcaaatcatttgtatattaattaaaatactatactgtaaattacattttatttacaatca 4800
      N  P  T  Q  Y  I  I  V  K  *  E  L  L  S  N  H  L  Y  I  N  *  N  T  I  L  *  I  T  F  Y  L  Q  S

          4810      4820      4830      4840      4850      4860      4870      4880      4890      4900
        *    *    *    *    *    *    *    *    *    *    *    *    *    *    *    *    *    *    *    *
           4813 NlaIII                     4846 NlaIII      4863 Hpy8I
          4812 NcoI                    4842 AciI   4854 TaqI                 4881 MseI
          ||                           |   |       |        |                 |
4801 aaggagatataccATGgcacaccatcaccaccatcacagcagcggcATGAAAATCGAAGAAGGTAAACTGGTAATCTGGATTAACGGCGATAAAGGCTAT 4900
     K  E  I  Y  H  G  T  P  S  P  P  S  Q  Q  R  H  E  N  R  R  R  *  T  G  N  L  D  *  R  R  *  R  L  *
                 >>>>>>>>>>>>>>>>>>>>>>>>>   >>>>>>>>>>>>>>>>>>>>>>>>>>>>>>>>>>>>>>>>>>>>>>>>>>>>>>>>>>>>
                 His6                        MBP
```

```
        4910      4920      4930      4940      4950      4960      4970      4980      4990      5000
    *    *    *    *    *    *    *    *    *    *    *    *    *    *    *    *    *    *    *    *
                                                    4947 MseI
                                  4929 TaqI       4945 MluCI                4967 HpaII
                                4926 MluCI      4941 HpaII                4966 BspEI          4986 MluCI
    4904 BsaI                  4925 ApoI       4940 BsaWI                 4966 BsaWI         4985 ApoI      5000 AciI
     |                         ||  |           ||   | |                   ||                 ||             |
4901 AACGGTCTCGCTGAAGTCGGTAAGAAATTCGAGAAAGATACCGGAATTAAAGTCACCGTTGAGCATCCGGATAAACTGGAAGAGAAATTCCCACAGGTTG 5000
       R  S  R  *  S  R  *  E  I  R  E  R  Y  R  N  *  S  H  R  *  A  S  G  *  T  G  R  E  I  P  T  G  C
     >>>>>>>>>>>>>>>>>>>>>>>>>>>>>>>>>>>>>>>>>>>>>>>>>>>>>>>>>>>>>>>>>>>>>>>>>>>>>>>>>>>>>>>>>>>>>>>>>>>>
     MBP

        5010      5020      5030      5040      5050      5060      5070      5080      5090      5100
    *    *    *    *    *    *    *    *    *    *    *    *    *    *    *    *    *    *    *    *
               5015 HaeIII                  5044 AciI                 5069 HaeIII         5090 HpaII
               |                            |                         |                   |
5001 CGGCAACTGGCGATGGCCCTGACATTATCTTCTGGGCACACGACCGCTTTGGTGGCTACGCTCAATCTGGCCTGTTGGCTGAAATCACCCCGGACAAAGC 5100
      G  N  W  R  W  P  *  H  Y  L  L  G  T  R  P  L  W  W  L  R  S  I  W  P  V  G  *  N  H  P  G  Q  S
     >>>>>>>>>>>>>>>>>>>>>>>>>>>>>>>>>>>>>>>>>>>>>>>>>>>>>>>>>>>>>>>>>>>>>>>>>>>>>>>>>>>>>>>>>>>>>>>>>>>>
     MBP

        5110      5120      5130      5140      5150      5160      5170      5180      5190      5200
    *    *    *    *    *    *    *    *    *    *    *    *    *    *    *    *    *    *    *    *
                                        5140 HpyCH4IV
                                      5138 RsaI       5154 AluI       5170 DpnI
            5112 AluI 5122 Hpy8I     5137 BsiWI     5151 Cac8I      5169 PvuI                  5197 PsiI
                 |         |              || |          |  |            ||                        |
5101 GTTCCAGGACAAGCTGTATCCGTTTACCTGGGATGCCGTACGTTACAACGGCAAGCTGATTGCTTACCCGATCGCTGTTGAAGCGTTATCGCTGATTTAT 5200
     V  P  G  Q  A  V  S  V  Y  L  G  C  R  T  L  Q  R  Q  A  D  C  L  P  D  R  C  *  S  V  I  A  D  L  *
     >>>>>>>>>>>>>>>>>>>>>>>>>>>>>>>>>>>>>>>>>>>>>>>>>>>>>>>>>>>>>>>>>>>>>>>>>>>>>>>>>>>>>>>>>>>>>>>>>>>>
     MBP

        5210      5220      5230      5240      5250      5260      5270      5280      5290      5300
    *    *    *    *    *    *    *    *    *    *    *    *    *    *    *    *    *    *    *    *
                                                       5249 HhaI                           5285 HhaI
                                                      5248 HaeII                          5284 BstUI
      5207 DpnI                                   5246 HpaII                            5283 HhaI
     5206 BstYI                               5242 DpnI                                 5283 Cac8I
     5206 BglII      5222 AciI               5241 BstYI                                 5283 BssHII
      ||               |                      ||   | ||                                  |||
5201 AACAAAGATCTGCTGCCGAACCCGCCAAAAACCTGGGAAGAGATCCCGGCGCTGGATAAAGAACTGAAAGCGAAAGGTAAGAGCGCGCTGATGTTCAACC 5300
       Q  R  S  A  A  E  P  A  K  N  L  G  R  D  P  G  A  G  *  R  T  E  S  E  R  *  E  R  A  D  V  Q  P
     >>>>>>>>>>>>>>>>>>>>>>>>>>>>>>>>>>>>>>>>>>>>>>>>>>>>>>>>>>>>>>>>>>>>>>>>>>>>>>>>>>>>>>>>>>>>>>>>>>>>
     MBP

        5310      5320      5330      5340      5350      5360      5370      5380      5390      5400
    *    *    *    *    *    *    *    *    *    *    *    *    *    *    *    *    *    *    *    *
                                                                                     5388 HpyCH4IV
          5311 RsaI    5324 AciI                                                5382 MseI
     5301 HpyCH4V     5322 HaeIII                                          5374 RsaI    5387 BmgBI
     |         |         | |                                                   |       ||
5301 TGCAAGAACCGTACTTCACCTGGCCGCTGATTGCTGCTGACGGGGGTTATGCGTTCAAGTATGAAAACGGCAAGTACGACATTAAAGACGTGGGCGTGGA 5400
      A  R  T  V  L  H  L  A  A  D  C  C  *  R  G  L  C  V  Q  V  *  K  R  Q  V  R  H  *  R  R  G  R  G
     >>>>>>>>>>>>>>>>>>>>>>>>>>>>>>>>>>>>>>>>>>>>>>>>>>>>>>>>>>>>>>>>>>>>>>>>>>>>>>>>>>>>>>>>>>>>>>>>>>>>
     MBP

        5410      5420      5430      5440      5450      5460      5470      5480      5490      5500
    *    *    *    *    *    *    *    *    *    *    *    *    *    *    *    *    *    *    *    *
         5410 BstUI                        5445 MseI
        5409 HhaI                 5435 Hpy8I          5464 HpyCH4V                       5499 MseI
    5405 Cac8I 5417 AciI          5435 HincII     5458 NlaIII                   5491 AluI
     |   ||     |                 |         |      |     |                       |       |
5401 TAACGCTGGCGCGAAAGCGGGTCTGACCTTCCTGGTTGACCTGATTAAAAACAAACACATGAATGCAGACACCGATTACTCCATCGCAGAAGCTGCCTTT 5500
     *  R  W  R  E  S  G  S  D  L  P  G  *  P  D  *  K  Q  T  H  E  C  R  H  R  L  L  H  R  R  S  C  L  *
     >>>>>>>>>>>>>>>>>>>>>>>>>>>>>>>>>>>>>>>>>>>>>>>>>>>>>>>>>>>>>>>>>>>>>>>>>>>>>>>>>>>>>>>>>>>>>>>>>>>>
     MBP

        5510      5520      5530      5540      5550      5560      5570      5580      5590      5600
    *    *    *    *    *    *    *    *    *    *    *    *    *    *    *    *    *    *    *    *
                                        5541 NlaIII
                 5518 BsaBI   5531 HaeIII          5553 TaqI        5570 MluCI     5585 RsaI
                  |            |         |           |                |              |
5501 AATAAAGGCGAAACAGCGATGACCATCAACGGCCCGTGGGCATGGTCCAACATCGACACCAGCAAAGTGAATTATGGTGTAACGGTACTGCCGACCTTCA 5600
       *  R  R  N  S  D  D  H  Q  R  P  V  G  M  V  Q  H  R  H  Q  Q  S  E  L  W  C  N  G  T  A  D  L  Q
     >>>>>>>>>>>>>>>>>>>>>>>>>>>>>>>>>>>>>>>>>>>>>>>>>>>>>>>>>>>>>>>>>>>>>>>>>>>>>>>>>>>>>>>>>>>>>>>>>>>>
     MBP

        5610      5620      5630      5640      5650      5660      5670      5680      5690      5700
    *    *    *    *    *    *    *    *    *    *    *    *    *    *    *    *    *    *    *    *
    5604 Hpy8I                        5637 HhaI     5652 AciI         5671 Cac8I
    5604 HincII           5628 Cac8I         5646 MseI               5670 AluI          5688 TaqI
```

```
         |                                 |            |            |       |                        ||                      |
5601 AGGGTCAACCATCCAAACCGTTCGTTGGCGTGCTGAGCGCAGGTATTAACGCCGCCAGTCCGAACAAAGAGCTGGCAAAAGAGTTCCTCGAAAACTATCT 5700
      G  S  T  I  Q  T  V  R  W  R  A  E  R  R  Y  *  R  R  Q  S  E  Q  R  A  G  K  R  V  P  R  K  L  S
     >>>>>>>>>>>>>>>>>>>>>>>>>>>>>>>>>>>>>>>>>>>>>>>>>>>>>>>>>>>>>>>>>>>>>>>>>>>>>>>>>>>>>>>>>>>>>>>>>>>
     MBP

             5710      5720      5730      5740      5750      5760      5770      5780      5790      5800
         *    *    *    *    *    *    *    *    *    *    *    *    *    *    *    *    *    *    *    *
                                                                                                 5794 HpyCH4IV
                                                                 5756 HhaI                      5793 BsaAI
                                   5727 MseI                    5755 HaeII                  5789 DpnI
                               5723 AciI          5741 AciI     5755 AfeI                  5788 BstYI
                                      |   |              |             ||                              ||   ||
5701 GCTGACTGATGAAGGTCTGGAAGCGGTTAATAAAGACAAACCGCTGGGTGCCGTAGCGCTGAAGTCTTACGAGGAAGAGTTGGCGAAAGATCCACGTATT 5800
     A  D  *  *  R  S  G  S  G  *  *  R  Q  T  A  G  C  R  S  A  E  V  L  R  G  R  V  G  E  R  S  T  Y  C
     >>>>>>>>>>>>>>>>>>>>>>>>>>>>>>>>>>>>>>>>>>>>>>>>>>>>>>>>>>>>>>>>>>>>>>>>>>>>>>>>>>>>>>>>>>>>>>>>>>>
     MBP

             5810      5820      5830      5840      5850      5860      5870      5880      5890      5900
         *    *    *    *    *    *    *    *    *    *    *    *    *    *    *    *    *    *    *    *
                                                                                          5885 AciI    5898 AciI
      5802 AciI                                5836 NlaIII  5849 AciI 5859 AciI        5880 RsaI 5890 DpnI 5900 Cac8I
      |                                         |            |         |                 |    |    |       | |
5801 GCCGCCACTATGGAAAACGCCCAGAAAGGTGAAATCATGCCGAACATCCCGCAGATGTCCGCTTTCTGGTATGCCGTGCGTACTGCGGTGATCAACGCCG 5900
       R  H  Y  G  K  R  P  E  R  *  N  H  A  E  H  P  A  D  V  R  F  L  V  C  R  A  Y  C  G  D  Q  R  R
     >>>>>>>>>>>>>>>>>>>>>>>>>>>>>>>>>>>>>>>>>>>>>>>>>>>>>>>>>>>>>>>>>>>>>>>>>>>>>>>>>>>>>>>>>>>>>>>>>>>
     MBP

             5910      5920      5930      5940      5950      5960      5970      5980      5990      6000
         *    *    *    *    *    *    *    *    *    *    *    *    *    *    *    *    *    *    *    *
                                                           5946 BfaI                    5976 RsaI
                                                    5939 HhaI                          5975 KpnI                6000 HaeIII
       5904 AciI      5919 TaqI                    5938 BstUI    5952 AciI             5975 Acc65I             5999 ApaI
        |               |                            ||       |      |                      ||                       ||
5901 CCAGCGGTCGTCAGACTGTCGATGAAGCCCTGAAAGACGCGCAGACTagcagcggtctggaagttctgtttcagGGTACCctggaagttctgttccaggg 6000
      Q  R  S  S  D  C  R  *  S  P  E  R  R  A  D  *  Q  R  S  G  S  S  V  S  G  Y  P  G  S  S  V  P  G
     >>>>>>>>>>>>>>>>>>>>>>>>>>>>>>>>>>>>>>>>>>>>>>>
     MBP

             6010      6020      6030      6040      6050      6060      6070      6080      6090      6100
         *    *    *    *    *    *    *    *    *    *    *    *    *    *    *    *    *    *    *    *
        6006 TaqI                        6039 HpaII                                      6083 HaeIII
       6005 XhoI                     6034 Hpy8I                       6063 AluI     6077 Hpy8I
       6005 AvaI                 6030 AluI                        6060 TaqI      6075 HpyCH4IV
        ||                         |   |    |                        |  |           | |     |
6001 gcccCTCGAGatggtgagcaagggcgaggagctgttcaccggggtggtgcccatcctggtcgagctggacggcgacgtaaacggccacaagttcagcgtg 6100
     A  P  R  D  G  E  Q  G  R  G  A  V  H  R  G  G  A  H  P  G  R  A  G  R  R  R  K  R  P  Q  V  Q  R  V
         >>>>>>>>>>>>>>>>>>>>>>>>>>>>>>>>>>>>>>>>>>>>>>>>>>>>>>>>>>>>>>>>>>>>>>>>>>>>>>>>>>>>>>>>>>>>>>>>>>
         XhoI  eYFP-ORF

             6110      6120      6130      6140      6150      6160      6170      6180      6190      6200
         *    *    *    *    *    *    *    *    *    *    *    *    *    *    *    *    *    *    *    *
                                                                         6165 Cac8I
                                        6135 AluI                     6162 HpaII                6191 BssSI
      6102 HpaII                     6132 Cac8I            6155 HpyCH4V 6168 AluI      6183 HaeIII
      |                                |  |                  |      |  |  |              |       |
6101 tccggcgagggcgagggcgatgccacctacggcaagctgaccctgaagttcatctgcaccaccggcaagctgcccgtgccctggcccaccctcgtgacca 6200
       R  R  G  R  G  R  C  H  L  R  Q  A  D  P  E  V  H  L  H  H  R  Q  A  A  R  A  L  A  H  P  R  D  H
     >>>>>>>>>>>>>>>>>>>>>>>>>>>>>>>>>>>>>>>>>>>>>>>>>>>>>>>>>>>>>>>>>>>>>>>>>>>>>>>>>>>>>>>>>>>>>>>>>>>
     eYFP-ORF

             6210      6220      6230      6240      6250      6260      6270      6280      6290      6300
         *    *    *    *    *    *    *    *    *    *    *    *    *    *    *    *    *    *    *    *
                 6216 HpyCH4V
               6215 PstI
             6213 Cac8I      6229 AciI                                    6274 NlaIII              6298 HhaI
            6212 HaeIII    6227 Cac8I       6244 NlaIII               6270 AciI         6288 HpyCH4IV
              || ||           | |              |                          |   |             |         |
6201 ccttcggctacggcctgcagtgcttcgcccgctaccccgaccacatgaagcagcacgacttcttcaagtccgccatgcccgaaggctacgtccaggagcg 6300
      L  R  L  R  P  A  V  L  R  P  L  P  R  P  H  E  A  A  R  L  L  Q  V  R  H  A  R  R  L  R  P  G  A
     >>>>>>>>>>>>>>>>>>>>>>>>>>>>>>>>>>>>>>>>>>>>>>>>>>>>>>>>>>>>>>>>>>>>>>>>>>>>>>>>>>>>>>>>>>>>>>>>>>>
     eYFP-ORF

             6310      6320      6330      6340      6350      6360      6370      6380      6390      6400
         *    *    *    *    *    *    *    *    *    *    *    *    *    *    *    *    *    *    *    *
                                                    6339 HhaI                              6381 TaqI
                                                   6338 BstUI                         6376 AciI
                                                  6337 AciI        6354 TaqI        6371 Hpy8I   6384 AluI   6396 TaqI
                                                  |||                |                 |    |    |  |           |
6301 caccatcttcttcaaggacgacggcaactacaagacccgcgccgaggtgaagttcgagggcgacaccctggtgaaccgcatcgagctgaagggcatcgac 6400
     H  H  L  L  Q  G  R  R  Q  L  Q  D  P  R  R  G  E  V  R  G  R  H  P  G  E  P  H  R  A  E  G  H  R  L
     >>>>>>>>>>>>>>>>>>>>>>>>>>>>>>>>>>>>>>>>>>>>>>>>>>>>>>>>>>>>>>>>>>>>>>>>>>>>>>>>>>>>>>>>>>>>>>>>>>>
     eYFP-ORF
```

```
             6410      6420      6430      6440      6450      6460      6470      6480      6490      6500
         *    *    *    *    *    *    *    *    *    *    *    *    *    *    *    *    *    *    *    *
                                            6439 RsaI                         6472 HaeIII
                                           6438 TatI                       6469 NlaIII
                                    6432 AluI                  6459 HpyCH4IV                           6500 Hpy8I
                                    |     ||                   |         |  |                          |
6401 ttcaaggaggacggcaacatcctggggcacaagctggagtacaactacaacagccacaacgtctatatcatggccgacaagcagaagaacggcatcaagg 6500
       Q  G  G  R  Q  H  P  G  A  Q  A  G  V  Q  L  Q  Q  P  Q  R  L  Y  H  G  R  Q  A  E  E  R  H  Q  G
     >>>>>>>>>>>>>>>>>>>>>>>>>>>>>>>>>>>>>>>>>>>>>>>>>>>>>>>>>>>>>>>>>>>>>>>>>>>>>>>>>>>>>>>>>>>>>>>>>>
     eYFP-ORF


             6510      6520      6530      6540      6550      6560      6570      6580      6590      6600
         *    *    *    *    *    *    *    *    *    *    *    *    *    *    *    *    *    *    *    *
                                                 6544 Cac8I
                                                6543 AluI
             6514 AciI                         6540 HpyCH4V
          6511 DpnI
         6510 BstYI     6525 TaqI   6537 Cac8I                                      6584 HaeIII
          ||  |          |           |  |  ||                                        |
6501 tgaacttcaagatccgccacaacatcgaggacggcagcgtgcagctcgccgaccactaccagcagaacacccccatcggcgacggccccgtgctgctgCC 6600
      E  L  Q  D  P  P  Q  H  R  G  R  Q  R  A  A  R  R  P  L  P  A  E  H  P  H  R  R  R  P  R  A  A  A
     >>>>>>>>>>>>>>>>>>>>>>>>>>>>>>>>>>>>>>>>>>>>>>>>>>>>>>>>>>>>>>>>>>>>>>>>>>>>>>>>>>>>>>>>>>>>>>>>>>
     eYFP-ORF


             6610      6620      6630      6640      6650      6660      6670      6680      6690      6700
         *    *    *    *    *    *    *    *    *    *    *    *    *    *    *    *    *    *    *    *
                                                                6664 NlaIII
                                                           6659 DpnI                        6693 HpaII
                                                        6656 BstUI                       6690 AciI
                 6617 AluI 6627 AciI                   6655 HhaI                      6687 AciI 6697 DpnI
                    |         |                          ||  |    |                      |  |  |  |
6601 CGACAACCACTACCTGAGCtaccagtccgccctgagcaaagaccccaacgagaagcgcgatcacatggtcctgctggagttcgtgaccgccgccgggatc 6700
     R  Q  P  L  P  E  L  P  V  R  P  E  Q  R  P  Q  R  E  A  R  S  H  G  P  A  G  V  R  D  R  R  R  D  H
     >>>>>>>>>>>>>>>>>>>>>>>>>>>>>>>>>>>>>>>>>>>>>>>>>>>>>>>>>>>>>>>>>>>>>>>>>>>>>>>>>>>>>>>>>>>>>>>>>>
     eYFP-ORF


             6710      6720      6730      6740      6750      6760      6770      6780      6790      6800
         *    *    *    *    *    *    *    *    *    *    *    *    *    *    *    *    *    *    *    *
                                    6729 TaqI
                                   6728 XhoI
                                 6726 AluI 6736 HaeIII
                           6720 TatI      6735 EagI
                           6720 BsrGI    6734 NotI
                        6717 AluI  6728 AvaI 6738 AciI                    6771 AluI
                6709 NlaIII 6721 RsaI    6734 AciI                6762 Cac8I   6775 DpnI 6785 BbsI
                 |       |  ||      | ||     ||| |                    |        |   |         |
6701 actctcggcatggacgagctgtacaagCTCGAGGCGGCCGCaGAAAACTTGTATTTCCAGGGCgagcagaagctgatctcagaggaagacctgatgagcc 6800
       S  R  H  G  R  A  V  Q  A  R  G  G  R  R  K  L  V  F  P  G  R  A  E  A  D  L  R  G  R  P  D  E  P
     >>>>>>>>>>>>>>>>>>>>>>>>>>>>>>>>>>>>>>>>>>                                                 >>>
     eYFP-ORF                    XhoI  NotI                                                     START codon
                                                                                                   >>>>
                                                                                                   1st part


             6810      6820      6830      6840      6850      6860      6870      6880      6890      6900
         *    *    *    *    *    *    *    *    *    *    *    *    *    *    *    *    *    *    *    *
                                                               6858 NcoI
                                                       6850 AciI
                                 6830 BbsI           6848 BsrBI 6859 NlaIII
                                   |                   | |        ||
6801 ctgaaaagtcccaagaggagagcccagaagaagacacagagagaacagagcggaagcccatggtcaaagatgccttcaaagacatttccatatacttcac 6900
      *  K  V  P  R  G  E  P  R  R  R  H  R  E  N  R  A  E  A  H  G  Q  R  C  L  Q  R  H  F  H  I  L  H
     >>>>>>>>>>>>>>>>>>>>>>>>>>>>>>>>>>>>>>>>>>>>>>>>>>>>>>>>>>>>>>>>>>>>>>>>>>>>>>>>>>>>>>>>>>>>>>>>>>
     1st part


             6910      6920      6930      6940      6950      6960      6970      6980      6990      7000
         *    *    *    *    *    *    *    *    *    *    *    *    *    *    *    *    *    *    *    *
                                                                        6970 HpyCH4V    6986 BsaI     7000 TaqI
                                                                         |                |             |
6901 caaggaagaatgggcagagatgggagactgggagaaaactcgctataggaatgtgaaaaggaactataatgcactgattactataggtctcagagccact 7000
     Q  G  R  M  G  R  D  G  R  L  G  E  N  S  L  *  E  C  E  K  E  L  *  C  T  D  Y  Y  R  S  Q  S  H  S
     >>>>>>>>>>>>>>>>>>>>>>>>>>>>>>>>>>>>>>>>>>>>>>>>>>>>>>>>>>>>>>>>>>>>>>>>>>>>>>>>>>>>>>>>>>>>>>>>>>
     1st part


             7010      7020      7030      7040      7050      7060      7070      7080      7090      7100
         *    *    *    *    *    *    *    *    *    *    *    *    *    *    *    *    *    *    *    *
               7012 NlaIII       7030 HaeIII                                      7083 BfaI
          7006 AluI            7027 Cac8I                                        7082 AvrII
           |     |               |  |                                             ||
7001 cgaccagctttcatgtgtcaccgaaggcaggccatcaaactccaggtggatgacacagaagattctgatgaagaatggacccctaggcagcaagtcaaac 7100
       T  S  F  H  V  S  P  K  A  G  H  Q  T  P  G  G  *  H  R  R  F  *  *  R  M  D  P  *  A  A  S  Q  T
     >>>>>>>>>>>>>>>>>>>>>>>>>>>>>>>>>>>>>>>>>>>>>>>>>>>>>>>>>>>>>>>>>>>>>>>>>>>>>>>>>>>>>>>>>>>>>>>>>>
     1st part
```

```
        7110      7120      7130      7140      7150      7160      7170      7180      7190      7200
           *    *    *    *    *    *    *    *    *    *    *    *    *    *    *    *    *    *    *    *
                  7115 MseI
                 7114 AflII
             7111 HaeIII         7131 Hpy8I                                    7176 BfaI   7188 MluCI
             |   ||              |                                             |           |
7101 ctccttggatggccttaagagtggaacagcgtaaacaccagaagggaatgcccaaggcgtcattcagtaatgaatctagtttgaaagaattgtcaagaac 7200
      S  L  D  G  L  K  S  G  T  A  *  T  P  E  G  N  A  Q  G  V  I  Q  *  *  I  *  F  E  R  I  V  K  N
     >>>>>>>>>>>>>>>>>>>>>>>>>>>>>>>>>>>>>>>>>>>>>>>>>>>>>>>>>>>>>>>>>>>>>>>>>>>>>>>>>>>>>>>>>>>>>>>>>>>>
     1st part

        7210      7220      7230      7240      7250      7260      7270      7280      7290      7300
           *    *    *    *    *    *    *    *    *    *    *    *    *    *    *    *    *    *    *    *
        7205 MluCI
       7204 ApoI   7216 HpyCH4V   7231 Cac8I                              7269 RsaI
       ||          |              |                                       |
7201 agcaaatttactgaatgcaagtggctcagagcaggctcagaaaccagtgtccccttctggagaagcaagtacctctggacagcactcaagactaaaactg 7300
     S  K  F  T  E  C  K  W  L  R  A  G  S  E  T  S  V  P  F  W  R  S  K  Y  L  W  T  A  L  K  T  K  T  G
     >>>>>>>>>>>>>>>>>>>>>>>>>>>>>>>>>>>>>>>>>>>>>>>>>>>>>>>>>>>>>>>>>>>>>>>>>>>>>>>>>>>>>>>>>>>>>>>>>>>>
     1st part

        7310      7320      7330      7340      7350      7360      7370      7380      7390      7400
           *    *    *    *    *    *    *    *    *    *    *    *    *    *    *    *    *    *    *    *
                                                            7360 HpyCH4V
                                                           7359 NsiI              7382 AciI
                                     7338 Cac8I           7358 NlaIII        7377 Cac8I
                                     |                    |||                |    |
7301 gaactcaggaagaaggagactgaaagaaagatgtatagcctgcgagaaagaaagggtcatgcatacaaagaggtcagcgagccgcaggatgatgattacc 7400
       T  Q  E  E  G  D  *  K  K  D  V  *  P  A  R  K  K  G  S  C  I  Q  R  G  Q  R  A  A  G  *  *  L  P
     >>>>>>>>>>>>>>>>>>>>>>>>>>>>>>>>>>>>>>>>>>>>>>>>>>>>>>>>>>>>>>>>>>>>>>>>>>>>>>>>>>>>>>>>>>>>>>>>>>>>
     1st part

        7410      7420      7430      7440      7450      7460      7470      7480      7490      7500
           *    *    *    *    *    *    *    *    *    *    *    *    *    *    *    *    *    *    *    *
                                                     7452 HaeIII
                                                    7451 ApaI
                                   7436 AluI   7448 NlaIII                   7481 Hpy8I
                   7420 XmnI      7435 PvuII  7447 NcoI                  7477 HpyCH4V
                   |              ||          ||  ||                     |   |
7401 tctattgtgagatgtgtcagaacttcttcattgacagctgtgctgcccatgggcccccctacatttgtaaaggacagtgcagtggacaaggggcaccccaa 7500
      L  L  *  D  V  S  E  L  L  H  *  Q  L  C  C  P  W  A  P  Y  I  C  K  G  Q  C  S  G  Q  G  A  P  Q
     >>>>>>>>>>>>>>>>>>>>>>>>>>>>>>>>>>>>>>>>>>>>>>>>>>>>>>>>>>>>>>>>>>>>>>>>>>>>>>>>>>>>>>>>>>>>>>>>>>>>
     1st part

        7510      7520      7530      7540      7550      7560      7570      7580      7590      7600
           *    *    *    *    *    *    *    *    *    *    *    *    *    *    *    *    *    *    *    *
                                      7539 HaeIII                                         7595 AciI
                                 7534 MluCI                                         7589 DpnI
                                 |    |                                             |     |
7501 ccgttcagccctcagtctgcccccagggctgagaattgggccatcaggcatccctcaggctgggcttggagtatggaatgaggcatctgatctgccgctg 7600
     P  F  S  P  Q  S  A  P  R  A  E  N  W  A  I  R  H  P  S  G  W  A  W  S  M  E  *  G  I  *  S  A  A  G
     >>>>>>>>>>>>>>>>>>>>>>>>>>>>>>>>>>>>>>>>>>>>>>>>>>>>>>>>>>>>>>>>>>>>>>>>>>>>>>>>>>>>>>>>>>>>>>>>>>>>
     1st part

        7610      7620      7630      7640      7650      7660      7670      7680      7690      7700
           *    *    *    *    *    *    *    *    *    *    *    *    *    *    *    *    *    *    *    *
                7613 HaeIII      7630 MluCI
        7605 HpyCH4V       7625 HaeIII 7637 BbsI                          7675 DpnI
        |       |           |    |      |                                 |
7601 ggtctgcactttggcccttatgagggccgaattacagaagacgaagaggcagccaacaatggatactcctggctgatcaccaaggggagaaactgctatg 7700
       S  A  L  W  P  L  *  G  P  N  Y  R  R  R  R  G  S  Q  Q  W  I  L  L  A  D  H  Q  G  E  K  L  L  *
     >>>>>>>>>>>>>>>>>>>>>>>>>>>>>>>>>>>>>>>>>>>>>>>>>>>>>>>>>>>>>>>>>>>>>>>>>>>>>>>>>>>>>>>>>>>>>>>>>>>>
     1st part

        7710      7720      7730      7740      7750      7760      7770      7780      7790      7800
           *    *    *    *    *    *    *    *    *    *    *    *    *    *    *    *    *    *    *    *
                                                          7758 XmaI
                                                          7758 SmaI
                                                          7758 AvaI  7769 SapI            7795 RsaI
                             7729 HaeIII        7748 Hpy8I 7759 HpaII               7786 HaeIII
                             |                  |        ||         |               |        |
7701 agtatgtggatggaaaaGATAAATCCTGGGCCAACTGGatgaggtatgtgaactgtgcccgggatgatgaagagcagaacctggtggccttccagtacca 7800
      V  C  G  W  K  R  *  I  L  G  Q  L  D  E  V  C  E  L  C  P  G  *  *  R  A  E  P  G  G  L  P  V  P
     >>>>>>>>>>>>>>>>>>>>>>>>>>>>>>>>>>>>>>>>>>>>>>>>>>>>>>>>>>>>>>>>>>>>>>>>>>>>>>>>>>>>>>>>>>>>>>>>>>>>
     1st part

        7810      7820      7830      7840      7850      7860      7870      7880      7890      7900
           *    *    *    *    *    *    *    *    *    *    *    *    *    *    *    *    *    *    *    *
         7807 DpnI
        7806 BstYI                           7845 Hpy8I
        7806 BglII                  7836 HaeIII                            7877 HaeIII
```

```
      ||                            |        |                             |
7801 caggcagatcttctatagaacctgccgagtcattaggccaggctgtgaactgctggtctggtatggggatgaatacggccaggaactgggcatcaagtgg 7900
     Q  A  D  L  L  *  N  L  P  S  H  *  A  R  L  *  T  A  G  L  V  W  G  *  I  R  P  G  T  G  H  Q  V  G
     >>>>>>>>>>>>>>>>>>>>>>>>>>>>>>>>>>>>>>>>>>>>>>>>>>>>>>>>>>>>>>>>>>>>>>>>>>>>>>>>>>>>>>>>>>>>>>>>>>>>
     1st part

              7910      7920      7930      7940      7950      7960      7970      7980      7990      8000
         *    *    *    *    *    *    *    *    *    *    *    *    *    *    *    *    *    *    *    *
                        7924 NlaIII                        7959 NlaIII
                    7920 AluI                      7951 DpnI                  7978 HaeIII       7995 MluCI
                   7919 SacI                      7950 BstYI        7968 NlaIII                7994 ApoI
                   ||   |                         ||         |        |         |                ||
7901 ggcagcaagtggaagaaagagctcATGGCAGGGAGAGAACCAAAGCCAGAGATCCATCCATGTCCCTCATGCTGTCTGGCCTTTTCAAGTCAGAAATTTC 8000
       Q  Q  V  E  E  R  A  H  G  R  E  R  T  K  A  R  D  P  S  M  S  L  M  L  S  G  L  F  K  S  E  I  S
     >>>>>>>>>>>>>>>>>>>>>>>>>>>>>>>>>>>>>>>>>>>>>>>>>>>>>>>>>>>>>>>>>>>>>>>>>>>>>>>>>>>>>>>>>>>>>>>>>>>>
     1st part          SacI  2nd part
                                         >>>
                                         Start-Exon10

              8010      8020      8030      8040      8050      8060      8070      8080      8090      8100
         *    *    *    *    *    *    *    *    *    *    *    *    *    *    *    *    *    *    *    *
               8009 NlaIII
              8008 PciI
              8008 NspI
          8004 Hpy8I
          8004 HincII                              8053 HpyCH4V                                8093 DpnI
          |   ||                                   |                                           |
8001 TCAGTCAACATGTAGAACGCAATCACTCCTCTCAGAACTTCCCAGGACCATCTGCAAGAAAACTCCTCCAACCAGAGAATCCCTGCCCAGGGGATCAGAA 8100
      Q  S  T  C  R  T  Q  S  L  L  S  E  L  P  R  T  I  C  K  K  T  P  P  T  R  E  S  L  P  R  G  S  E
     >>>>>>>>>>>>>>>>>>>>>>>>>>>>>>>>>>>>>>>>>>>>>>>>>>>>>>>>>>>>>>>>>>>>>>>>>>>>>>>>>>>>>>>>>>>>>>>>>>>>
     2nd part

              8110      8120      8130      8140      8150      8160      8170      8180      8190      8200
         *    *    *    *    *    *    *    *    *    *    *    *    *    *    *    *    *    *    *    *
                    8120 DpnI
                   8119 BstYI                          8158 DpnI                           8193 NlaIII
                   ||                                      |                                   |
8101 TCAGGAGCAGCAATATCCAGATCCACACAGCCGTAATGACAAAACCAAAGGTCAAGAGATCAAAGAAAGGTCCAAACTCTTGAATAAAAGGACATGGCAG 8200
     S  G  A  A  I  S  R  S  T  Q  P  *  *  Q  N  Q  R  S  R  D  Q  R  K  V  Q  T  L  E  *  K  D  M  A  E
     >>>>>>>>>>>>>>>>>>>>>>>>>>>>>>>>>>>>>>>>>>>>>>>>>>>>>>>>>>>>>>>>>>>>>>>>>>>>>>>>>>>>>>>>>>>>>>>>>>>>
     2nd part

              8210      8220      8230      8240      8250      8260      8270      8280      8290      8300
         *    *    *    *    *    *    *    *    *    *    *    *    *    *    *    *    *    *    *    *
                       8223 BfaI
               8215 HaeIII      8232 XcmI        8249 AluI                                   8294 HaeIII
               |       |        |                |                                            |
8201 AGGGAGATTTCAAGGGCCTTTTCTAGCCCACCCAAAGGACAAATGGGGAGCTGTAGAGTGGGAAAAAGAATAATGGAAGAAGAGTCCAGAACAGGCCAGA 8300
       G  D  F  K  G  L  F  *  P  T  Q  R  T  N  G  E  L  *  S  G  K  K  N  N  G  R  R  V  Q  N  R  P  E
     >>>>>>>>>>>>>>>>>>>>>>>>>>>>>>>>>>>>>>>>>>>>>>>>>>>>>>>>>>>>>>>>>>>>>>>>>>>>>>>>>>>>>>>>>>>>>>>>>>>>
     2nd part

              8310      8320      8330      8340      8350      8360      8370      8380      8390      8400
         *    *    *    *    *    *    *    *    *    *    *    *    *    *    *    *    *    *    *    *
                                                      8356 HpyCH4V
                         8325 MluCI                8353 MluCI                 8379 Hpy8I        8397 MseI
                         |                         |  |                       |                 |
8301 AAGTGAATCCAGGGAACACAGgcaaattatttgtgggggtaggAATCTCAAGAATTGCAAAAGTCAAGTATGGAGAGTGTGGACAAGGTTTCAGTGTTAA 8400
      S  E  S  R  E  H  R  Q  I  I  C  G  G  R  N  L  K  N  C  K  S  Q  V  W  R  V  W  T  R  F  Q  C  *
     >>>>>>>>>>>>>>>>>>>>>>>>>>>>>>>>>>>>>>>>>>>>>>>>>>>>>>>>>>>>>>>>>>>>>>>>>>>>>>>>>>>>>>>>>>>>>>>>>>>>
     2nd part

              8410      8420      8430      8440      8450      8460      8470      8480      8490      8500
         *    *    *    *    *    *    *    *    *    *    *    *    *    *    *    *    *    *    *    *
                                                   8453 HpyCH4V
                                              8448 HpyCH4IV
                                        8442 AluI 8452 PstI      8467 AciI 8477 AluI
                                        |     |   ||             |         |
8401 ATCAGATGTTATTACACACCAAAGGACACATACAGGGGGAGAAGCTCTACGTCTGCAGGGAGTGTGGGCGGGGCTTTAGCTGGAAGTCACACCTCCTCATT 8500
     I  R  C  Y  Y  T  P  K  D  T  Y  R  G  E  A  L  R  L  Q  G  V  W  A  G  L  *  L  E  V  T  P  P  H  S
     >>>>>>>>>>>>>>>>>>>>>>>>>>>>>>>>>>>>>>>>>>>>>>>>>>>>>>>>>>>>>>>>>>>>>>>>>>>>>>>>>>>>>>>>>>>>>>>>>>>>
     2nd part

              8510      8520      8530      8540      8550      8560      8570      8580      8590      8600
         *    *    *    *    *    *    *    *    *    *    *    *    *    *    *    *    *    *    *    *
                                    8537 HpyCH4V             8562 Cac8I
                                   8536 PstI      8551 AciI 8561 AluI
                                   ||             |         ||
8501 CACCAGAGGATACACACAGGGGAGAAGCCCTATGTCTGCAGGGAGTGTGGGCGGGGCTTTAGCTGGCAGTCAGTCCTCCTCACTCACCAGAGGACACACA 8600
       P  E  D  T  H  R  G  E  A  L  C  L  Q  G  V  W  A  G  L  *  L  A  V  S  P  P  H  S  P  E  D  T  H
     >>>>>>>>>>>>>>>>>>>>>>>>>>>>>>>>>>>>>>>>>>>>>>>>>>>>>>>>>>>>>>>>>>>>>>>>>>>>>>>>>>>>>>>>>>>>>>>>>>>>
     2nd part
```

```
           8610      8620      8630      8640      8650      8660      8670      8680      8690      8700
         *    *    *    *    *    *    *    *    *    *    *    *    *    *    *    *    *    *    *    *
                                                   8647 HpaII
                                                  8646 NgoMIV
                         8621 HpyCH4V             8646 NaeI
                        8620 PstI      8635 AciI  8646 Cac8I
                        ||             |          ||
8601 CAGGGGAGAAGCCCTATGTCTGCAGGGAGTGTGGGCGGGGCTTTAGCCGGCAGTCAGTCCTCCTCACTCACCAGAGGAGACACACAGGGGAGAAGCCCTA 8700
      R  G  E  A  L  C  L  Q  G  V  W  A  G  L  *  P  A  V  S  P  P  H  S  P  E  E  T  H  R  G  E  A  L
     >>>>>>>>>>>>>>>>>>>>>>>>>>>>>>>>>>>>>>>>>>>>>>>>>>>>>>>>>>>>>>>>>>>>>>>>>>>>>>>>>>>>>>>>>>>>>>>>>>>>
     2nd part

           8710      8720      8730      8740      8750      8760      8770      8780      8790      8800
         *    *    *    *    *    *    *    *    *    *    *    *    *    *    *    *    *    *    *    *
                                   8731 HpaII
                                  8730 NgoMIV
         8705 HpyCH4V             8730 NaeI                                          8789 HpyCH4V
        8704 PstI      8719 AciI  8730 Cac8I                                        8788 PstI
        ||             |          ||                                                ||
8701 TGTCTGCAGGGAGTGTGGGCGGGGCTTTAGCCGGCAGTCAGTCCTCCTCACTCACCAGAGGAGACACACAGGGGAGAAGCCCTATGTCTGCAGGGAGTGT 8800
     C  L  Q  G  V  W  A  G  L  *  P  A  V  S  P  P  H  S  P  E  E  T  H  R  G  E  A  L  C  L  Q  G  V  W
     >>>>>>>>>>>>>>>>>>>>>>>>>>>>>>>>>>>>>>>>>>>>>>>>>>>>>>>>>>>>>>>>>>>>>>>>>>>>>>>>>>>>>>>>>>>>>>>>>>>>
     2nd part

           8810      8820      8830      8840      8850      8860      8870      8880      8890      8900
         *    *    *    *    *    *    *    *    *    *    *    *    *    *    *    *    *    *    *    *
                  8814 Cac8I                                                 8873 HpyCH4V             8898 Cac8I
       8803 AciI 8813 AluI                                                  8872 PstI      8887 AciI 8897 AluI
       |         ||                                                         ||             |         ||
8801 GGGCGGGGCTTTAGCTGGCAGTCAGTCCTCCTCAGTCACCAGAGGACACACACAGGGGAGAAGCCCTATGTCTGCAGGGAGTGTGGGCGGGGCTTTAGCT 8900
       A  G  L  *  L  A  V  S  P  P  Q  S  P  E  D  T  H  R  G  E  A  L  C  L  Q  G  V  W  A  G  L  *  L
     >>>>>>>>>>>>>>>>>>>>>>>>>>>>>>>>>>>>>>>>>>>>>>>>>>>>>>>>>>>>>>>>>>>>>>>>>>>>>>>>>>>>>>>>>>>>>>>>>>>>
     2nd part

           8910      8920      8930      8940      8950      8960      8970      8980      8990      9000
         *    *    *    *    *    *    *    *    *    *    *    *    *    *    *    *    *    *    *    *
                                                             8957 HpyCH4V
                                                            8956 PstI      8971 AciI
                                                            ||             |
8901 GGCAGTCAGTCCTCCTCACTCACCAGAGGACACACACAGGGGAGAAGCCCTATGTCTGCAGGGAGTGTGGGCGGGGCTTTAGCAATAAGTCACACCTCCT 9000
      A  V  S  P  P  H  S  P  E  D  T  H  R  G  E  A  L  C  L  Q  G  V  W  A  G  L  *  Q  *  V  T  P  P
     >>>>>>>>>>>>>>>>>>>>>>>>>>>>>>>>>>>>>>>>>>>>>>>>>>>>>>>>>>>>>>>>>>>>>>>>>>>>>>>>>>>>>>>>>>>>>>>>>>>>
     2nd part

           9010      9020      9030      9040      9050      9060      9070      9080      9090      9100
         *    *    *    *    *    *    *    *    *    *    *    *    *    *    *    *    *    *    *    *
                                             9041 HpyCH4V           9064 NruI
                                            9040 PstI      9055 AciI 9065 BstUI
                                            ||             |        ||
9001 CAGACACCAGAGGACACACACAGGGGAGAAGCCCTATGTCTGCAGGGAGTGTGGGCGGGGCTTTCGCGATAAGTCACACCTCCTCAGACACCAGAGGACA 9100
     Q  T  P  E  D  T  H  R  G  E  A  L  C  L  Q  G  V  W  A  G  L  S  R  *  V  T  P  P  Q  T  P  E  D  T
     >>>>>>>>>>>>>>>>>>>>>>>>>>>>>>>>>>>>>>>>>>>>>>>>>>>>>>>>>>>>>>>>>>>>>>>>>>>>>>>>>>>>>>>>>>>>>>>>>>>>
     2nd part

           9110      9120      9130      9140      9150      9160      9170      9180      9190      9200
         *    *    *    *    *    *    *    *    *    *    *    *    *    *    *    *    *    *    *    *
                             9125 HpyCH4V
                            9124 PstI      9139 AciI
                            ||             |
9101 CACACAGGGGAGAAGCCCTATGTCTGCAGGGAGTGTGGGCGGGGCTTTAGAGATAAGTCAAACCTCCTCAGTCACCAGAGGACACACACAGGGGAGAAGC 9200
       H  R  G  E  A  L  C  L  Q  G  V  W  A  G  L  *  R  *  V  K  P  P  Q  S  P  E  D  T  H  R  G  E  A
     >>>>>>>>>>>>>>>>>>>>>>>>>>>>>>>>>>>>>>>>>>>>>>>>>>>>>>>>>>>>>>>>>>>>>>>>>>>>>>>>>>>>>>>>>>>>>>>>>>>>
     2nd part

           9210      9220      9230      9240      9250      9260      9270      9280      9290      9300
         *    *    *    *    *    *    *    *    *    *    *    *    *    *    *    *    *    *    *    *
             9209 HpyCH4V                                                                       9293 HpyCH4V
            9208 PstI      9223 AciI                                                           9292 PstI
            ||             |                                                                   ||
9201 CCTATGTCTGCAGGGAGTGTGGGCGGGGCTTTAGCAATAAGTCACACCTCCTCAGACACCAGAGGACACACACAGGGGAGAAGCCCTATGTCTGCAGGGA 9300
      L  C  L  Q  G  V  W  A  G  L  *  Q  *  V  T  P  P  Q  T  P  E  D  T  H  R  G  E  A  L  C  L  Q  G
     >>>>>>>>>>>>>>>>>>>>>>>>>>>>>>>>>>>>>>>>>>>>>>>>>>>>>>>>>>>>>>>>>>>>>>>>>>>>>>>>>>>>>>>>>>>>>>>>>>>>
     2nd part

           9310      9320      9330      9340      9350      9360      9370      9380      9390      9400
         *    *    *    *    *    *    *    *    *    *    *    *    *    *    *    *    *    *    *    *
                                                                                 9377 HpyCH4V
           9307 AciI                                                            9376 PstI      9391 AciI
           |                                                                    ||             |
9301 GTGTGGGCGGGGCTTTCGCAATAAGTCACACCTCCTCAGACACCAGAGGACACACACAGGGGAGAAGCCCTATGTCTGCAGGGAGTGTGGGCGGGGCTTT 9400
     V  W  A  G  L  S  Q  *  V  T  P  P  Q  T  P  E  D  T  H  R  G  E  A  L  C  L  Q  G  V  W  A  G  L  *
```

```
     >>>>>>>>>>>>>>>>>>>>>>>>>>>>>>>>>>>>>>>>>>>>>>>>>>>>>>>>>>>>>>>>>>>>>>>>>>>>>>>>>>>>>>>>>>>>>>>>>>
     2nd part


            9410      9420      9430      9440      9450      9460      9470      9480      9490      9500
         *    *    *    *    *    *    *    *    *    *    *    *    *    *    *    *    *    *    *    *
                                                                   9461 HpyCH4V
                                                                  9460 PstI
                                                              9456 HpyCH4IV
                                                               |    ||
9401 AGCGATAGGTCAAGCCTCTGCTATCACCAGAGGACACACACAGGGGAGAAGCCCTACGTCTGCAGGGAGGATGAGTAAGTCATTAGTAATAAAACCTCAT 9500
       R  *  V  K  P  L  L  S  P  E  D  T  H  R  G  E  A  L  R  L  Q  G  G  *  V  S  H  *  *  *  N  L  I
     >>>>>>>>>>>>>>>>>>>>>>>>>>>>>>>>>>>>>>>>>>>>>>>>>>>>>>>>>>>>>>>>>>>>>>>>>>>>>>>>>>>>>>>>>>>>>>>>>>
     2nd part                                                                      hPrdm9-3'UTR


            9510      9520      9530      9540      9550      9560      9570      9580      9590      9600
         *    *    *    *    *    *    *    *    *    *    *    *    *    *    *    *    *    *    *    *
                                                          9547 BfaI
                                                         9546 XbaI  9557 MseI
              9509 AciI                            9541 AluI      9556 DraI                             9599 Cac8I
      9502 TaqI                                   9540 HindIII   9555 PmeI                   9587 MseI  9598 HpyCH4V
      |      |                                     ||     ||        |||                              |          ||
9501 CTCGATAGCCGCAAAAAGACAAATGTGGTCACCACACACAAGCTTTCTAGACCAGTTTAAACATCACCATCACCATCACTAAGTGATTAACCTCAGGTGC 9600
      S  I  A  A  K  R  Q  M  W  S  P  H  T  S  F  L  D  Q  F  K  H  H  H  H  H  H  *  V  I  N  L  R  C
     >>>>>>>>>>>>>>>>>>>>>>>>>>>>>>>>>>>>>>>>>>>>>>>>>>>>
     hPrdm9-3'UTR                                  HindIII


            9610      9620      9630      9640      9650      9660      9670      9680      9690      9700
         *    *    *    *    *    *    *    *    *    *    *    *    *    *    *    *    *    *    *    *
                                               9632 HaeIII                          9667 DpnI
                                              9631 MscI                          9663 DpnI                9688 MluCI
                                              ||                                  |   |                    |
9601 AGGCTGCCTATCAGAAGGtggtggctggtgtggccaatgcCCTGGCTCACAAATACCACTGAgatcgatctttttccctctgccaaaaattatggggaca 9700
     R  L  P  I  R  R  W  W  L  V  W  P  M  P  W  L  T  N  T  T  E  I  D  L  F  P  S  A  K  N  Y  G  D  I


            9710      9720      9730      9740      9750      9760      9770      9780      9790      9800
         *    *    *    *    *    *    *    *    *    *    *    *    *    *    *    *    *    *    *    *
      9702 NlaIII                                 9743 MluCI                     9772 MluCI
     9701 BspHI                                  9742 ApoI     9756 HpyCH4V   9771 ApoI                     9800 NdeI
     ||                                           ||             |              ||                           |
9701 tcatgaagccccttgagcatctgacttctggctaataaaggaaatttattttcattgcaatagtgtgttggaatttttttgtgtctctcactcggaaggac 9800
       M  K  P  L  E  H  L  T  S  G  *  *  R  K  F  I  F  I  A  I  V  C  W  N  F  L  C  L  S  L  G  R  T


            9810      9820      9830      9840      9850      9860      9870      9880      9890
         *    *    *    *    *    *    *    *    *    *    *    *    *    *    *    *    *
                         9820 MseI                                      9868 NdeI
                        9819 DraI                                9860 NdeI       9877 BfaI
                        ||                                        |       |         |
9801 atatgggagggcaaatcatttaaaacatcagaatgagtatttggtttagagtttggcaacatatgcccatatgtaactagcataacccccttgg 9893
      Y  G  R  A  N  H  L  K  H  Q  N  E  Y  L  V  *  S  L  A  T  Y  A  H  M  *  L  A  *  P  L
```
